# Decoding the Molecular Language of Proteins with Evolla

**DOI:** 10.1101/2025.01.05.630192

**Authors:** Xibin Zhou, Chenchen Han, Yingqi Zhang, Huan Du, Jiayuan Tian, Jin Su, Renju Liu, Kai Zhuang, Shiyu Jiang, Anthony Gitter, Li Liu, Huayu Li, Meiqi Wu, Shiyang You, Zichen Yuan, Feng Ju, Huilin Zhang, Wei Zheng, Fengyuan Dai, Yuyang Zhou, Yuyang Tao, Dan Wu, Zongze Shao, Yang Liu, Hongyuan Lu, Fajie Yuan

## Abstract

Proteins, nature’s intricate molecular machines, are the products of billions of years of evolution and play fundamental roles in sustaining life. Yet, deciphering their molecular language—understanding how sequences and structures encode biological functions—remains a cornerstone challenge. Here, we introduce Evolla, an interactive protein-language model designed to transcend static classification by interpreting protein function through natural language queries. Trained on 546 million protein–text pairs and refined via Direct Preference Optimization, Evolla couples high-dimensional molecular representations with generative semantic decoding. Benchmarking establishes Evolla’s superiority over general large language models in functional inference, demonstrates zero-shot performance parity with the state-of-the-art supervised model, and exposes remote functional relationships invisible to conventional alignment. We validate Evolla through two distinct applications: identifying candidate eukaryotic signature proteins in Asgard archaea, with functional Vps4 homologs validated via yeast complementation; and interactively discovering a novel deep-sea polyethylene terephthalate (PET) hydrolase, *Ps*PETase, confirmed to degrade plastic films. These results position Evolla not merely as a predictor, but as a generative engine capable of complex hypothesis formulation, shifting the paradigm from static annotation to interactive, actionable discovery. The Evolla online service is available at http://www.chat-protein.com/.

---

Elucidating protein function is fundamental to understanding cellular mechanisms and advancing biotechnology. While biological functions are encoded in sequences and manifested through structures [1], decoding this information remains a formidable challenge. Recent breakthroughs in structure prediction, exemplified by AlphaFold [2, 3], have revolutionized structural biology but paradoxically highlighted the widening gap between structural determination and functional understanding. With billions of sequences now available yet fewer than one million proteins possessing expert-curated annotations [4], the scarcity of functional insights has become a critical bottleneck in biological research.

Current annotation methodologies face significant challenges in bridging this gap. Traditional alignment-based tools, such as BLAST [5], DIAMOND [6], and even structural aligners like Foldseek [7], often falter when applied to proteins lacking clear evolutionary relatives or structural analogs. Conversely, discriminative deep learning classifiers [8–24] have achieved success without explicit alignment but are typically restricted to narrow, predefined output spaces—such as models trained exclusively on specific protein families. While these models can effectively categorize proteins, they often lack the capacity to elucidate the molecular context or provide mechanistic rationales underlying their predictions. More recently, multimodal protein-language models have sought to address these limitations through generative approaches [25–28]. However, frequently constrained by limited scale or shallow cross-modal alignment, these models often struggle to ground their outputs in biological reality, thereby limiting their utility for reliable hypothesis generation. This underscores the need for a critical evolution: advancing from shallow cross-modal association to achieve deep, context-aware interpretation that truly synthesizes biological knowledge.

Here, we introduce Evolla, a large-scale protein-language generative model with architectures spanning from 10 to 80 billion parameters, designed to interpret protein function through natural language queries. Evolla integrates molecular modalities with linguistic context, trained on a massive corpus of 546 million protein–text pairs that synergizes manually curated annotations with high-confidence computational descriptions, and refined via Direct Preference Optimization (DPO) [29]. This framework enables Evolla to transcend static annotation, serving as a dynamic partner capable of processing complex and open-ended biological inquiries. Comprehensive evaluations against general-purpose large language models (LLMs) and specialized protein models demonstrate Evolla’s superior ability to generate precise, nuanced, and contextually relevant functional descriptions, as well as its robustness in zero-shot classification and the identification of remote functional similarity.

We demonstrate Evolla’s capacity to facilitate hypothesis-driven discovery through two distinct applications. First, we explored the deep evolutionary transition from prokaryotes to eukaryotes by analyzing Asgard archaea. Rather than relying solely on rigid alignment thresholds, we employed Evolla to perform high-throughput semantic screening, specifically interrogating proteins for eukaryotic signatures such as the Microtubule Interacting and Transport (MIT) domain. This semantic approach successfully identified functional Vps4 homologs, whose ability to remodel endosomal membranes was subsequently confirmed via heterologous complementation assays in *Saccharomyces cerevisiae*. Second, addressing the environmental crisis of plastic pollution, we utilized Evolla to guide the discovery of a novel PET hydrolase from a deep-sea *Pseudomonas* isolate. In this workflow, Evolla first pinpointed a high-confidence candidate, *Ps*PETase, via semantic filtering, and subsequently transcended simple classification by assisting in experimental design—reasoning over biophysical constraints such as secretory nature and thermal adaptation. This candidate was experimentally validated to degrade PET films. These results illustrate Evolla’s potential not merely as an annotation tool, but as a generative engine that bridges the gap between static molecular data and actionable biological insights.

## 1 Evolla’s Architecture and Training

Evolla’s architecture is conceived as a generative system to bridge the distinct modalities of protein and human language (see Figure 1a). Its design philosophy posits that a profound functional understanding of proteins emerges from the synergy between a model adept at interpreting molecular data and another skilled in complex reasoning and linguistic expression. This modular architecture comprises three core components: a protein encoder, a language decoder, and a novel interfacing module that connects them, enabling efficient and scalable training.

**Fig. 1.**
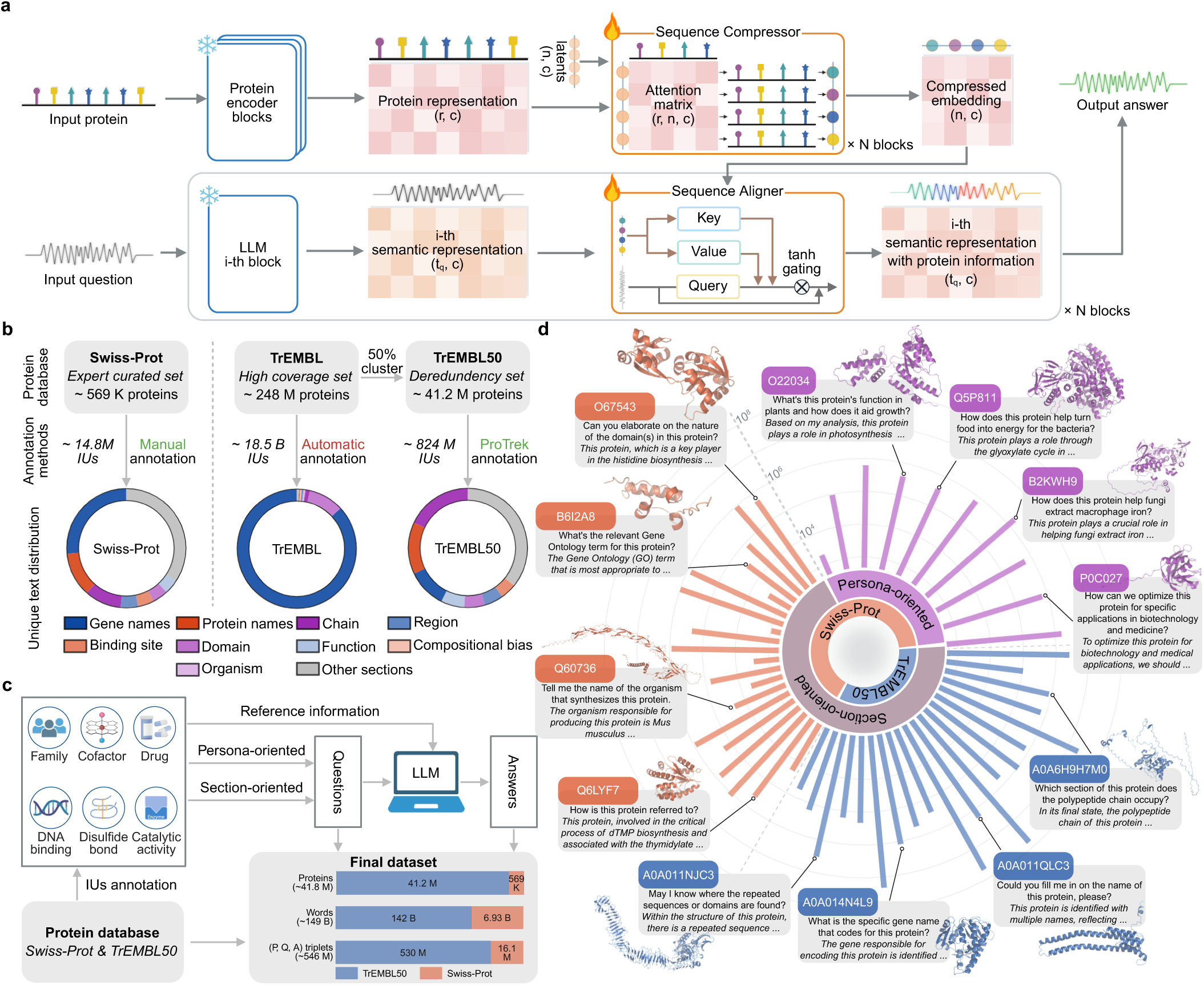
Architecture of the Evolla framework and the construction of its instruction-tuning dataset. **(a)** The Evolla architecture. A frozen protein encoder generates residue-level representations, which are distilled into fixed-length embeddings by a trainable Sequence Compressor. These compressed embeddings are injected into a frozen LLM (Llama3) via trainable, interleaved Sequence Aligner blocks equipped with a tanh gating mechanism to synthesize textual responses. **(b)** Curation of protein Information Units (IUs). The dataset integrates expert-reviewed Swiss-Prot entries (569K proteins, manual annotation) and a dereplicated TrEMBL50 set (41.2M proteins, clustered at 50% sequence identity). TrEMBL50 entries were re-annotated using ProTrek to yield 824M IUs. Donut charts visualize the distribution of unique text categories (e.g., gene names, functional domains) across datasets. **(c)** Dataset construction pipeline. IUs derived from biological databases serve as reference knowledge. An LLM generates question-answer pairs using two distinct strategies: persona-oriented (simulating diverse user roles) and section-oriented (targeting specific protein attributes). The final corpus comprises approximately 546 million protein-question-answer triplets, covering 41.8 million proteins and 150 billion tokens. **(d)** Prompt diversity and distribution. The sunburst chart illustrates the hierarchical distribution of instruction types—categorized by persona and section—for both Swiss-Prot and TrEMBL50 subsets. Outer callouts display representative Q&A examples for specific proteins.

### 1.1 Evolla Architecture for Protein-Text Alignment

The architecture of Evolla begins with a pre-trained protein language model (PLM) as the encoder. We selected SaProt [14, 23] as the encoder backbone after a rigorous comparative benchmark against other leading PLMs, including ESM2 [30] and MSA-Transformer [31], due to its enhanced performance in capturing comprehensive protein representations (Methods 7.1.1 and Extended Fig. S1a). The PLM’s primary role is to distill the high-dimensional information from protein sequences—and structures where applicable—into a compact, semantically rich latent representation. For the decoder, we use Llama3 [32] (specifically the 8B and 70B variants), a powerful open-weight LLM, which serves as the reasoning and articulation engine (Methods 7.1.4). During training, both the PLM and LLM backbones remain frozen. This strategy significantly enhances training efficiency, allowing computational resources to be focused on the critical task of aligning these two powerful but disparate modalities.

Evolla is implemented at two scales—a 10B-parameter version and a frontier 80B-parameter model—establishing it as one of the largest generative models in the biological domain. Each version incorporates PLM and LLM backbones scaled to match model capacity (Methods 7.4). Notably, while the SaProt-650M backbone in the 10B version is limited to structure-aware (SA) token sequences, the SaProt-1.3B backbone in the 80B version supports both SA tokens and standard amino acid sequences. Consequently, Evolla-80B offers the flexibility to process either structural or pure sequence inputs. A pivotal component of our design is the interfacing module that connects these backbones. Comprising two neural network blocks—the Sequence Compressor and the Sequence Aligner—this module bridges the fundamental gap between the numerical protein representations generated by the encoder and the linguistic inference process of the decoder.

The Sequence Compressor, a Transformer-based module shown to be critical for model performance (see Extended Fig. S1b), is designed to distill essential functional information from variable-length protein representations into a fixed-length ‘protein functional codebook’ (Methods 7.1.2 and Extended Fig. S2a). This is achieved through a Flamingo-style [33] cross-attention mechanism where a small set of learnable latent tokens act as queries. These queries attend to the full protein representation from the PLM encoder, learning to identify and aggregate conserved functional motifs. This process effectively condenses complex molecular information into a standardized, information-dense codebook, forming the basis for all subsequent reasoning.

The Sequence Aligner then integrates this protein codebook into the LLM’s reasoning process through an architecture that inserts the aligner module directly between the layers of the frozen LLM (Methods 7.1.3 and Extended Fig. S2b). This allows it to modulate the flow of information within the LLM’s internal pathways. The Aligner itself is a gated cross-attention block [33, 34], where the LLM’s hidden states query the protein codebook. A learnable gating mechanism dynamically controls the strength of the protein-specific signal injected at each layer. This design ensures that the protein context provides a subtle yet targeted guidance to the LLM’s generation process, rather than overwhelming its vast pre-existing knowledge.

### 1.2 Training Evolla on a Massive Protein-Text Dataset

Evolla is trained to minimize a causal protein-language modeling (CPLM) objective. For each training instance (*p, q, t*) comprising a protein *p*, a question *q*, and an answer *t* from the training dataset *D*, the model, parameterized by Θ, is trained to predict the next token *t_k_* in the answer, conditioned on the preceding text *t_<k_*, the protein *p* and question *q* context:

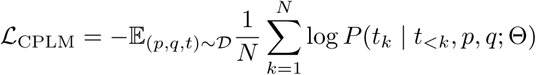

This objective forces the model to establish a robust mapping between the protein’s molecular features and the linguistic concepts required to answer complex questions.

A critical bottleneck in protein-language modeling is the scarcity of large-scale, high-quality protein-text data. To address this, we developed an automated data curation pipeline designed to synthesize a training corpus of extensive breadth and depth (Methods 7.5). Our strategy centered on aggregating Information Units (IUs)—concise, attribute-specific textual annotations—into a comprehensive knowledge repository (Fig. 1b). We integrated high-confidence, expert-reviewed annotations from Swiss-Prot with IUs derived specifically for a non-redundant subset of TrEMBL sequences. To mitigate the skewed annotation distribution inherent in TrEMBL, we constructed a non-redundant TrEMBL50 dataset via sequence clustering and employed the cross-modal model ProTrek [22] to retrieve a diverse set of IUs. This repository served as the factual substrate for generating a large-scale corpus of protein-question-answer triplets (Fig. 1c). Leveraging an LLM, we first generated diverse questions simulating scientific inquiry, conditioned on the IUs for a given protein (Methods 7.5.3). Subsequently, we synthesized detailed answers grounded in these IUs (Methods 7.5.4). This synthesis step proved essential; directly utilizing raw IUs as targets yielded dismal performance and truncated outputs (Extended Fig. S3). The resulting corpus comprises 546 million triplets totaling over 150 billion tokens (Fig. 1c). This scale surpasses that of recent studies by two to three orders of magnitude [25, 26], providing the volume and diversity necessary to capture the complexity of the protein universe (Fig. 1d).

Scaling both data volume and model capacity was instrumental in enhancing performance, as quantified by GPT score (Fig. 2d, blue trajectory; Methods 7.6, 7.7). Our experiments demonstrated consistent improvements in accuracy as we scaled the training data from 1% (Medium) to 10% (Large) of the corpus, and finally to the full 546 million samples (Total). Similarly, scaling from 10B to 80B parameters yielded pronounced gains on the hard, low-homology test sets (sequence identity *<* 30%)(Methods 7.6), suggesting that sufficient model capacity is crucial for robust generalization in this complex task.

**Fig. 2.**
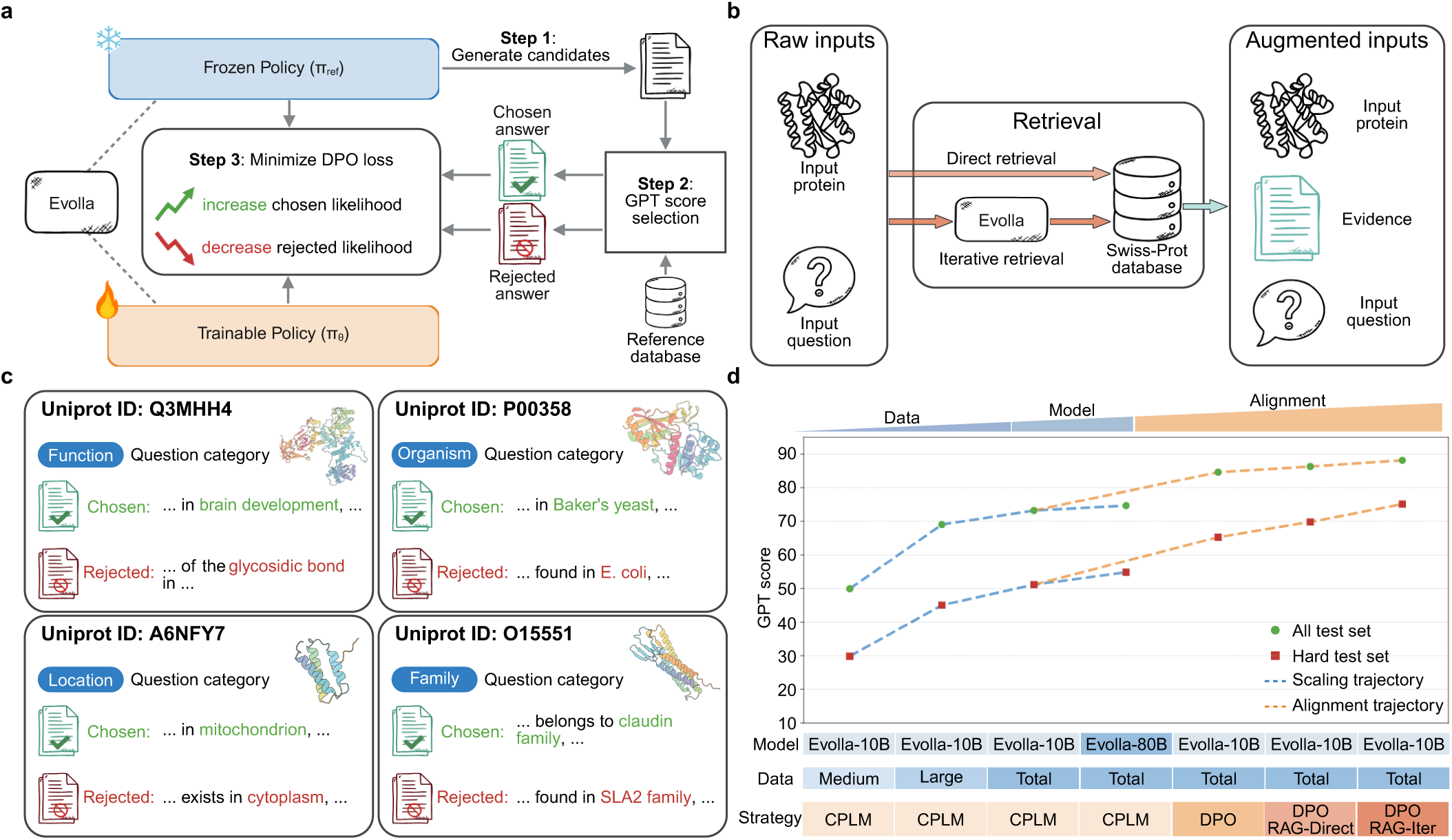
Optimization and alignment strategies for Evolla. **(a)** Schematic of the Direct Preference Optimization (DPO) workflow. Evolla is duplicated into a frozen reference policy (*π*_ref_) and a trainable policy (*π_θ_*). The frozen policy generates candidate answers (Step 1), which are evaluated against a reference database using GPT score (Step 2) to identify chosen (correct) and rejected (incorrect) responses. These pairs are used to minimize the DPO loss (Step 3), optimizing the model to prefer factual outputs. **(b)** Retrieval-Augmented Generation (RAG) framework. Two retrieval mechanisms are implemented: *Direct retrieval* queries the Swiss-Prot database using the raw input, while *Iterative retrieval* utilizes an initial hypothesis generated by Evolla to refine the search. The retrieved evidence is concatenated with the protein and question to form augmented inputs for the final generation. **(c)** Representative training instances from the DPO dataset. Examples across different categories (Function, Organism, Location, Family) show the construction of preference pairs, where factually accurate answers are labeled as ‘chosen’ (green) and hallucinations or inaccuracies as ‘rejected’ (red). **(d)** Comparative analysis of scaling versus alignment strategies. The chart tracks GPT scores across two development trajectories: the *scaling trajectory* (blue dashed lines) increases data volume (Medium: 1% of Total; Large: 10% of Total) or model size (10B to 80B) via causal protein-language modeling (CPLM); the *alignment trajectory* (orange dashed lines) applies DPO and RAG (Direct and Iterative) to the fixed 10B model. Performance is reported on the full test set (green circles) and a hard subset (red squares).

## 2 Aligning Evolla with Functional Veracity

Although pre-training on massive corpora equips Evolla with an expansive biological knowledge base, ensuring the factual validity of its generative outputs remains a critical challenge. To mitigate the risk of “hallucination”—the generation of plausible but functionally incorrect hypotheses [35]—we implemented a dual-component alignment framework (Fig. 2a, b). This approach first calibrates the model’s internal representations through preference optimization and subsequently grounds its generation in external evidence.

To steer the model’s generative process towards scientifically validated statements, we employed Direct Preference Optimization (DPO) [29] (Fig. 2a and Methods 7.8). We fine-tuned the base model using a curated dataset, D_pref_, comprising 37,544 preference quadruplets (*p, q, t_c_, t_r_*). For each protein-question pair (*p, q*), we generated multiple candidate responses and utilized GPT score as a scalable proxy for expert assessment [36] to distinguish high-veracity (chosen, *t_c_*) from low-veracity (rejected, *t_r_*) answers (Methods 7.7 and Fig. 2c). The model was optimized via the DPO objective:

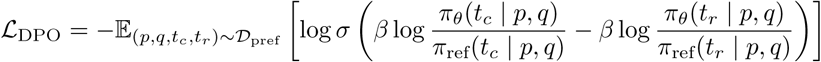

This objective optimizes the policy *π_θ_* to increase the likelihood of the preferred response *t_c_* relative to *t_r_*, constrained by the reference policy *π*_ref_. By tuning the hyperparameter *β*, this procedure effectively suppresses generation patterns associated with factual inaccuracies, refining the model’s internal knowledge alignment.

To complement this internal calibration, we implemented a Retrieval Augmented Generation (RAG) [37] strategy, enabling Evolla to consult external databases prior to response formulation (Methods 7.9). The framework operates via two distinct retrieval modes (Fig. 2b and Extended Fig. S4). The first, direct retrieval, serves as an efficient heuristic, rapidly linking queries to thematically relevant data from related proteins (Extended Fig. S4a). The second, iterative retrieval (or rethinking retrieval), emulates a refinement process wherein the model formulates a preliminary hypothesis and actively seeks textual evidence to validate or adjust its reasoning (Extended Fig. S4b). In both modes, retrieved information is synthesized with the original query to construct an evidence-rich prompt (Extended Fig. S4c). This strategy ensures that generative outputs are a grounded synthesis of parametric knowledge and established biological fact.

Enhanced by the dual alignment strategies, Evolla achieves substantially improved performance compared to its vanilla version, particularly on the hard test set (Methods 7.6, Fig. 2d, orange trajectory). These results also underscore a key principle: while large-scale model training establishes a robust foundation (section 1.2), unlocking the potential for factually accurate, expert-level insights requires a sophisticated alignment strategy that synergizes internal knowledge calibration (DPO) with external evidence grounding (RAG).

## 3 Generative Understanding of Protein Functions

To assess Evolla’s comprehensive capabilities in protein function interpretation, we benchmarked the 10B-DPO version across a spectrum of downstream tasks against a diverse array of representative models. Crucially, all evaluations in this section were conducted without external RAG to strictly isolate the model’s intrinsic parametric knowledge. We first benchmarked it against mainstream general-domain large language models (LLMs) on the full test set (Methods 7.6). Since these LLMs occasionally refuse to process raw amino acid sequences, we engineered specific prompts to enable comparison (Methods 7.10; Supplementary Table S8). In this benchmark, Evolla substantially outperformed two widely used models, GPT-4o [38] and DeepSeek-V3 [39], in generating accurate functional descriptions, as measured by the GPT score (Fig. 3a; Methods 7.7). These results highlight the necessity of domain-specific training for specialized biological tasks.

**Fig. 3.**
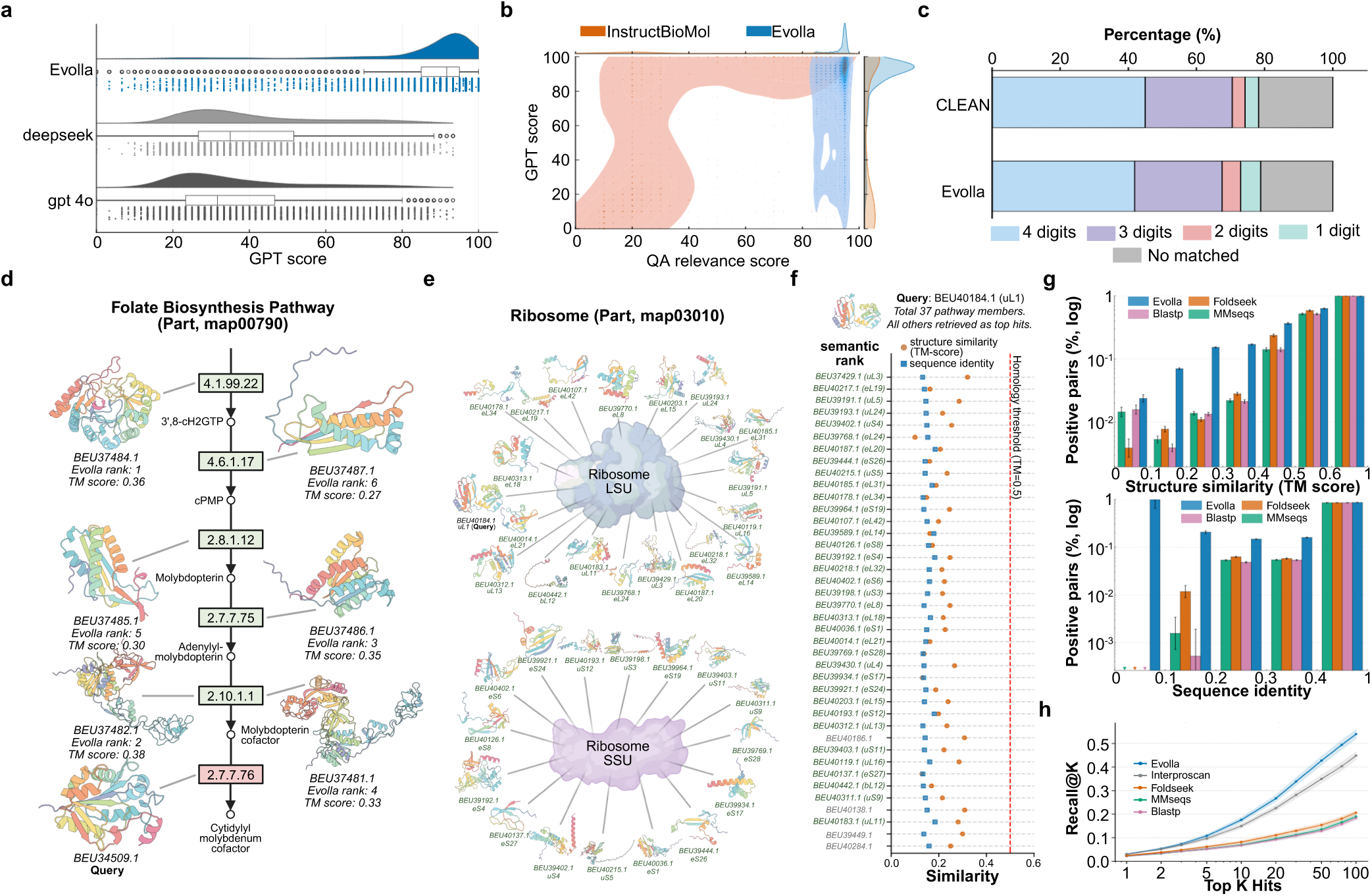
Comprehensive benchmarking of Evolla against general-purpose LLMs and specialized bioinformatics tools. **(a)** Performance against LLMs. Raincloud plots display the density and individual data points of GPT scores for protein descriptions generated by Evolla, DeepSeek, and GPT-4o. Evolla exhibits a distribution concentrated at the higher end of the score spectrum compared to the broader variance observed in general LLMs. **(b)** Performance against InstructBioMol. A 2D density estimation of GPT score (factual accuracy) versus QA relevance score. Evolla (blue) data points cluster in the upper-right quadrant, indicating simultaneous high accuracy and relevance, whereas InstructBioMol (orange) shows a dispersed distribution extending into lower accuracy or relevance regions. **(c)** Zero-shot EC number classification. Bar chart comparing the accuracy of Evolla (zero-shot) against the supervised specialist model CLEAN on a temporal hold-out set. **(d)** Semantic retrieval of metabolic partners. A query using a single enzyme (red) from the Folate Biosynthesis pathway retrieves upstream metabolic partners (green). Annotated TM-scores (0.27–0.38) indicate that retrieval occurs independently of significant structural similarity. **(e, f)** Reconstruction of the Ribosome complex. **(e)** Network visualization of the archaeal ribosome subunits retrieved using a single subunit uL1 query. **(f)** Rank-ordered retrieval results showing that 36 subunits are identified within the top 40 hits. The retrieved hits exhibit low sequence identity (blue squares) and structural similarity (orange circles), with structural similarity consistently below the empirical structural homology threshold (red dashed line, TM-score = 0.5). **(g)** Retrieval performance stratified by similarity. The bar charts show the fraction of positive pairs (sharing GO terms) retrieved by different methods, binned by structural (top) and sequence (bottom) similarity. Error bars represent the 95% confidence interval of the mean. Evolla demonstrates higher retrieval rates in the low-similarity bins (*<* 0.3 TM-score or sequence identity) compared to homology-based tools (Blastp, Foldseek, MMseqs). **(h)** Pathway discovery in a novel archaeon. Recall@K curves for pathway co-member retrieval. Evolla (blue line) shows higher recall across top-K hits compared to sequence-based and structure-based baselines. Shaded areas represent 95% confidence intervals.

We next benchmarked Evolla against two recent multimodal protein-language models on the same test set: ChatNT [26] and InstructBioMol [40] (Supplementary Table S9; Methods 7.11). As the nucleotide-centric ChatNT failed to process protein-related queries (Supplementary Table S9), our analysis focused on InstructBioMol. Evaluations based on factual accuracy (GPT score) and QA relevance (Methods 7.7) revealed significant limitations in InstructBioMol (Fig. 3b). While occasionally factually correct, InstructBioMol struggled to provide relevant answers, with most outputs scoring poorly on relevance. Furthermore, its responses frequently exhibited signs of memorization, reproducing training data excerpts rather than generating novel insights (Extended Fig. S5a). In contrast, Evolla consistently excelled in both accuracy and relevance, offering generative solutions that transcend simple memorization.

To evaluate its zero-shot generalization ability, we assessed Evolla on Enzyme Commission (EC) number assignment using a temporal hold-out set (Methods 7.12). Given Evolla’s generative nature, we developed an Instructional Response Matching (IRM) framework to map its outputs to EC predictions (Methods 7.13.2 and 7.13.1), benchmarking performance against CLEAN [41], a state-of-the-art supervised classifier specialized for this domain. Remarkably, despite lacking any task-specific fine-tuning, Evolla achieved an accuracy of 41.85% at the 4-digit level. This performance is competitive with the 44.98% accuracy of CLEAN, a fully supervised specialist (Fig. 3c). Furthermore, Evolla demonstrated robust performance on partial (1–3 digit) matches, matching or surpassing CLEAN. These results are particularly significant considering the fundamental difference in model scope: while CLEAN is purpose-built solely for this specific classification task, EC number prediction represents just one facet of Evolla’s broad, multi-task capabilities.

Leveraging its generative descriptions and the IRM framework (Methods 7.13.3), Evolla enables semantic protein search, bypassing conventional sequence or structural alignment. This approach is uniquely suited for identifying remote functional analogs—proteins that share biological functions despite divergent sequences and structures. To validate this, we benchmarked retrieval performance using shared Gene Ontology (GO) terms as the ground truth for functional relatedness (Methods 7.13.4). When compared to classical tools (BLASTp [42], MMseqs2 [43], and Foldseek [7]), Evolla performed comparably in high-similarity regimes but demonstrated a decisive advantage in the low-homology regime (TM-score [44] *<* 0.4 or sequence identity *<* 0.4). In this challenging zone, Evolla retrieves functionally related pairs with nearly an order of magnitude higher sensitivity (Fig. 3g), effectively uncovering functional relationships that remain invisible to sequence- or structure-based alignment methods.

We next evaluated whether Evolla could generalize from identifying protein pairs to reconstructing entire biological pathways. Using the proteome of the Asgard archaeon HC1 [45] and KEGG pathway co-membership as ground truth (Methods 7.13.5), we benchmarked Evolla against standard homology-based tools and the functional annotation tool InterProScan [46]. Evolla notably outperformed all baselines, recovering nearly half of true pathway members within its top 75 hits (Fig. 3h). This performance is consistent across diverse pathway types, including metabolic cascades and macromolecular assemblies. For instance, in the Folate Biosynthesis pathway (map00790), a single enzyme query (BEU34509.1) successfully retrieved direct upstream partners despite minimal structural similarity (Fig. 3d). Even more strikingly, for the Ribosome (map03010), querying a single subunit (uL1, BEU40184.1) retrieved the complete set of 36 remaining subunits within the top 40 hits (Fig. 3e,f). These results underscore Evolla’s capacity to reconstruct complex biological systems and capture deep functional logic where traditional homology search fails.

## 4 Inference of Eukaryotic Complexity in Asgard Archaea by Chatting with Evolla

The evolutionary transition from prokaryotes to eukaryotes represents a fundamental biological enigma [47]. While the discovery of Asgard archaea and their Eukaryotic Signature Proteins (ESPs) has bridged this gap, identifying these proteins typically necessitates laborious phylogenetic reconstructions and complex homology searches [48–50]. Here, we demonstrate how Evolla overcomes these traditional limitations by effectively distilling deep evolutionary signals from a vast proteomic background through high-dimensional semantic reasoning.

Our investigation utilized a comprehensive dataset comprising proteomes predicted from 14 assembled Asgard archaeal genomes, totaling 48,128 protein sequences (Methods 7.15). We deployed Evolla for high-throughput semantic screening, probing each protein with the hypothesis-driven query: ‘Is this protein potentially a eukaryotic signature protein?’ (Fig. 4a). To rigorously filter the outputs, we implemented a secondary LLM-based evaluation framework that stratified Evolla’s semantic responses into five confidence levels, ranging from ‘confidently yes’ to ‘confidently no’ (Methods 7.15). By retaining only proteins in the highest confidence tier (‘confidently yes’), we distilled the initial dataset into 1,855 preliminary ESP candidates (Supplementary File 4, Table 2).

**Fig. 4.**
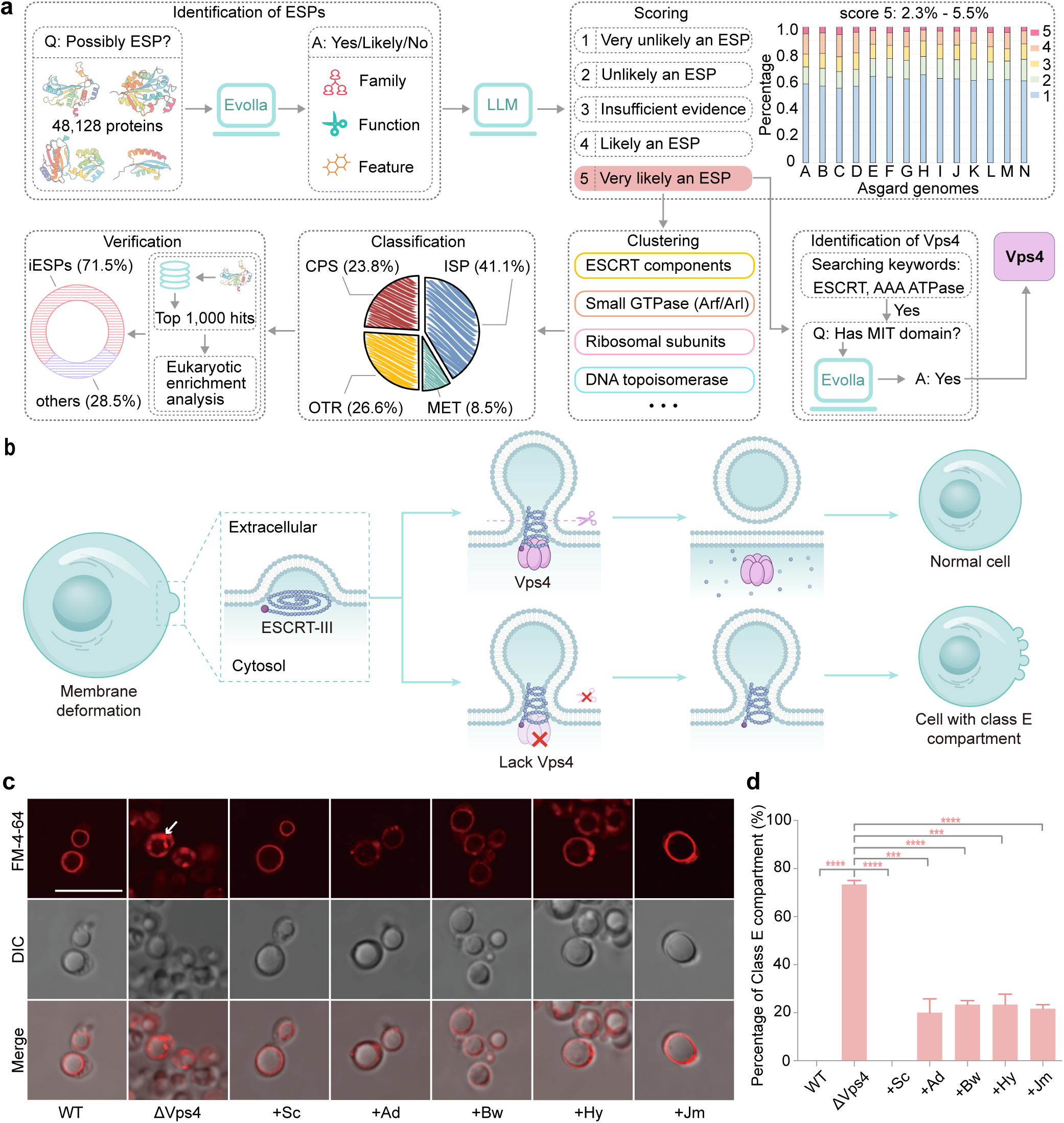
Global identification of Asgard archaeal eukaryotic signature proteins (ESPs) and functional characterization of Vps4. **(a)** General computational pipeline for identifying ESPs. The workflow screens 48,128 protein structures across 14 Asgard genomes. Evolla is queried to provide a classification decision and rationale based on protein family, function, and features. Responses are scored by an LLM on a 1-5 scale. High-confidence candidates (score 5), which account for 2.3-5.5% of the proteome per genome (with genomes A to N corresponding to the Asgard genomes listed in Supplementary File 4, Table 1), are selected for downstream Vps4 identification and functional clustering. Clusters are divided into four categories-ISP (information storage and processing), CPS (cellular processes and signaling), MET (metabolism), and OTR (others)-followed by structural verification using Foldseek. **(b)** Schematic of Vps4, the ESP selected for experimental verification. Vps4 promotes ATP-dependent membrane scission and ESCRT-III disassembly; its absence leads to the formation of class E compartments. **(c)** Vacuolar morphology visualized by FM 4-64 staining. The Δ*vps4* (*vps4* null mutant) strain exhibited the hallmark class E compartment (marked with a white arrow) adjacent to the vacuole, which was fully rescued by native *S. cerevisiae* Vps4 (+Sc). Expression of Asgard Vps4 homologs, including *Atabeyarchaeum deiterrae* (+Ad), *Borrarchaeum weybense* (+Bw), *Hermodarchaeum yapensis* (+Hy), *Jordiarchaeum madagascariense* (Jm), partially reversed this defect. Scale bars, 10 *µ*m. **(d)** Quantification of class E compartments. Expression of Asgard Vps4 reduced compartment accumulation from 70% in the Δ*vps4* strain to 20%. Data are presented as means *±* SD from three biological replicates (n=20 cells per experiment). Statistical significance was determined using an unpaired two-tailed Student’s t-test (***, *p <* 0.001; ****, *p <* 0.0001). Abbreviations include iESP (isomorphic ESP), WT (wild type), and DIC (differential interference contrast).

To experimentally validate the biological relevance of Evolla’s predictions, we focused on the Endosomal Sorting Complex Required for Transport (ESCRT) machinery. Traditionally considered exclusive to eukaryotes [51], this machinery relies on the Vps4 ATPase and ESCRT-III subunits to mediate membrane remodeling [52, 53]. Mechanistically, Vps4 promotes the ATP-dependent scission of membranes and the disassembly of ESCRT-III polymers (Fig. 4b). A critical distinction between eukaryotic Vps4 and its canonical archaeal homolog (Cdv ATPase) is the presence of an N-terminal microtubule-interacting and transport (MIT) domain, which is essential for recognizing ESCRT-III motifs [54].

We refined our screening by subjecting the initial candidates to an iterative semantic interrogation. Following a keyword-based filtration for ‘ESCRT-III’ and ‘ATPase’, we explicitly probed Evolla with the query: ‘Does this protein have an MIT domain?’ (Fig. 4a, Methods 7.15). This targeted inquiry enabled the discrimination of eukaryotic-like Vps4 candidates from canonical archaeal Cdv homologs. From this curated pool, we selected four Vps4 representatives spanning diverse Asgard lineages (Ca. Atabeyarchaeum, Borrarchaeum, Hermodarchaeum, and Jordiarchaeum) to assess their functional robustness.

We reasoned that if Evolla’s semantic identification was accurate, these proteins from uncultured Asgard lineages should exhibit functional equivalence to their modern eukaryotic counterparts. To test this hypothesis, we performed heterologous complementation assays in *S. cerevisiae* (Methods 7.17, 7.18, 7.19, 7.20, 7.21). Deletion of the native *vps4* gene is lethal to yeast under heat stress. Remarkably, expression of the Asgard homologs partially complemented this phenotype, restoring cell viability under restrictive conditions (Extended Fig. S6a). Subsequent characterization confirmed that this rescue was underpinned by genuine mechanistic conservation. Biochemical analysis demonstrated that the candidates possess intrinsic ATPase activity—a prerequisite for ESCRT function (Extended Fig. S6b). Crucially, at the cellular level, the Asgard proteins orchestrated the remodeling of endosomal membranes. While the *vps4* mutant cells accumulated aberrant ‘class E’ prevacuolar compartments, the introduction of Evolla-identified candidates partially reversed this defect, effectively re-establishing proper vacuolar architecture (Fig. 4c, d).

Building on the experimental validation of specific targets, we next assess the global accuracy of Evolla across the full candidate spectrum. We performed semantic clustering on the initial 1,855 candidates based on their functional landscapes, identifying distinct 67 groups (Fig. 4a, Supplementary File 4, Table 2). These groups encompassed known ESPs, such as ribosomal subunits, actin family proteins, and ESCRT components. The candidates were categorized into three major functional classes: Information Storage and Processing (ISP, N=762), Cellular Processes and Signaling (CPS, N=441), and Metabolism (MET, N=159). Proteins (N = 493) that did not cluster into these primary categories—including those identified as viral or nonspecific functions—were assigned to an ‘Others’ (OTR) group.

To benchmark the robustness of our predictions, we compared our candidates against the classification of isomorphic ESPs (iESPs) [55]. The identification of iESPs relies on a rigorous pipeline integrating pangenome-wide structural modeling, structure-based database searches, and statistical assessment of eukaryotic structural overrepresentation. Applying this analytical framework to our dataset (Methods 7.16), we found that 71.5% of the Evolla-predicted candidates (N=1,326) aligned with the strict criteria for iESPs (Fig. 4a, Supplementary File 4, Table 2). The strong concordance validates that Evolla effectively internalizes deep evolutionary patterns, highlighting a methodological shift: instead of identifying evolutionary signals through explicit, rule-based pipelines, Evolla captures the high-dimensional structural and functional logic within the semantic space. Furthermore, the remaining candidates that do not strictly classify as iESPs may represent divergent homologs that elude rigid structural thresholds, suggesting that these semantic predictions warrant further investigation to uncover potentially overlooked evolutionary relationships.

## 5 Discovery of a Novel Deep-sea PET Hydrolase via Evolla

The global accumulation of polyethylene terephthalate (PET) necessitates the rapid discovery of robust plastic-degrading enzymes [56]. While the exponential growth of environmental genomic data offers a vast reservoir of potential biocatalysts [57], mining this sequence space remains a bottleneck, requiring efficient navigation from raw genomic data to functional characterization. Here, we demonstrate an AI-guided discovery workflow employing Evolla as a generative reasoning engine to streamline the path from environmental isolates to the validation of a novel PET hydrolase.

We focused our search on the deep-sea microbiome, a largely unexplored reservoir of biological diversity [58]. Specifically, we targeted a *Pseudomonas sp.* isolate that we previously enriched on PET as the sole carbon and energy source from Pacific Ocean sediments collected across ten distinct sites (138^◦^26^′^-153^◦^56^′^ W, 10^◦^02^′^-13^◦^21^′^ N) [59]. To pinpoint the specific enzyme responsible for its PET-degrading capability, we performed whole-genome sequencing to define a search space of 4,182 predicted protein-coding genes (Methods 7.22). This vast dataset presented a significant challenge for the specific identification of PET hydrolases (Fig. 5a).

**Fig. 5.**
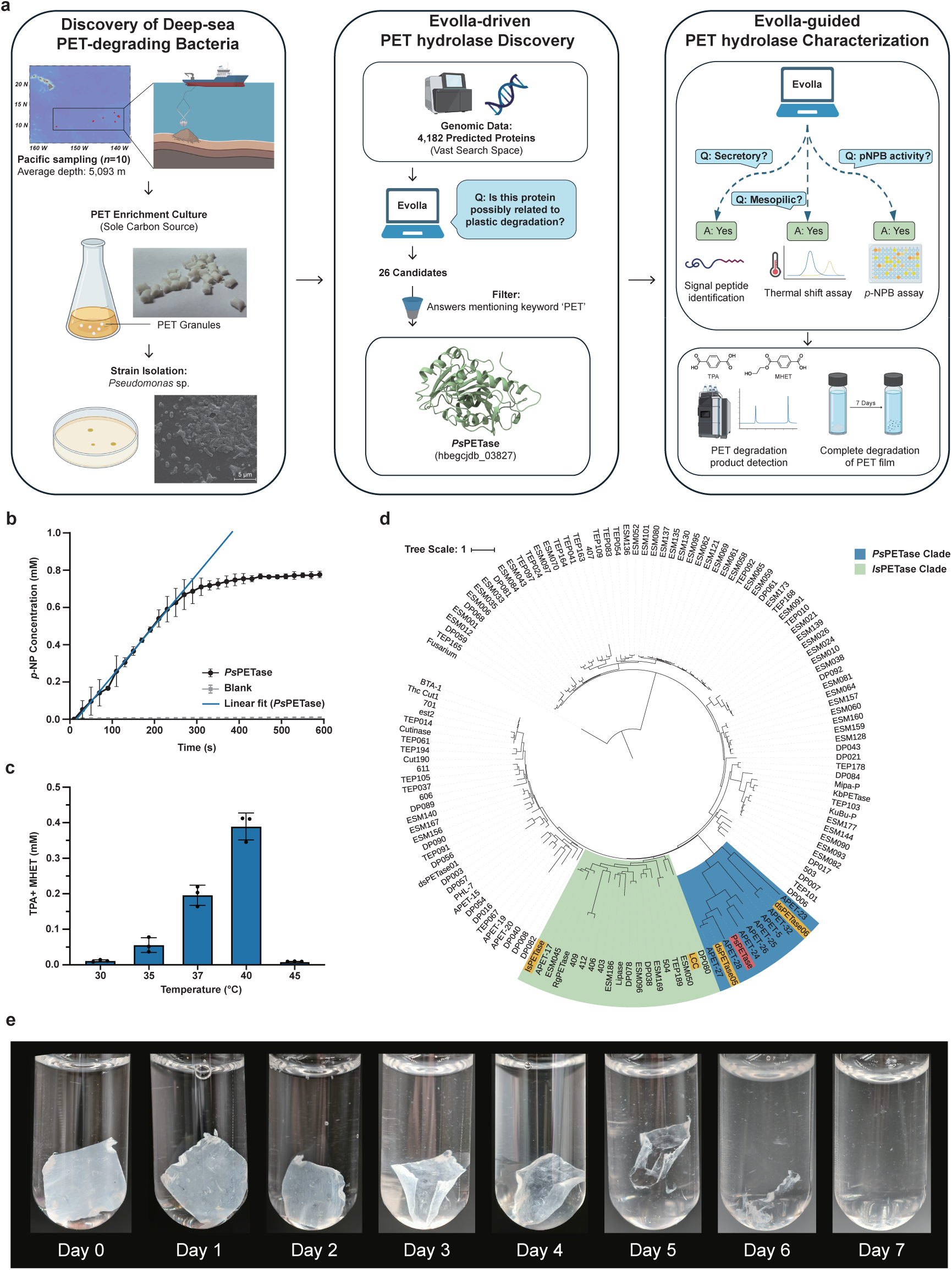
Discovery and Evolla-guided functional mining of the deep-sea PET hydrolases, *Ps*PETase. **(a)** Schematic of the Evolla-guided PET hydrolases discovery and characterization workflow. Deep-sea sediment samples (average depth 5,093 m) yielded a *Pseudomonas sp.* isolate through PET-enrichment culture. Evolla was deployed as a generative reasoning engine to screen 4,182 predicted protein-coding genes via natural language interrogation, identifying *Ps*PETase (Supplementary Table S16) as a high-priority candidate. Subsequent interactive dialogue with Evolla guided experimental design, including signal peptide truncation, mesophilic property prediction, and the selection of validation assays. **(b)** Esterase activity of purified *Ps*PETase determined by the hydrolytic release of *p*-nitrophenol (*p*-NP) from *p*-nitrophenyl butyrate (*p*-NPB). The reaction was monitored in 50 mM KH_2_PO_4_-NaOH buffer (pH 8.0) at 37 °C. *Ps*PETase exhibited robust activity with a linear initial velocity (v0) of 0.164 mM min*^−^*^1^ (represented by the blue fit). Data represent mean *±* s.d. from three independent experiments (n = 3). **(c)** Temperature-dependent solvent-cast PET (scPET) films depolymerization profile of *Ps*PETase. Activity was measured by quantifying released aromatic monomers (TPA and MHET) via HPLC after 24 h of incubation with amorphous PET films. Consistent with the mesophilic classification by Evolla, the enzyme exhibits an optimal reaction temperature of 40 °C. Bars and error bars represent mean *±* s.d. of triplicates; solid circles denote individual data points. **(d)** Maximum-likelihood phylogenetic analysis of *Ps*PETase relative to 137 experimentally validated PET hydrolases (Supplementary File 5), rooted with *Fusarium solani* cutinase as the outgroup. *Ps*PETase defines a distinct marine lineage (*Ps*PETase clade, blue) that is evolutionarily divergent from terrestrial archetypes such as *Is*PETase and LCC (green). The tree was constructed using the LG+F+I+R5 model; the scale bar represents amino acid substitutions per site. **(e)** Time-course macroscopic degradation of scPET films. scPET films were incubated with 1000 nM *Ps*PETase, with buffer and enzyme replenished every 24 h. The series show substantial film thinning and complete degradation within 7 days.

To navigate this dataset, we deployed Evolla for large-scale functional screening (Methods 7.23). Through a query-driven approach targeting specific biotechnological traits, we directly assessed catalytic potential via natural language interrogation. We queried the complete dataset with the prompt: ‘Is this protein possibly related to plastic degradation?’ This initial screen filtered the pool to 26 candidates. Subsequent filtering for the keyword ‘PET’ within the model’s responses refined this pool to a single high-priority target *Ps*PETase from *Pseudomonas sp.* (Fig. 5a, Supplementary Table S16 for the sequence).

Following candidate selection, Evolla transitioned from a high-throughput discovery tool to an interactive partner for experimental design (Fig. 5a). To guide cloning and expression strategies, we queried the biophysical properties of the target (Supplementary Table S15 for specific questions). Guided by Evolla’s prediction that *Ps*PETase is secretory, we used SignalP 5.0 [60] to identify and truncate signal peptides for optimized mature protein expression (Methods 7.24). Furthermore, Evolla classified *Ps*PETase as mesophilic. This prediction was experimentally validated via thermal shift assays, which determined a melting temperature *T_m_*of 43.70°C (Extended Fig. S7, Methods 7.26). This value is comparable to the 46.8°C reported for the representative mesophilic *Is*PETase [61], confirming *Ps*PETase as a typical mesophilic PET hydrolase. This accurate prediction obviated the need for broad temperature screening, enabling us to focus activity assays on a high-resolution gradient (30–45°C).

To validate catalytic function, we adopted a stepwise verification strategy, moving from generic substrate screening to specific polymer degradation. Given that PET hydrolysis fundamentally relies on ester bond cleavage, we first utilized *p*-nitrophenyl butyrate (*p*-NPB) as a rapid surrogate substrate to confirm basal esterase activity. Consistent with Evolla’s prediction, *Ps*PETase exhibited robust hydrolytic capability against *p*-NPB, maintaining a linear initial velocity of 0.164 mM min^−1^ (Fig. 5b, Methods 7.27). Having established this fundamental catalytic prerequisite, we then proceeded to stringent polymer degradation tests using High-Performance Liquid Chromatography (HPLC). PET degradation assays confirmed that *Ps*PETase effectively depolymerized PET, releasing signature monomers—terephthalic acid (TPA) and mono-(2-hydroxyethyl) terephthalate (MHET)—with an optimal reaction temperature of 40°C (Fig. 5c). Macroscopic degradation was confirmed by monitoring solvent-cast PET films; notably, *Ps*PETase treatment resulted in complete degradation of the PET film within 7 days (Fig. 5e, Methods 7.25).

Phylogenetic analysis of 137 validated PET hydrolases (Supplementary File 5) places *Ps*PETase into a distinct lineage—designated here as the *Ps*PETase clade—that is evolutionarily distant from well-characterized terrestrial PET hydrolases such as *Is*PETase [62] and LCC [63] (Fig. 5d). This clade encompasses several recently identified marine-derived enzymes [57], including dsPETase05 and dsPETase06, pointing to a specialized marine lineage that probably evolved independently of its terrestrial homologues.

Collectively, these results demonstrate that Evolla bridges the gap between genomic potential and validated function. Its contribution is twofold: first, it provides a direct, holistic prediction of protein function—transcending simple family membership—by answering specific queries regarding catalytic potential; second, it facilitates an interactive dialogue that transforms the characterization process. By providing actionable insights into secretory status, substrate specificity, and thermal adaptation, Evolla shifts the paradigm from static annotation to dynamic, conversational scientific discovery, establishing a robust framework for mining functional enzymes from biodiversity.

## 6 Discussion

Just as AlphaFold expanded our structural view of the proteome, Evolla aims to decode the functional logic embedded within these molecular representations. By coupling sequence and structure with generative semantic reasoning, our results suggest that large protein-language models can transcend surface-level similarity to perform reasonable biological inference. Evolla moves beyond mere classification to articulate the mechanistic context of protein function, bridging the gap between static structural data and actionable biological insight.

Our computational benchmarks validate Evolla as an analytical engine capable of navigating the ‘evolutionary twilight zone’ [64]. Where traditional alignment tools depend on explicit sequence or structural conservation, Evolla captures ‘functional semantics’—abstract properties that persist even as physical homology fades. By identifying functional relatives in the low-homology regime and rivaling the supervised specialist in enzymatic classification without task-specific fine-tuning, Evolla signals a shift in protein analysis. These findings suggest that the model has internalized complex, non-linear rules of protein evolution, moving beyond explicit pattern matching to facilitate context-aware biological inference.

Beyond quantitative metrics, the identification and wet-lab validation of Asgard Eukaryotic Signature Proteins (ESPs) and the novel PET hydrolase illustrate a new frontier in researcher-model interaction. These cases highlight Evolla’s utility not merely as a static predictor, but as an interactive heuristic tool. Rather than relying on laborious phylogenetic pipelines, the success of these experiments depended on ‘iterative semantic interrogation’; through multi-turn dialogue, Evolla assisted in hypothesis refinement by identifying MIT domains, verifying secretory status, and assessing thermal adaptation. This transforms annotation from a unidirectional output into a dynamic collaboration, where the model actively supports the design-build-test cycle.

Nevertheless, we emphasize that Evolla remains an empirical approximation derived from inductive generalization rather than a first-principles simulation. Its limitations parallel those of structural foundation models like AlphaFold2. First, analogous to AlphaFold2, Evolla is most reliable on well-ordered single chains; whether operating on structural inputs or in sequence-only modes, the model is not explicitly optimized for multi-chain complexes or regions with low predicted structural confidence (low pLDDT). Second, mirroring AlphaFold2’s limited sensitivity to point mutations, Evolla captures broad functional semantics rather than granular quantitative shifts. Just as a single mutation rarely alters an AlphaFold2 backbone, it rarely shifts Evolla’s high-level classification. For example, while the model can identify a protein as fluorescent, it cannot predict how a single substitution alters emission wavelength. Third, the model’s inductive bias toward natural evolution (UniProt) may restrict its generalization to *de novo* designed proteins with non-natural topologies. Finally, as a generative model, it may hallucinate arbitrary identifiers (e.g., PDB codes or UniProt accessions). Unlike functional descriptions, these strings are non-semantic identifiers assigned by databases, lacking intrinsic correlation with protein features; consequently, the model cannot infer them through semantic reasoning and may generate plausible-looking but incorrect strings. Thus, its outputs should be viewed as valuable hypotheses necessitating experimental verification.

Fundamentally, however, the inferential ceiling of any deep learning system is tethered to the density of the functional landscape it traverses. Unlike the vast text corpora fueling breakthroughs in natural language processing, the protein universe suffers from a sparsity of ground-truth labels, with Swiss-Prot offering fewer than 600,000 manually reviewed protein records [65]. Replicating the generative versatility of LLMs in biology will thus necessitate a community-wide expansion of this data horizon, likely driven by high-throughput characterization. In this evolving context, as genomic data accumulation continues to outpace manual curation, computational systems capable of extracting semantic insights from structural and sequence data will become increasingly vital. By offering a framework that renders complex protein data semantically queryable, Evolla represents a significant step toward a more interactive and interpretative paradigm for deciphering protein function.

## 7 Methods

### 7.1 Model architecture

Evolla is designed as a modular generative system bridging protein and human language modalities. The architecture integrates a pre-trained Protein Language Model (PLM), serving as the encoder, with a pre-trained Large Language Model (LLM) acting as the decoder. To leverage pre-existing representations, both the PLM and LLM backbones remain frozen during training.

The core trainable component is a lightweight interfacing module positioned between the frozen encoder and decoder, designed to align the continuous protein representation with the semantic space of the LLM. This module comprises two components: a Sequence Compressor and a Sequence Aligner. The information flow proceeds from the Protein Encoder through the interfacing module into the LLM decoder for textual generation.

#### 7.1.1 Protein encoders

The Protein Encoder extracts high-dimensional features from protein data, yielding a per-residue embedding **e** ∈ R*^L^*^×*d*^*^e^*, where *L* denotes the sequence length and *d_e_* the embedding dimension. We implemented three encoding strategies:

##### Sequence-based encoding

This strategy utilizes ESM2-650M [30]. The representation for an amino acid sequence *x*_aa_ is computed as **e** = ESM2(*x*_aa_).

##### Structure-aware encoding

Utilizing SaProt-650M [14], this strategy processes a sequence of structure-aware tokens *x*_sa_ (combining residue and structural alphabet information) to yield **e** = SaProt(*x*_sa_) (see Methods 7.2).

##### Evolutionary-based encoding

This strategy employs the MSA Transformer [31]. It accepts a multiple-sequence alignment (MSA) *x*_msa_ as input. The model produces embeddings for each sequence in the alignment, which are aggregated via mean pooling across the MSA depth *M* (Ψ : R*^M^*^×*L*×*d*^ → R*^L^*^×*d*^) to generate the final representation **e** = Ψ(MSATransformer(*x*_msa_)) (see Methods 7.3).

#### 7.1.2 Sequence compressor

The Sequence Compressor projects the variable-length protein representation **e** into a fixed-size latent representation **z** (referred to as the protein functional codebook). This compression adapts the protein input to the fixed context window of the LLM (Extended Fig. S2a).

Inspired by the Flamingo architecture [33], the compressor consists of stacked transformer layers, each comprising a cross-attention block and a feed-forward network (FFN), with layer normalization and residual connections. A set of *k* learnable latent query tokens, ***θ***_latent_ ∈ R*^k^*^×*d*^*_z_*, is defined to attend to the protein features. In the cross-attention mechanism, ***θ***_latent_ serves as the query (*Q*), while the protein representation **e** serves as both key (*K*) and value (*V*). For the *i*-th layer with input **z**^(*i*−1)^ (where **z**^(0)^ = ***θ***_latent_), the operation is formalized as:

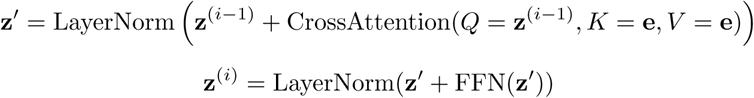

The final output **z** ∈ R*^k^*^×*d*^*_z_* constitutes the fixed-length protein functional codebook utilized by the subsequent alignment module.

#### 7.1.3 Sequence aligner

The Sequence Aligner integrates the protein codebook **z** into the frozen LLM decoder via interleaved adapter blocks. These trainable blocks are inserted between existing LLM layers to modulate internal representations with protein-specific information (Extended Fig. S2b).

Each Aligner block contains a cross-attention mechanism and an FFN, controlled by learnable gating parameters. Given the LLM hidden state from the preceding layer **h***_i_* ∈ R*^T^* ^×*d*llm^ (where *T* is the text sequence length and *d*_llm_ is the LLM’s hidden dimension), **h***_i_* acts as the query (*Q*), while the protein codebook **z** acts as key (*K*) and value (*V*).

Integration is regulated by two scalar gates, *γ*_attn_ and *γ*_ffn_, both initialized to zero to ensure the model initially approximates an identity function. The protein-contextualized signal **c***_i_* is computed and added to the residual stream:

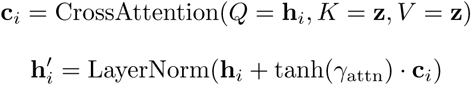

Subsequently, **h**^′^ is processed by the Aligner’s FFN, scaled by the second gate, and added to form the input for the next LLM layer, **s***_i_*_+1_:

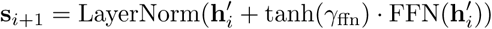

This gated mechanism allows the model to adaptively determine the magnitude of protein context injection during training.

#### 7.1.4 Large language model (LLM) decoder

The language decoder interprets the aligned protein context to generate textual outputs. We employ pre-trained, auto-regressive models from the Llama 3 family [32] (8B and 70B variants). The LLM weights remain frozen.

The protein functional codebook **z** is integrated by modifying the LLM’s forward pass. As described above, Sequence Aligner blocks are interleaved with the frozen LLM layers. For a layer *i* equipped with an aligner, the aligner output **s***_i_*_+1_ serves as the input to the (*i* + 1)-th LLM layer. The final hidden state of the decoder is used to compute the probability distribution over the vocabulary for auto-regressive generation.

### 7.2 Structures in training data

Protein structures were retrieved from AlphaFoldDB [66]. To ensure data quality, we utilized the predicted Local Distance Difference Test (pLDDT) score [3] as a confidence metric. Residues with pLDDT scores below 70 were assigned a special masking token (represented as #) during the tokenization process. This step ensures that the model is trained exclusively on structural features derived from high-confidence regions.

### 7.3 Multiple sequence alignment in training data

Multiple sequence alignments (MSAs) were sourced from the Uniclust30 database (accessed via the OpenFold repository [67]). For proteins not directly indexed in the database, we mapped the corresponding UniProt entries to their cluster representatives within Uniclust30 and adopted the associated cluster-level MSAs. This pipeline resulted in a total of 125,315 MSAs, covering approximately 566,921 sequences.

### 7.4 Training and inference details

#### Model configuration

For Evolla-10B, we utilize SaProt-650M [23] as the protein encoder, which takes protein sequences of up to 1024 structure-aware (SA) tokens as input. Following this, a Sequence Compressor module is employed, consisting of 6 layers, 8 attention heads, 64 latent units, and a feedforward multiplier of 4. We initialize the LLM decoder with weights from HuggingFace’s meta-llama/Meta-Llama-3-8B-Instruct, setting the maximum token length to 736. The Sequence Aligner comprises 8 cross-attention layers, each with an FFN multiplier of 4 and a 0.1 dropout rate applied to the attention probabilities, inserted after every 4 layers of the LLM decoder. For Evolla-80B, we utilize SaProt-1.3B as the protein encoder, with the same configuration for sequence compression. In contrast to the 650M SaProt that is restricted to SA token sequences, SaProt-1.3B supports either amino acid or SA token sequences, enabling Evolla-80B to process inputs in either sequence or structural modalities. We initialize the LLM decoder with weights from HuggingFace’s meta-llama/Llama-3.1-70B-Instruct, setting the maximum token length to 736. The Sequence Aligner consists of 20 cross-attention layers with an FFN multiplier of 1, applying a 0.1 dropout rate to the attention probabilities after every 4 layers of the LLM decoder. In both configurations, the SaProt encoder and Llama3 decoder are frozen, while the Sequence Compressor and Sequence Aligner are trainable, with a total of 1.7 billion trainable parameters for Evolla-10B and 8.2 billion trainable parameters for Evolla-80B. This configuration serves as the default setup for our experiments.

#### Training settings

We trained Evolla-10B using DeepSpeed [68] with the AdamW optimizer [69], setting *β*_1_ = 0.9, *β*_2_ = 0.98, and applying L2 weight decay of 0.05. The model was trained with bfloat16 mixed precision [70], and the learning rate was initially warmed up from 0 to 1 × 10^−5^ over the first 2000 steps, followed by decay to 1 × 10^−7^ according to a cosine annealing schedule [71].Training was conducted on 64 NVIDIA H800 80GB GPUs for 6 days. For Evolla-80B, we utilized FSDP [72] with a hybrid sharding strategy on 128 NVIDIA H800 80GB GPUs, training for 47 days. Other configurations were consistent with those used for Evolla-10B.

Training data mixture strategy.

The training corpus was constructed by integrating (protein, question, answer) triplets derived from two distinct lineages: the manually curated Swiss-Prot dataset and the ProTrek-augmented TrEMBL50 dataset (detailed dataset construction in Methods 7.5). To mitigate potential bias toward the significantly larger TrEMBL50 dataset, we adopted a balanced sampling strategy that enforces a 1:1 mixing ratio. This was achieved by constructing each training batch with an equal number of samples from both sources, irrespective of total dataset size. This ensures the model benefits from the high-quality standards of Swiss-Prot while maximizing the utility of the vast diversity found in TrEMBL50.

#### Generation

For sequence generation, we utilized nucleus sampling (top-*p* = 0.9) and a temperature setting of 0.6. The generation was capped at a maximum of 512 new tokens per sequence.

### 7.5 Data preparation

#### 7.5.1 Information Units in the UniProt database

**Information Units (IUs)** serve as the fundamental building blocks of protein annotation within the UniProt database. An IU is defined as a discrete textual description capturing specific attributes of a protein, its residues, or segments. These annotations are categorized into distinct sections according to the biological nature of the information.

A single protein entry typically contains annotations across multiple sections, and a single section may house multiple distinct IUs. For instance, an enzyme might have annotations in both the ‘Function’ and ‘Catalytic Activity’ sections; within the ‘Cofactor’ section, it may possess multiple entries if it interacts with distinct cofactors. In our framework, each annotation is treated as an independent IU, regardless of its section or context.

To construct our dataset, we extracted IUs from the UniProt database, covering 31 sequence-level sections (describing global protein attributes, e.g., ‘Function’) and 25 residue-level sections (detailing specific residues or regions, e.g., ‘Active Site’). This yielded a total of 56 distinct annotation categories (Supplementary Table S1). Detailed descriptions and examples for each section are provided in the Supplementary File 1.

Our primary dataset was derived from the Swiss-Prot database (Release 2023 03), which comprises 569,793 proteins annotated with 14,818,151 IUs. Swiss-Prot represents the gold standard for protein annotation, characterized by manually curated entries derived principally from experimental evidence and expert review to ensure high accuracy and reliability [73].

We further examined IUs from the TrEMBL database (Release 2023 03), which contained over 2.4 billion IUs. Unlike Swiss-Prot, TrEMBL entries are not manually reviewed [73]. More importantly, we observed that the TrEMBL dataset exhibited significant class imbalance, with certain sections being disproportionately represented (Fig. 1b, middle ring). Consequently, we adopted a strategy to utilize TrEMBL solely for its sequence diversity (see below).

#### 7.5.2 Retrieval strategy in TrEMBL by ProTrek

To overcome annotation sparsity, we incorporated ProTrek [22] to perform computational data augmentation. Specifically, we leveraged ProTrek to retrieve specific, high-confidence textual segments (IUs) from Swiss-Prot entries associated with specific proteins in TrEMBL. This allows us to add rich functional properties for proteins that lack experimental annotation.

We first performed redundancy reduction on the TrEMBL database using MMseqs2 [43], clustering sequences at 50% identity to yield 41,229,217 cluster representatives (referred to as TrEMBL50). Subsequently, we applied ProTrek to map these representatives to relevant text fragments using two complementary strategies: (1) Global Top-10 IUs: For each protein, we selected the 10 textual fragments with the highest retrieval scores from the entire global IU pool, yielding 412,292,170 IUs. (2) Stratified Top-10 IUs: To prevent bias toward specific annotation categories, we stratified the retrieval space into the 56 predefined sections. For each protein, we identified the highest-ranking IU within each section to form a candidate set, from which we then selected the top 10 IUs. This contributed an additional 412,292,170 IUs.

By focusing on retrieving high-confidence information units rather than full descriptions, this approach ensures our model is trained on diverse and structured semantic signals, totaling 824,584,340 IUs.

#### 7.5.3 Question generation

To generate a diverse and realistic corpus of user queries for our (protein, question, answer) triplets, we employed two complementary strategies: a *section-oriented strategy* and a *persona-oriented strategy*.

The *section-oriented strategy* leverages the standardized annotation architecture of the UniProt database to create a foundational set of questions. For each of the 56 predefined annotation sections (Supplementary Table S1), we manually formulated a canonical seed question that directly queries the information within that section. To enhance linguistic diversity and mitigate the risk of the model overfitting to a limited phrasing, each seed question underwent large-scale paraphrasing. We utilized a large language model (GPT-4) to produce 100 distinct linguistic variations for each of the 56 canonical questions, resulting in a comprehensive corpus of section-specific queries (Supplementary File 2).

The *persona-oriented strategy* aims to simulate the context-driven and varied inquiries of real-world users, thereby generating questions that reflect diverse levels of expertise and scientific interests. We established a library of ten distinct user personas, each defined by a detailed textual description of their background and objectives (Supplementary Table S4). For a given protein, its associated set of IUs was provided as context to a large language model. The model was then prompted to adopt three randomly selected personas and, informed by the provided IUs, generate an initial question that each persona would plausibly ask (Supplementary Table S2). To further broaden the scope and complexity of these inquiries, we employed a set of five question evolution strategies (Supplementary Table S5). For each initial question, three evolution directions were randomly selected to guide the LLM in rewriting the query based solely on the original question and the specific evolutionary goals (Supplementary Table S6). This process yielded a total of 12 questions per protein, comprising the three original and nine evolved inquiries.

#### 7.5.4 Answer generation

With a comprehensive corpus of IUs and generated questions established, the final stage of data preparation involved synthesizing a high-quality answer for each (protein, question) pair.

Initially, we evaluated a template-based methodology. For IU types that consist of keywords or non-sentential texts (e.g., ‘Cofactor’), we manually designed templates tailored to their specific data structure. To enhance linguistic variety, each canonical template served as a seed for large-scale paraphrasing, where GPT-4 generated 100 semantically invariant variations (Supplementary File 3). IUs already in a descriptive, narrative format (e.g., ‘Function’) were kept in their original form. However, pilot experiments pairing these templated answers with section-oriented questions revealed significant limitations, including overly brief responses, severe memorization issues, and much worse performance (Extended Fig. S3 and S5b).

To overcome these limitations, we implemented an LLM-driven answer synthesis pipeline leveraging the GLM [74] developed by ZhipuAI. For a given protein, its complete set of associated IUs and a randomly sampled question from our question repository were integrated into a structured prompt. This prompt instructed the LLM to synthesize a single, comprehensive, and well-structured answer (Supplementary Table S7). The primary objective was to ensure the generated text remained factually grounded in the provided IUs while being stylistically and semantically aligned with the specific question, thereby creating a comprehensive and more reliable answer suitable for training Evolla.

Finally, we applied a quality control strategy on these generated answers. For proteins in the Swiss-Prot database, all generated answers were retained without further filtering. In contrast, for proteins in the TrEMBL50 database, we empirically applied a semantic consistency filter to the TrEMBL50 dataset to mitigate potential hallucinations (or inaccuracies). Specifically, we employed ProTrek to calculate a matching score between each protein sequence and its generated description. To ensure high data quality, protein-answer pairs yielding a similarity score below 12 were excluded from the final training corpus.

### 7.6 Dataset partition

To ensure rigorous evaluation against high-quality standards, we constructed our validation and test sets based on the expert-curated Swiss-Prot database (Release 2023 03). Adopting the methodology and the exact data split from ProTrek [22], the data construction process was as follows: Protein sequence identity clustering was performed on all Swiss-Prot entries using MMseqs2 [43] at a 50% identity threshold. From the resulting clusters, 1,000 were randomly sampled to constitute the validation set, and an additional 1,000 clusters—comprising 4,040 proteins and 5,968 protein-question-answer triplets—were selected for the test set. The remaining Swiss-Prot proteins, along with TrEMBL50 sequences automatically annotated by our computational pipeline (detailed as ProTrek+LLM in section 7.5), were assigned to the training set.

To evaluate the model’s performance more precisely across varying difficulty levels, we further stratified the test set. Proteins exhibiting less than 30% sequence identity to any Swiss-Prot training set protein were categorized as the ‘hard’ test set (831 proteins with 917 triplets).

### 7.7 Metrics

The quantitative evaluation of biological validity and functional correctness in generated free-text responses remains an unexplored frontier within the community. Traditional metrics designed for the natural language processing (NLP) field primarily rely on n-gram overlap [75, 76]. Consequently, these metrics prove insufficient for assessing domain-specific biological narratives, as they fail to capture deep semantic nuances or verify factual consistency with biological mechanisms.

To address this limitation, we developed and employed an LLM-based evaluation, termed GPT score, which leverages the advanced reasoning capabilities of LLM to act as an expert evaluator [36]. The reliability and validity of this metric were rigorously established through a two-dimensional assessment (Supplementary Text A.1 for details). Our analysis demonstrated high test-retest stability (ICC = 0.97, Extended Fig. S8c) on the test set and a strong correlation with averaged ratings from three human experts (Spearman’s *ρ* = 0.9594, Extended Fig. S8d) on a smaller sampled dataset, confirming that GPT score serves as a robust proxy for manual evaluation of complex biological narratives.

The GPT score is calculated for each (protein, question, answer) triplet using a structured evaluation prompt (Supplementary Table S10, Extended Fig. S8a). This prompt provides the evaluator LLM with comprehensive context, including all known Information Units (IUs) for the target protein and the generated answer. The LLM is instructed to assign a score from 0 to 100, reflecting the semantic alignment and factual accuracy of the response generated by Evolla with respect to the provided ground-truth information. To mitigate the inherent stochasticity of LLM-based evaluation, each scoring procedure was performed in triplicate with independent queries and the average score was reported as the final result.

Beyond factual accuracy, assessing whether a generated answer is truly relevant to the specific question asked is another critical dimension of our evaluation. To quantify this aspect, we introduced a ‘QA relevance score’ (Supplementary Table S3, Extended Fig. S8b). For this metric, the evaluator LLM is instructed to disregard factual correctness and solely judge whether the provided answer addresses the posed question, assigning a score from 0 to 100. Analogous to the GPT score, this scoring procedure was repeated three times and the average was reported as the final score.

We selected gpt-4o-mini-2024-07-18 as our primary evaluator, as it represented the state-of-the-art at the time of this study and ranked among the top models on the Chatbot Arena Leaderboard [77] (accessed October 6, 2024).

In addition to evaluating quality, we developed a quantitative metric to assess their originality of the generated outputs. The aim was to measure the degree to which a model’s response represents a novel generation rather than a verbatim recitation of the reference knowledge base. To this end, we computed the local text similarity between each generated answer and its corresponding ground-truth annotation in the Swiss-Prot database.

This comparison was performed using the Smith-Waterman algorithm, implemented via the *pairwise2* module in Biopython [78]. The alignment was configured with a match score of 2, a mismatch penalty of −1, a gap open penalty of −5, and a gap extension penalty of −2; only the highest scoring alignment was considered. To create a bounded similarity metric, the raw alignment score (termed the SW score) was normalized by the self-alignment score of the generated answer. Finally, we defined the *Originality* of a response as 1 − Normalized SW Score. Under this metric, a score approaching 1.0 signifies a response with minimal textual overlap with the source annotations, indicating a high degree of rephrasing and synthesis.

### 7.8 Direct Preference Optimization

The DPO pipeline consists of three stages: preference dataset construction, DPO fine-tuning, and generation-based checkpoint selection.

#### 7.8.1 Preference dataset construction

The preference dataset was constructed by sampling 45,000 (protein, question) pairs from the training set. For each pair, we employed the vanilla Evolla model to generate four candidate responses. These candidates were evaluated using our GPT score framework, which assesses the factual consistency of each response against the known Information Units of the target protein. To establish a clear learning signal, we formed preference pairs by designating the highest-scoring response as the ‘chosen’ answer (*t_c_*) and the lowest-scoring response as the ‘rejected’ answer (*t_r_*). To ensure a distinct quality margin, we filtered these pairs to retain only those with a GPT score difference of 10 or greater. This process yielded a final preference dataset, D_pref_, containing 37,544 quadruplets of the form (*p*, *q*, *t_c_*, *t_r_*).

#### 7.8.2 DPO fine-tuning

The DPO algorithm fine-tunes the model policy, *π_θ_*, using a frozen copy of the vanilla Evolla model as a reference policy, *π*_ref_. This reference model implicitly constrains the optimization, preventing the active policy (*π_θ_*) from deviating excessively from the original distribution while maximizing the likelihood margin between preferred and rejected responses.

The model is trained to minimize the DPO loss function [29]:

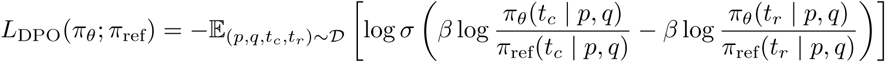

where *p* is the protein context, *q* is the question, *σ* is the logistic sigmoid function, and *β* is a hyperparameter controlling the deviation from the reference policy (KL penalty). By minimizing this loss, the policy *π_θ_* is optimized to assign a higher likelihood to the chosen answer *t_c_* and a lower likelihood to the rejected answer *t_r_*, relative to the reference policy *π*_ref_.

#### 7.8.3 Checkpoint selection

In standard supervised learning, the checkpoint with the lowest validation loss is typically selected as the optimal model. However, in preference alignment, the DPO loss serves as a proxy for relative preference alignment, rather than a direct indicator of the absolute generation quality. A low DPO loss has learned to distinguish between chosen and rejected samples, but it does not guarantee improved factually accurate or coherence in open-ended generation. Consequently, to identify the checkpoint with the superior overall performance, we implemented a generation-based evaluation strategy.

To balance evaluation rigor with computational efficiency, a validation subset of 500 (protein, question) pairs was randomly sampled from the primary validation set. During DPO training, model checkpoints were saved at the end of each epoch. Following training, we generated responses for this validation subset using each saved checkpoint. These responses were evaluated using GPT score, and the checkpoint yielding the highest average score was selected as the final, DPO-aligned model for all subsequent experiments.

### 7.9 Retrieval augmented generation

We integrated a Retrieval-Augmented Generation (RAG) framework to enhance the factual grounding of the model by dynamically incorporating information from an external, expert-curated knowledge base. The RAG pipeline comprises two primary stages: (1) information retrieval and (2) augmented generation.

For the information retrieval stage, we designated the Swiss-Prot database (Release 2023 03) as the external knowledge corpus, leveraging its high-quality, manually annotated protein entries. To ensure a fair evaluation, the entry corresponding to the query protein was dynamically excluded from the retrieval corpus during each inference process. We developed and evaluated two distinct retrieval strategies: *direct retrieval* and *iterative retrieval*.

Our first approach, direct retrieval, follows a hierarchical two-step process (Extended Fig. S4a). First, at the protein level, we employ the ProTrek retrieval tool to identify candidate proteins. From a pool of proteins with a ProTrek sequence-to-sequence similarity score exceeding 40 (excluding the query protein itself), we randomly sample up to three out of the top five candidates to ensure diversity. Subsequently, for each of these selected proteins, we perform text-level retrieval within their respective Swiss-Prot entries. We calculate the semantic similarity between the user’s query and the headers of the metadata sections (e.g., ‘Function’, ‘Catalytic activity’) using ProTrek. We then retrieve the full textual content for up to four of the highest-scoring sections per protein, provided their relevance score exceeds a threshold of 15. This procedure yields a maximum of 12 retrieved textual segments for downstream generation.

The second strategy, iterative retrieval, implements a generate-then-retrieve paradigm to refine the text-level search (Extended Fig. S4b). While the protein-level search remains identical to the direct retrieval approach, the text retrieval mechanism is modified to leverage the model’s intrinsic knowledge. Instead of querying with the raw user question, we first prompt Evolla to generate a preliminary response (or ‘hypothetical answer’) based solely on the original protein and question inputs. This generated response is then utilized as a semantic query to rank the full textual content of all annotations associated with the selected proteins. The top-four textual annotations exhibiting the highest relevance score to this hypothetical answer are selected (threshold score *>* 15). This content-centric approach guides the retrieval process using the model’s parametric knowledge, focusing the search on information that validates or corrects upon its preliminary hypothesis.

In the final augmented generation stage, the retrieved textual annotations from either strategy are concatenated with the original user query (Supplementary Table S18). This augmented text is integrated into a structured prompt alongside the query protein and fed to Evolla to produce the final response (Extended Fig. S4c).

### 7.10 Comparison with General LLMs

Establishing a direct comparison between the domain-specialized Evolla and general-purpose LLMs presents inherent challenges. Unlike Evolla, which is architecturally designed to interpret protein data, LLMs process protein sequences solely as strings of amino acid characters, lacking intrinsic biological context. Consequently, when presented with a raw protein sequence alongside a functional query, LLMs typically exhibit a tendency to generate generic or refusal responses, often defaulting to recommendations for specialized bioinformatics tools.

To mitigate this limitation and establish a rigorous basis for comparison, we designed a specialized, multi-faceted prompt (Supplementary Table S17) that standardizes the analytical task. This prompt explicitly instructs the LLM to conduct a comprehensive analysis covering a predefined spectrum of key biological attributes.

The scientific accuracy of the resulting outputs from both Evolla and the LLMs was quantitatively evaluated using our GPT score framework. For these experiments, we utilized the test set defined in Section 7.6 and the DPO-enhanced 10B parameter version of Evolla (Evolla-10B-DPO).

### 7.11 Comparison with recent protein-language generative models

We conducted a comparison with two recent protein-language generative models: ChatNT [26] and InstructBioMol [40]. Given the distinct input modalities required by these models, we developed specific data preparation pipelines to adapt our protein-centric test set. Our evaluation focused on the capacity of each model to generate accurate responses when provided with a biomolecular input and a natural language query.

#### Benchmarking against ChatNT

As ChatNT is a conversational model primarily designed for nucleotide sequences, benchmarking necessitated mapping our test set proteins to their biologically corresponding DNA sequences. Since direct computational reverse-translation is infeasible due to the degeneracy of the genetic code, we implemented a database-centric mapping pipeline. First, UniProtKB accession IDs were mapped to the RefSeq Protein database via the UniProt ID mapping service. Of the 4,040 test proteins, 3,362 yielded at least one corresponding RefSeq ID. For entries with direct transcript links (prefixes ‘XP’ or ‘NP’), DNA sequences were retrieved directly from NCBI. For non-redundant protein records (prefix ‘WP’), which lack a direct link to a single DNA record, we identified the ‘identical protein group’ and selected the DNA sequence of the group’s representative member. In cases where a single protein ID linked to multiple DNA sequences, one was randomly selected to resolve ambiguity. This comprehensive process yielded a final set of 2,996 DNA sequences suitable for ChatNT.

#### Benchmarking against InstructBioMol

The InstructBioMol framework requires a structural motif-based representation (conceptually functioning as a ‘protein fingerprint’) as an auxiliary input alongside the protein sequence. To generate these representations, we employed the gen protein motif.py script provided in the official InstructBioMol GitHub repository^1^. To ensure strict reproducibility, all protein fingerprints were generated using the code from a specific commit hash (ID: 5f63c6bd7c48a08b60d7994c4073070d8781d169).

All comparative experiments in this section utilized the same test set and model version (Evolla-10B-DPO) as described above.

### 7.12 Temporal hold-out set of newly annotated proteins

To assess the generalization capability of Evolla on entirely novel data and to simulate a real-world predictive scenario, we constructed an additional temporal hold-out test set. This set comprised proteins that were newly curated and deposited into the Swiss-Prot database subsequent to the data snapshot date of our primary training corpus.

Specifically, we identified all protein entries present in Swiss-Prot Release 2024 03 that were absent from Swiss-Prot Release 2023 03 (the baseline for our training, validation, and test datasets), identifying a total of 1,896 proteins. This strict temporal separation ensures that the proteins in this hold-out set were annotated and released to the public after the model was trained. From this collection of newly annotated proteins, we further filtered for entries assigned an Enzyme Commission (EC) number, resulting in a subset of 607 proteins. This subset served as a dedicated benchmark for evaluating the performance of Evolla on the specific task of EC number prediction for newly characterized enzymes.

### 7.13 Instructional response matching (IRM)

Evaluating generative models like Evolla presents a fundamental challenge: traditional bioinformatics tools output structured data—such as alignment scores (e.g., BLAST E-values) or discrete class labels (e.g., CLEAN’s EC numbers)—whereas Evolla generates unstructured natural language. To bridge this format gap and enable a rigorous, unified comparison, we introduce the **Instructional Response Matching (IRM)** framework.

IRM reformulates diverse biological tasks into a standardized semantic retrieval problem. Instead of measuring accuracy based on rigid output formats, IRM evaluates the semantic alignment between the model’s generation and a task-specific target. The framework operates in three steps:

1. **Instruction (***T_I_* ): We prompt the model with a task-specific query (e.g., ‘What is the catalytic activity?’ or ‘What is the function of this protein?’).
2. **Response (***T_P_* ): The model generates a descriptive text *T_P_* for the query protein based on the instruction.
3. **Matching (***S*): We compute a semantic similarity score *S*(*T_P_, T*_target_) between the generated response and a target text *T*_target_ using a pre-trained text embedding model.

Crucially, by varying the definition of the target text *T*_target_, IRM adapts to two distinct evaluation paradigms used in our experiments:

- **Label-Matching for classification (Methods 7.13.1):** To compare against classifiers, *T*_target_ is defined as the natural language description of a ground-truth label (e.g., the definition of an EC number). The model’s prediction is correct if *T_P_* is semantically closer to the correct label’s description than to others.
- **Peer-Matching for retrieval (Methods 7.13.3-7.13.5):** To compare against homology search tools (BLASTp, Foldseek), *T*_target_ is defined as the generated description of another protein in the dataset. This allows us to perform all-versus-all searches in semantic space, retrieving functionally related proteins (defined by GO terms or KEGG pathways) based on the similarity of their generated descriptions.

This framework effectively decouples the evaluation metric from the model architecture, allowing us to benchmark Evolla’s zero-shot generation capabilities directly against specialized, state-of-the-art baselines across multiple domains.

#### 7.13.1 EC number prediction via label-matching

To benchmark Evolla against specialist classifiers, we applied the **Label-Matching** mode of the IRM framework (Methods 7.13) to the task of Enzyme Commission (EC) number assignment. In this setting, the target set {*T*_EC_} consists of the natural language descriptions for all official EC categories (Methods 7.13.2).

The procedure follows the standard IRM pipeline with task-specific adaptations. First, the instructional prompt *T_I_*is set to ‘What is the catalytic activity of this protein?’, guiding Evolla (specifically the Evolla-10B-DPO version) to generate a functional description *T_P_*. Second, we compute the semantic similarity between *T_P_* and every target description in {*T*_EC_} using the text-embedding-3-large model.

To convert these continuous similarity scores into discrete class predictions, we employ the maximum separation algorithm [41]. This algorithm ranks the similarity scores and identifies the largest margin between consecutive matches; all candidates ranking above this margin are selected as predictions. This approach effectively translates Evolla’s generative output into multi-label classification results, enabling a direct quantitative comparison with contrastive learning baselines like CLEAN. All results reported in this benchmark were generated using the Evolla-10B-DPO version.

#### 7.13.2 Target set construction: converting EC numbers to text

To enable the **Label-Matching** mode described in Methods 7.13.1, we constructed a comprehensive target set {*T*_EC_}, where each discrete Enzyme Commission (EC) number is mapped to a rich natural language description. This process bridges the gap between the hierarchical nomenclature system and the semantic embedding space.

The construction pipeline involved three steps:

1. **Data acquisition:** The primary list of valid EC numbers was sourced from the official Enzyme Nomenclature list, maintained by the Nomenclature Committee of the International Union of Biochemistry and Molecular Biology (NC-IUBMB) [79] (accessed March 2024, hosted at Queen Mary University of London). This provided the authoritative backbone of the classification hierarchy.
2. **Metadata enrichment:** For each entry, we retrieved detailed structured metadata from the KEGG ENZYME database [80], including the canonical name, class, systematic name, the IUBMB-defined chemical reaction, substrate(s), product(s), and any associated functional comments.
3. **Semantic synthesis:** We employed GPT-4o to synthesize these structured fields into a coherent, descriptive paragraph (*T*_EC_) for each enzyme (see Supplementary Table S11 for prompt details and Supplementary Table S12 for examples).

This procedure transforms the rigid tabular data into a semantic textual format, allowing the IRM framework to perform functional assignment via semantic similarity matching.

#### 7.13.3 All-versus-all functional search via Peer-Matching

For retrieval tasks, we adapted the IRM framework into its **Peer-Matching** mode to perform all-versus-all functional similarity searches. This approach provides a semantic analogue to traditional homology-based methods (e.g., BLASTp). Unlike the classification task, which matches a query against a fixed set of external labels, this mode computes relationships within a candidate set of proteins {*P*_1_*, P*_2_*, …, P_n_*}.

The process proceeds as follows:

1. **Generation:** For every protein in the set, we generate a functional description *T_Pi_* using the query prompt *T_I_* : ‘What is the function of this protein?’.
2. **Matrix construction:** We compute a pairwise semantic similarity matrix by scoring every pair of generated descriptions (*T_Pi_*, *T_Pj_*) using the text-embedding-3-large model. This results in an *n*×*n* matrix where each entry quantifies the semantic functional similarity between two proteins.
3. **Retrieval:** For any query protein *P_i_*, its corresponding row in the matrix serves as a ranked list of candidates.

By sorting these scores in descending order, we can identify functionally similar proteins solely based on Evolla’s textual understanding. The quality of this ranking is then evaluated against ground-truth functional associations, such as shared Gene Ontology (GO) terms or KEGG pathways, as detailed in the following sections.

#### 7.13.4 Gene Ontology detection evaluation

We evaluated Evolla’s semantic retrieval performance on a held-out test set of 4,040 proteins (strictly separated from training data, see Methods 7.6). Using the **Peer-Matching** protocol described in Methods 7.13.3, we performed an all-versus-all search where each protein served as a query against the entire test set. Performance was benchmarked against three established homology-based methods: BLASTp and MMseqs2 (sequence alignment) and Foldseek (structural alignment).

##### Retrieval protocol

A ‘functionally related pair’ was defined as any two proteins sharing at least one GO term. To determine valid hits from the ranked results, we applied a dynamic termination criterion consistent with previous studies [7, 22]. Specifically, for each query, we traversed the ranked list and ceased retrieval upon encountering the fifth false positive (defined as a protein sharing no GO annotations with the query). All true positives identified prior to this cutoff were retained as ‘retrieved positive pairs’.

##### Stratified performance analysis

To rigorously assess generalization, we computed recall rates stratified by structural and sequence similarity. First, we identified the global set of all ground-truth pairs within the test set and calculated their pairwise TM-scores [44] and sequence identities (via MMseqs2). We then partitioned both the ground-truth set and the set of ‘retrieved positive pairs’ into bins (TM-score width: 0.1; Sequence Identity width: 10%). For each bin, the recall rate was calculated as the ratio of retrieved positive pairs to the total number of ground-truth pairs falling within that specific similarity range. This stratification reveals the model’s ability to detect functional relationships even in the absence of significant sequence or structural homology.

All results reported in this benchmark were generated using the Evolla-10B-DPO version.

#### 7.13.5 KEGG pathway reconstruction evaluation

We further assessed Evolla’s ability to reconstruct complex biological pathways by applying the **Peer-Matching** protocol (defined in Methods 7.13.3) to the proteome of the Asgard archaeon HC1 [45] (6,540 proteins). Unlike the GO benchmark which looks for broad functional similarity, this task tests the model’s capacity to cluster proteins into specific metabolic modules.

##### Ground truth and baselines

Ground truth pathway memberships were established by mapping the proteome to KEGG Orthology (KO) identifiers using KofamKOALA [81], followed by linking these KOs to global KEGG pathways. We compared Evolla against two categories of baselines:

- **Homology-based:** BLASTp, MMseqs2 (sequence), and Foldseek (structure).
- **Annotation-based:** A text-embedding baseline derived from InterProScan [46]. For this, we concatenated all textual annotations for each protein into a single string and encoded it using the same embedding model (text-embedding-3-large) which was also employed to encode Evolla’s generated responses, allowing for a direct comparison of ‘generated text’ versus ‘retrieved annotation’.

##### Evaluation metric

Performance was quantified using Recall@K (*K* ∈ [1, 100]). For each query protein, this metric measures the fraction of its true pathway partners (proteins sharing the same KEGG pathway) that appear within the top *K* retrieved candidates. The final curves (Fig. 3h) report the average Recall@K across all pathway-annotated proteins, directly reflecting the model’s utility in pathway reconstruction tasks. All results reported in this benchmark were generated using the Evolla-10B-DPO version.

### 7.14 Using computational tools

#### 7.14.1 Foldseek

We used Foldseek with the **70cea935733e0f70e3e10a0572b45ba6e21bfa2d** version. The command line was executed with the following parameters: **foldseek easy-search structure dir structure dir aln.m8 tmp -s 15.0 -e inf –max-seqs 1000000 –min-ungapped-score 0**

#### 7.14.2 MMseqs2

We used MMseqs2 with the **b804fbe384e6f6c9fe96322ec0e92d48bccd0a42** version. The command line was executed with the following steps:

1. **mmseqs createdb protein.fasta proteinDB**
2. **mmseqs prefilter proteinDB proteinDB protein pref DB -s 15.0 –max-seqs 1000000 –min-ungapped-score 0**
3. **mmseqs align proteinDB proteinDB protein pref DB protein aln DB -e inf**
4. **mmseqs convertalis proteinDB proteinDB protein aln DB aln.m8**

#### 7.14.3 BLASTp

We used BLASTp with the **Protein-Protein BLAST 2.16.0+** version. The command line was executed with the following steps:

1. **makeblastdb -in protein.fasta -dbtype prot -parse seqids -out tmp/blastp db**
2. **blastp -query protein.fasta -db tmp/blastp db -outfmt 6 -out alnRes blastp -num threads 32 -evalue 1e6 -max target seqs 50000**

#### 7.14.4 CLEAN

We used CLEAN from the Github repository located in https://github.com/tttianhao/CLEAN (commit id: **f2bf2a4f497fa2cc87dac2a1bb314fee587c0a15**). We ran CLEAN with max-separation model following the *README* instruction.

#### 7.14.5 InterProScan

The version of InterProScan we use is **5.73-104.0**, the default command is used as follow:

**interproscan.sh -i input file name -o output file name -f tsv**

#### 7.14.6 KofamKOALA

We used the online server of KofamKOALA (KEGG release 115.0) to perform the analysis of the proteins.

### 7.15 Identification of ESPs by Evolla

Based on the GTDB database (release 226) [82], we selected 14 high-quality genomes representing 12 classes of Asgard archaea and downloaded their corresponding amino acid sequences. Two genomes of isolated Lokiarchaeia [83, 84] and two isolated Hodarchaeales within Heimdallarchaeie [85] were included (Supplementary File 4, Table 1).

We first predicted the structures for all proteins from these genomes using AlphaFold3 [2]. Each predicted protein structure was then queried using the 10B-parameter version of Evolla (with the DPO and Iterative RAG enhancement) with the question, ‘Is this protein possibly a eukaryotic signature protein?’ to generate an answer. To systematically score these answers, we employed the gpt-4o-mini-2024-07-18 large language model (LLM) for its cost-effectiveness. Following a specific prompt (see Supplementary Table S13), the LLM scored each answer on a five-tier scale: ‘Very likely a ESP (5)’, ‘Likely a ESP (4)’, ‘Insufficient Evidence (3)’, ‘Unlikely to be a ESP (2)’, and ‘Very unlikely to be a ESP (1)’. The model was explicitly instructed to base its scoring solely on the provided text from Evolla, without leveraging its own intrinsic knowledge. For proteins that received the highest score of 5, we converted their corresponding answers into vector representations using a text-embedding model (text-embedding-3-large, provided by OpenAI). These vectors were then subjected to unsupervised clustering using the HDBSCAN algorithm (via the hdbscan.HDBSCAN class with default parameters), which yielded 67 groups, comprising 66 distinct clusters and one group of outliers (designated as −1). For each of the 66 clusters, the collected answers were provided to an LLM (using gemini-3-pro for its supreme performance) to summarize the common characteristics within the cluster’s answers and distill these features into representative keywords (see Supplementary Table S14 for prompt). These clusters were subsequently manually assigned to four major functional categories: Information Storage and Processing (ISP), Cellular Processes and Signaling (CPS), Metabolism (MET), and Others (OTR).

### 7.16 Identification of isomorphic ESPs by eukaryotic enrichment analysis

To identify structural homologs, we queried the protein sequences against the AFDB50 database [86] (foldseek database corresponding to UniRef50 sequence database, downloaded on October 2024) using Foldseek [7] (version 70cea935733e0f70e3e10a0572b45ba6e21bfa2d). The search was executed with the command Foldseek search and the parameter ‘–max-seqs 1000 -e inf –format-output query,target,fident,alnlen,evalue,bits,taxid,taxname’. To establish the background distribution for overrepresentation analysis, we first determined the overall taxonomic composition of the AFDB50 database using the ‘foldseek taxonomyreport’ command. Taxonomic lineage information for each hit was retrieved using ete3.NCBITaxa class with the NCBI Taxonomy database file (taxdump.tar.gz, downloaded on November 21, 2025). To assess eukaryotic enrichment, we analyzed the top 1,000 structural hits for each query protein. We performed a one-tailed Fisher’s exact test (using scipy.stats.fisher exact, alternative=‘greater’) to evaluate the overrepresentation of eukaryotic taxa at the domain level. To account for multiple hypothesis testing, p-values were adjusted using the Bonferroni correction (implemented via statsmodels.stats.multitest.fdrcorrection). Proteins with an adjusted q-value *<* 0.05 were classified as candidate isomorphic ESPs (iESPs)—defined as Asgard proteins exhibiting statistically significant structural similarity to eukaryotic counterparts.

### 7.17 Preparation of the recombinant proteins

The Vps4 coding sequences from *S. cerevisiae*, *Atabeyarchaeum deiterrae*, *Borrarchaeum weybense*, *Hermodarchaeum yapensis* and *Jordiarchaeum madagascariense* were codon optimized for expression in *Escherichia coli* BL21, synthesized and inserted into the pCold-TF vector, respectively. The corresponding gene sequences and other additional information are provided in Supplementary File 4, Table 3.

The *E. coli* BL21 with the recombinant vectors were inoculated in lysogeny broth (LB) medium containing 100 mg ml^−1^ ampicillin and cultivated at 37 °C until the optical density at 600 nm (OD_600_) reached 0.4 to 0.6, and then isopropyl *β*-D-1-thiogalactopyranoside was added at the final concentration of 0.5 mM, followed by cultivation at 15 °C for 16 to 18 h. The cell pellets were harvested and resuspended in 25 ml binding buffer (20 mM Tris, 300 mM NaCl, 1 mM dithiothreitol, pH 7.8), followed by ultrasonic decomposition. The proteins were purified by 1-mL HisTrap HP column (Cytiva, USA), which had been equilibrated with 20 mM Tris, 300 mM NaCl, 20 mM imidazole (pH 7.8). The column was washed with the same buffer containing 50 mM imidazole to elute unwanted protein. The proteins were eluted with the same buffer containing with 500 mM imidazole. Finally, the proteins were exchanged to the reaction system (20 mM HEPES, 100 mM NaCl, 10 mM MgCl_2_, pH 7.4) by using a desalting column (PD-10, Cytiva) and concentrated by ultrafiltration (30 kDa JetSpin, Biofil). Protein purity was confirmed by sodium dodecyl sulfate-polyacrylamide gel electrophoresis (SDS-PAGE) and visualized by Coomassie Blue Super Fast Staining Solution (Beyotime, China). Protein concentrations were determined with the Bradford Protein Assay Kit (Beyotime, China), using bovine serum albumin (BSA) as a standard.

### 7.18 Measurement of ATPase activity

The ATPase activity was measured by a modified malachite green assay [52, 87]. Recombinant protein (4 *µ*M) was incubated in a reaction buffer containing 1 mM ATP, 20 mM HEPES, 100 mM NaCl, 10 mM MgCl_2_, 1 mM dithiothreitol (pH 7.4) in a total volume of 50 *µ*L. The reaction was carried out at either 30 °C or 39 °C for 90 minutes, and then immediately terminated by freezing in liquid nitrogen. To develop color, 100 *µ*L of malachite green reagent (14 mM ammonium molybdate, 1.3 M HCl, and 1.5 mM malachite green) was added to the mixture, followed by 50 *µ*L of 21% (w/v) citric acid. After incubating at room temperature for 30 minutes, the absorbance of the green complex was measured at 660 nm using a microplate reader. The absorbance value, which reflects the amount of free phosphate released by Vps4 ATP hydrolysis, was used as an indicator of ATPase activity. Reactions performed in the absence of enzyme served as negative controls to confirm that the colorimetric signal (shift from gold to green) was specific to Vps4 activity.

### 7.19 *S. cerevisiae* strains and cultivation

The *S. cerevisiae* strain BY4741 (*MATa leu2*Δ*0 met15*Δ*0 ura3*Δ*0 his3*Δ*1*) and its derived *vps4* null mutant strain YPR173Ca (here designated *S. cerevisiae* Δ*vps4*), were obtained from *S. cerevisiae* deletion mutant library [88]. Strains were routinely cultured in YPD medium (10 g L^−1^ yeast extract, 20 g L^−1^ peptone, 20 g L^−1^ glucose) or SD-Ura synthetic dropout medium (Coolaber, China) at 30 °C. Solid media were prepared by adding 2% (w/v) agar to the corresponding liquid medium prior to autoclaving.

### 7.20 Construction of complementary strains and spot growth assay

The Vps4 coding sequences from *S. cerevisiae*, *A. deiterrae*, *B. weybense*, *H. yapensis* and *J. madagascariense* (Supplementary File 4, Table 3) were codon optimized for expression in *S. cerevisiae*. To minimize variation in transcriptional regulation, all heterologous genes were placed under the control of the native *S. cerevisiae vps4* promoter, a 500 bp region upstream of the start codon. Each codon-optimized gene sequence, along with the native promoter and the *CYC1* terminator, was synthesized into the pPOT1-RFP vector [89]. The resulting construct carrying the full-length *S. cerevisiae vps4* gene (including its promoter and the *CYC1* terminator) was introduced into the *S. cerevisiae* Δ*vps4* mutant to generate the complemented strain [90], designated ‘*+Sc*’ in the figures. As controls, both the wild-type BY4741 and the Δ*vps4* mutant were transformed with the empty pPOT1-RFP vector. For phenotypic analysis, all strains were cultured in SD-Ura medium at 30 °C to mid-log phase (OD_600_ between 0.5 and 0.8). Cultures were then diluted to an OD_600_ of 0.1, and a series of 10-fold dilutions were prepared. Five microliters of each dilution were spotted onto SD-Ura solid medium and incubated for subsequent observation.

### 7.21 FM 4-64 staining

Each strain of *S. cerevisiae* was cultured in SD-Ura medium at 30 °C and harvested at mid-log phase (OD_600_ between 0.5 and 0.8). The cells were then stained with 5 *µ*M FM 4-64 (Macklin, China) at 30 °C for 20 minutes, followed by two washes with fresh medium. After washing, the cells were further incubated for 2 hours to allow for dye internalization and vacuolar membrane labeling [91]. Finally, fluorescence imaging was performed using an Eclipse Ti2 microscope (Nikon, Japan). In the quantitative analysis of the class E compartment phenotype, we followed a standardized counting procedure across biological replicates. For each experimental condition, approximately 20 cells were scored from each of three randomly chosen microscopic fields. A cell was counted as positive if it displayed the characteristic aberrant prevacuolar compartment. The percentage of positive cells was calculated per field, and the results from the three fields were averaged to generate the final data for that condition. Statistical comparisons and graphical presentations were based on these averaged values from independent replicates.

### 7.22 Whole-genome sequencing and gene prediction

Building on our prior work [59], the genomes of *Pseudomonas sp.* were sequenced using the Illumina HiSeq 2000 platform (Majorbio Bio-Pharm Technology Co., Ltd, China). Genomic DNA was used to construct paired-end libraries (insert size 300-400 bp), followed by PE150 sequencing. Raw reads were processed using Trimmomatic v0.32 [92] and assembled via SPAdes v3.12.0 [93] (with k-mer parameters 21–121). Subsequently, protein-coding genes were predicted using Prokka v1.14.6 [94], generating a search space of 4,182 predicted genes. The assembled genome sequence has been deposited in the NCBI database under BioProject accession number PRJNA1414862 and BioSample SAMN54915589.

### 7.23 Identification of PET hydrolase by Evolla

We utilized the Evolla-10B-DPO version to screen the complete set of predicted protein-coding genes from the *Pseudomonas sp.* genome for potential plastic-degrading activity. Each protein sequence was individually queried using the specific question: ‘Is this protein possibly related to plastic degradation?’. We implemented a two-step filtering protocol to identify candidates. First, we selected proteins for which the model generated a definitive affirmative response (i.e., starting with ‘Yes’), resulting in an initial pool of 26 candidates. Second, we further refined this selection by searching for the specific keyword ‘PET’ within the generated answers of these candidates to isolate the final target.

### 7.24 Protein expression and purification

Genes encoding PET hydrolases from *Pseudomonas sp.* (*Ps*PETase, full sequence provided in the Supplementary Table S16) and *Ideonella sakaiensis* 201-F6 (*Is*PETase/WT, UniProt accession A0A0K8P6T7) were codon-optimized and synthesized with a C-terminal hexa-histidine (*His*_6_) tag. These constructs were subcloned into the pET-28a(+) vector. To ensure intracellular soluble expression, N-terminal signal peptides were identified using SignalP 5.0 and subsequently removed via an Exnase-based seamless cloning strategy.

The resulting expression plasmids were transformed into *Escherichia coli* BL21(DE3). A 16 mL seed culture was cultivated at 37 °C in LB medium supplemented with 50 µg/mL kanamycin, which was subsequently used to inoculate 1.6 L of the same medium. Cultures were grown at 37 °C with shaking at 220 r.p.m. until an OD_600_ of 0.6 was reached. Protein expression was then induced with 0.1 mM IPTG for 20 h at 16 °C. Cells were harvested by centrifugation (5,000 × g, 30 min, 4 °C), resuspended in lysis buffer (25 mM Tris-HCl, 500 mM NaCl, pH 7.4), and disrupted using a high-pressure homogenizer (KS-2000M, Kaibai Nano). The crude lysate was clarified by centrifugation (12,000 × g, 30 min, 4 °C), and the *His*_6_-tagged proteins were purified from the supernatant using Ni-charged MagBeads (GenScript) following the manufacturer’s protocol. Protein purity was assessed by SDS-PAGE, and concentrations were determined using the BCA toolkit (Sangon).

### 7.25 Preparation of amorphous PET films and degradation assays

Amorphous PET films were fabricated by dissolving commercial amorphous gf-PET film (Goodfellow, UOM Code: 577-529-50; specification: 1.3–1.4 g cm^−3^ density, 1.58–1.64 refractive index, 100 × 10^−13^ cm^3^. cm cm^−2^ s^−1^ Pa^−1^ permeability to water at 25 °C, 20–80 × 10^−6^ K^−1^ coefficient of thermal expansion, 0.13–0.15 W m^−1^ K^−1^ at 23 °C thermal conductivity) in 1,1,1,3,3,3-hexafluoro-2-propanol (HFIP) to a concentration of 20 mg/mL. A 4 mL aliquot of the solution was cast onto a 10-cm diameter flat glass plate. Following overnight solvent evaporation, the resulting films were sterilized in 75% ethanol for 2 h, carefully peeled, and partitioned into 1 × 1 cm^2^ for enzymatic assays.

Temperature-dependent degradation profiles were assessed at 30, 35, 40, and 45 °C. Reactions were performed in 1 mL of 50 mM KH_2_PO_4_-NaOH buffer (pH 8.0) containing 500 nM enzyme and incubated for 24 h.

The concentrations of PET monomers (TPA and MHET) released during enzymatic depolymerization were quantified using a Thermo Scientific Vanquish HPLC system (Thermo Fisher Scientific) equipped with a Shim-pack GIST C18-AQ column (4.6×150 mm, 3 *µ*m; Shimadzu). Before analysis, samples were filtered through 0.2-*µ*M nylon syringe filters (JIN TENG). The mobile phase consisted of 0.1% (v/v) trifluoroacetic acid (TFA) in water (A) and 0.1% (v/v) TFA in acetonitrile (B). The column temperature was maintained at 25 °C, and the flow rate was fixed at 0.8 mL min^−1^. The UV detection wavelength was set to 260 nm. The gradient elution program was optimized as follows: 0–3 min, 5% B (isocratic); 3–6 min, 5–20% B; 6–11 min, 20–25% B; 11–19 min, 25–95% B; 19–23 min, 95% B (column wash); 23–25 min, 95–5% B; and 25–32 min, 5% B (re-equilibration). Monomers were identified by comparing retention times with those of commercial TPA (*>*99.0%, MACKLIN) and MHET (*>*98.0%, MACKLIN) standards.

For long-term visualization of film morphology change, degradation was performed in 2 mL of the same buffer (pH 8.0) with 1,000 nM enzyme at 45 °C over a 7-day period. The reaction medium was replenished every 24 h with fresh buffer containing 1,000 nM of the respective enzyme.

### 7.26 Protein thermal stability measured by nanoDSF

The thermal stability of purified proteins was characterized using nano-differential scanning fluorimetry (nanoDSF) on a Prometheus Panta instrument (NanoTemper Technologies). Samples (1 mg/mL) were loaded into chemically inert high-sensitivity glass capillaries (NanoTemper, Cat. #K003). The temperature was increased from 25 °C to 80 °C at a linear ramp rate of 1 °C /min. Intrinsic tryptophan/tyrosine fluorescence was monitored at 330 nm and 350 nm. The melting temperature (*T_m_*) was determined by identifying the peak of the first derivative of the fluorescence ratio (*F*_350_*/F*_330_) relative to temperature.

### 7.27 *p*-Nitrophenyl Butyrate (*p*-NPB) assay

Esterase activity was determined by monitoring the continuous hydrolytic release of *p*-nitrophenol (*p*-NP) from *p*-nitrophenyl butyrate (*p*-NPB). Reactions were performed in a 100 *µ*L system consisting of 50 mM KH_2_PO_4_-NaOH buffer (pH 8.0), 200 nM enzyme, and 1 mM *p*-NPB. The substrate was introduced from a 100 mM stock solution in dimethyl sulfoxide (DMSO).

The reaction progress was monitored at 37 °C by recording the absorbance at 405 nm every 20 s for a total of 600 s using a Tecan microplate reader. Absorbance values were converted to *p*-NP concentrations (mM) using a pre-established *p*-NP standard curve (*y* = 4.3047*x* + 0.212). All data were baseline-normalized by subtracting the initial concentration at *t* = 0 to ensure all progress curves originated from the origin. The initial velocity (*v*_0_) was calculated by linear regression of the progress curves during the initial linear phase (0–250 s). The net *v*_0_ was determined by subtracting the spontaneous hydrolysis rate of the blank control (buffer only) from the enzymatic reaction rate. Data are presented as mean ± s.d. from three independent experiments (*n* = 3).

### 7.28 Phylogenetic analysis

To delineate the evolutionary relationships of the identified candidates, a representative dataset was compiled comprising 137 PET hydrolases with experimentally validated PET-degrading activity curated from extant literature (Supplementary File 5). N-terminal signal peptides were predicted via SignalP 5.0 [60] and subsequently removed to ensure the alignment of mature catalytic domains. Mature sequences were aligned via MAFFT v7.526 (L-INS-i) [95] and trimmed using trimAl v1.4 (-automated1) [96]. The maximum-likelihood phylogeny was reconstructed with IQ-TREE v2.2.2.6 [97] using the BIC-selected LG+F+I+R5 model and 1,000 ultrafast bootstrap replicates. The resulting phylogeny was rooted using homologous cutinases from Fusarium species as the outgroup, and the final tree was visualized via iTOL v6 [98].

## Data availability

The Evolla-10B weights can be downloaded from https://huggingface.co/westlake-repl/Evolla-10B. The Evolla-80B weights can be downloaded from https://huggingface.co/westlake-repl/Evolla-80B. The Evolla-10B-DPO weights can be downloaded from https://huggingface.co/westlake-repl/Evolla-10B-DPO. The SaProt 650M and 1.3B weights can be downloaded from https://huggingface.co/westlake-repl/SaProt 650M AF2 and https://huggingface.co/westlake-repl/SaProt 1.3B AF2, respectively. The LLM decoders for Evolla-10B and Evolla-80B can be downloaded from https://huggingface.co/meta-llama/Meta-Llama-3-8B-Instruct and https://huggingface.co/meta-llama/Llama-3.1-70B-Instruct, respectively.

## Code availability

Evolla is open-sourced under the MIT license. The code repository is available at https://github.com/westlake-repl/ Evolla. The Evolla webserver is located at http://www.chat-protein.com/.

## Acknowledgement

We are grateful to the Zhipu BigModel Open Platform for their support in data generation. We thank Nan Li and the Westlake University HPC Center for providing the essential computing resources. We thank Prof. Dr. Junbiao Dai from the Agricultural Genomics Institute at Shenzhen, Chinese Academy of Agricultural Sciences, and Dr. Shuangying Jiang from the Shenzhen Institute of Advanced Technology, Chinese Academy of Sciences, for providing the yeast BY4741 strain and the YPR173Ca mutant. We also extend our sincere gratitude to Igor Tolstoy, Sergey Ovchinnikov, Christine Orengo, Maria Martin, Vishal Joshi, Weining Lin, Haiyang Cui, Shuping Xu, Qi Hu, Jingwei Xu, Florian Wollweber, Xinyu Huang, Ying Zhen, Rongao Kou, Jin Sun, Wenhao Chen, Jiahao Mei, Ruoyu Zhou, Shurui Ning, Lei Hu, Huijie Mi, Huijiao Yang, and Shunyi Yang for their valuable discussions and insights.

## Appendix A Appendix

**Supplementary Table S1:** The 56 sections collected from the UniProt database include both protein-level and residue-level annotations.

| section type | sections |
| --- | --- |
| protein level | Function, Miscellaneous, Caution, Catalytic activity, Cofactor, Activity regulation, Biophysicochemical properties, Pathway, Involvement in disease, Allergenic properties, Toxic dose, Pharmaceutical use, Disruption phenotype, Subcellular location, Post-translational modification, Subunit, Domain (non-positional annotation), Sequence similarities, RNA Editing, Tissue specificity, Developmental stage, Induction, Biotechnology, Polymorphism, GO annotation, Proteomes, Protein names, Gene names, Organism, Taxonomic lineage, Virus host |
| residue level | Active site, Binding site, Site, DNA binding, Natural variant, Mutagenesis, Transmembrane, Topological domain, Intramembrane, Signal peptide, Propeptide, Transit peptide, Chain, Peptide, Modified residue, Lipidation, Glycosylation, Disulfide bond, Cross-link, Domain, Repeat, Compositional bias, Region, Coiled coil, Motif |

**Supplementary Table S2:** Prompt for generation of persona-oriented questions. Detailed role information and examples of IUs described here are in Supplementary Table S4 and Supplementary File 1, respectfully.

|  |
| --- |
| Prompt |
| You receive a few sentences, each describing the same protein you are analysing.<br>In addition, specific amino acid features within the protein are given, along with detailed spans or specific amino acid sites.<br>The spans are in the form of tuples, represented as (x, y) with integer numbers ranging from 0 to the length of the protein.<br>These values correspond to the start amino acid index x and the end amino acid index y.<br>The sites are in the form of tuples, represented as (A, x, B), where A means the original amino acid, x means the index of the site and B means the mutated amino acid.<br>If A and B are the same amino acid, some important note will be in the site. The single amino acids are represented as (x, ), where x means the index of the amino acid.<br>You will play three different roles, and based on the provided protein description, each role will ask a question. Make sure the questions do not overlap and are asked from the perspective of each role.<br>#Role 1#: {0}<br>#Role 2#: {1}<br>#Role 3#: {2}<br>The generated questions should be formatted as follows:<br>Question 1: [Question from Role 1]<br>Question 2: [Question from Role 2]<br>Question 3: [Question from Role 3]<br>When using the information from the caption and sites information, directly analyse the protein, and do not mention that the information source is the caption or the sites. Don't include the name of the protein, refer it as "this protein"<br>'#The Given Description#', 'given description', 'description', 'detailed information' are not allowed to appear in #Question#.<br>#Here is the Given Description#:<br>{}<br>#Questions#: |

**Supplementary Table S3:**
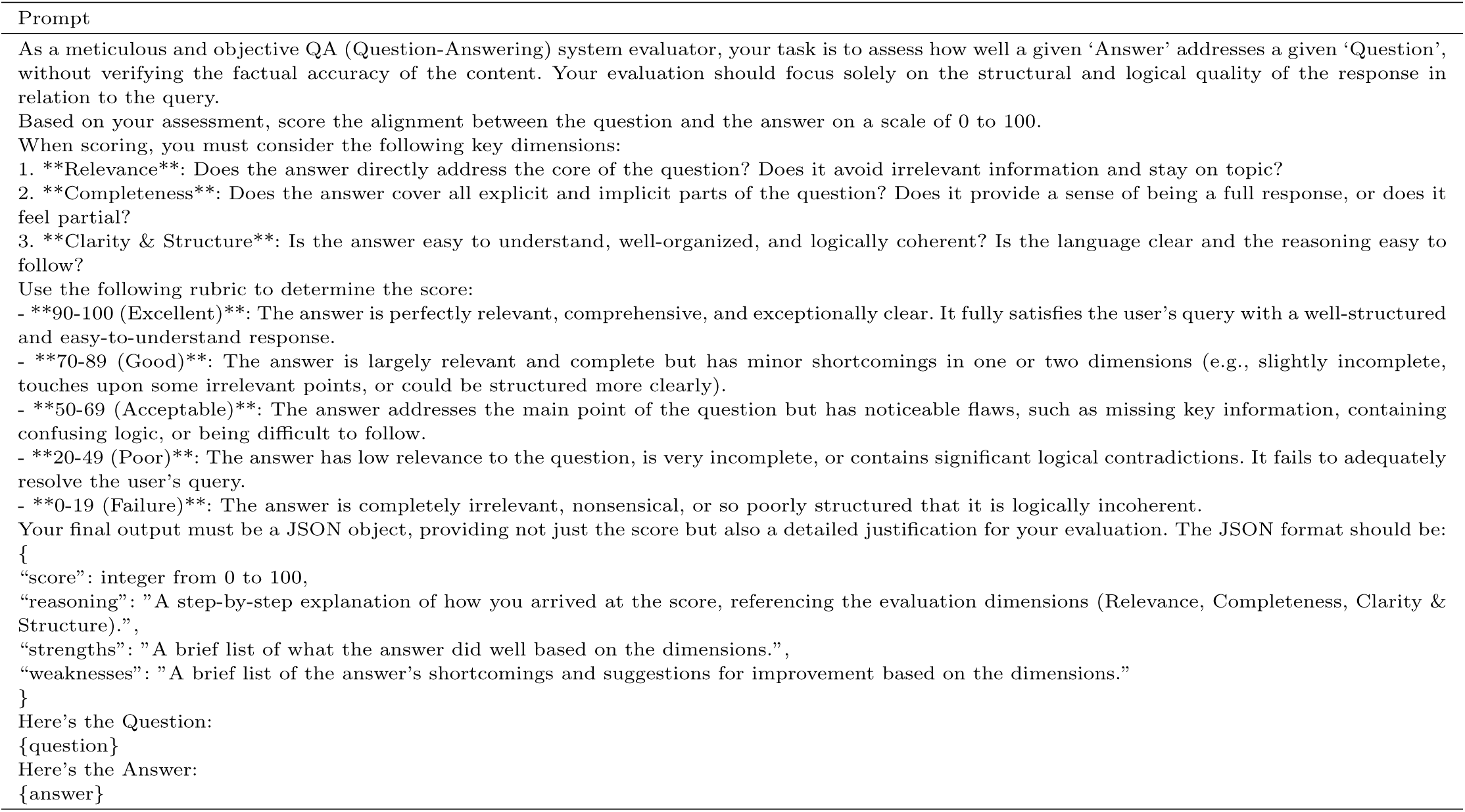
Prompt for evaluation of question-answer relevance.

|  |
| --- |
| Prompt |
| As a meticulous and objective QA (Question-Answering) system evaluator, your task is to assess how well a given 'Answer' addresses a given 'Question', without verifying the factual accuracy of the content. Your evaluation should focus solely on the structural and logical quality of the response in relation to the query.<br>Based on your assessment, score the alignment between the question and the answer on a scale of 0 to 100.<br>When scoring, you must consider the following key dimensions:<br>**Relevance**: Does the answer directly address the core of the question? Does it avoid irrelevant information and stay on topic?<br>**Completeness**: Does the answer cover all explicit and implicit parts of the question? Does it provide a sense of being a full response, or does it feel partial?<br>**Clarity & Structure**: Is the answer easy to understand, well-organized, and logically coherent? Is the language clear and the reasoning easy to follow?<br>Use the following rubric to determine the score:<br>**90-100 (Excellent)**: The answer is perfectly relevant, comprehensive, and exceptionally clear. It fully satisfies the user's query with a well-structured and easy-to-understand response.<br>**70-89 (Good)**: The answer is largely relevant and complete but has minor shortcomings in one or two dimensions (e.g., slightly incomplete, touches upon some irrelevant points, or could be structured more clearly).<br>**50-69 (Acceptable)**: The answer addresses the main point of the question but has noticeable flaws, such as missing key information, containing confusing logic, or being difficult to follow.<br>**20-49 (Poor)**: The answer has low relevance to the question, is very incomplete, or contains significant logical contradictions. It fails to adequately resolve the user's query.<br>**0-19 (Failure)**: The answer is completely irrelevant, nonsensical, or so poorly structured that it is logically incoherent.<br>Your final output must be a JSON object, providing not just the score but also a detailed justification for your evaluation. The JSON format should be:<br>`{ "score": integer from 0 to 100, "reasoning": "A step-by-step explanation of how you arrived at the score, referencing the evaluation dimensions (Relevance, Completeness, Clarity & Structure).", "strengths": "A brief list of what the answer did well based on the dimensions.", "weaknesses": "A brief list of the answer's shortcomings and suggestions for improvement based on the dimensions." }`<br>Here's the Question:<br>`{question}`<br>Here's the Answer:<br>`{answer}` |

**Supplementary Table S4:** The description of Personas.

| Personas | description |
| --- | --- |
| clinical biochemist | You are a clinical biochemist who analyzes proteins in biological fluids to diagnose and monitor diseases. Your questions focus on the clinical significance of protein markers, the development of diagnostic assays, and the interpretation of protein profiles in patient samples. |
| high school student | You are a high school student with a budding interest in biology. Your questions are basic and seek to understand the fundamental concepts of proteins, such as their role in the body, how they are made, and why they are important for health. |
| undergraduate biology student | You are an undergraduate biology student learning about the structure and function of proteins. Your questions reflect your coursework, including inquiries about protein synthesis, the relationship between amino acids and protein structure, and the basics of protein folding. |
| postdoctoral fellow | You are a postdoctoral fellow specializing in a particular aspect of protein science. Your questions are highly specialized and may involve complex experimental designs, cutting-edge technologies, and the integration of multiple research disciplines. |
| medical student | You are a medical student learning about the clinical relevance of proteins. Your questions are about the role of proteins in disease, how protein abnormalities can lead to medical conditions, and the potential for protein-based treatments in healthcare. |
| structural biologist | You are a structural biologist using techniques like X-ray crystallography and cryo-EM to determine protein structures. Your questions are about the relationship between protein structure and function, the dynamics of protein folding, and the implications of structural data for drug design. |
| geneticist | You are a geneticist examining the genetic basis of protein function and variation. Your questions explore the inheritance patterns of protein-coding genes, the effects of genetic polymorphisms on protein activity, and the role of proteins in genetic disorders. |
| synthetic biologist | You are a synthetic biologist designing and engineering proteins for specific applications. Your questions are about protein optimization, the creation of novel protein functions, and the use of synthetic proteins in biotechnology and medicine. |
| science communicator | You are a science communicator who translates complex protein research for the general public. Your questions are about the best ways to explain protein science in an accessible manner, the use of analogies and metaphors, and the impact of public understanding on scientific progress. |
| middle school student | You are a middle school student with a curiosity about the natural world. Your questions are very simple and direct, focusing on the basic properties of proteins, how they are related to DNA, and why they are important for living organisms. |

**Supplementary Table S5:** Question evolution strategy and the corresponding prompts.

| Evolutions | Prompts |
| --- | --- |
| Adding constraints | Please add one more constraints/requirements into #The Given Prompt# |
| Deepening and expanding | If #The Given Prompt# contains inquiries about certain issues, the depth and breadth of the inquiry can be increased. |
| Concept specification | Please replace general concepts with more specific concepts. |
| Multi-step reasoning | If #The Given Prompt# can be solved with just a few simple thinking processes, you can rewrite it to explicitly request multiple-step reasoning. |
| Generating rare prompts | Please draw inspiration from the #Given Prompt# to create a brand new prompt. This new prompt should belong to the same domain as the #Given Prompt# but be even more rare. The LENGTH and complexity of the #Created Prompt# should be similar to that of the #Given Prompt#. |

**Supplementary Table S6:**
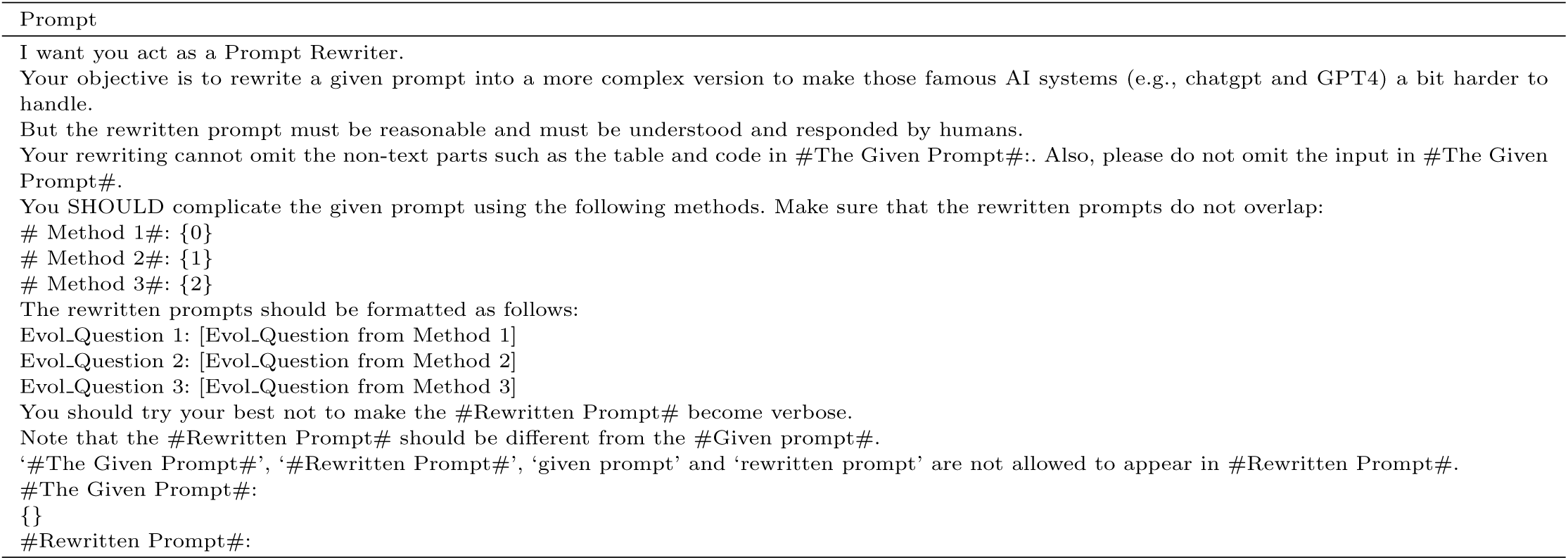
Prompt to evolve the question without providing any protein related information.

**Supplementary Table S7:** Prompt to generate answers with IUs and questions. Examples of IUs described here are in Supplementary File 1.

| Prompt |
| --- |
| You are an AI biology assistant that can analyze a single protein. You receive a few sentences, each describing the same protein you are analysing. In addition, specific amino acid features within the protein are given, along with detailed spans or specific amino acid sites. The spans are in the form of tuples, represented as (x, y) with integer numbers ranging from 0 to the length of the protein. These values correspond to the start amino acid index x and the end amino acid index y. The sites are in the form of tuples, represented as (A, x, B), where A means the original amino acid, x means the index of the site and B means the mutated amino acid. If A and B are the same amino acid, some important note will be in the site. The single amino acids are represented as (x, ), where x means the index of the amino acid.<br>Using the provided caption and sites information, answer the question related to this protein in a detailed manner.<br>Instead of directly mentioning the sites information, utilize this data to analyse the protein using natural language. Include details like site functions, relation to other proteins, contributions of sites to functions.<br>When using the information from the caption and sites information, directly analyse the protein, and do not mention that the information source is the caption or the sites. Don't include the name of the protein, refer it as "this protein". Always answer as if you are directly analysing the protein.<br>When the question cannot be answered based on the information, don't mention the information source is the caption or the sites. These are the sentences about the protein.<br>{Information units}<br>{Question} |

**Supplementary Table S8:**
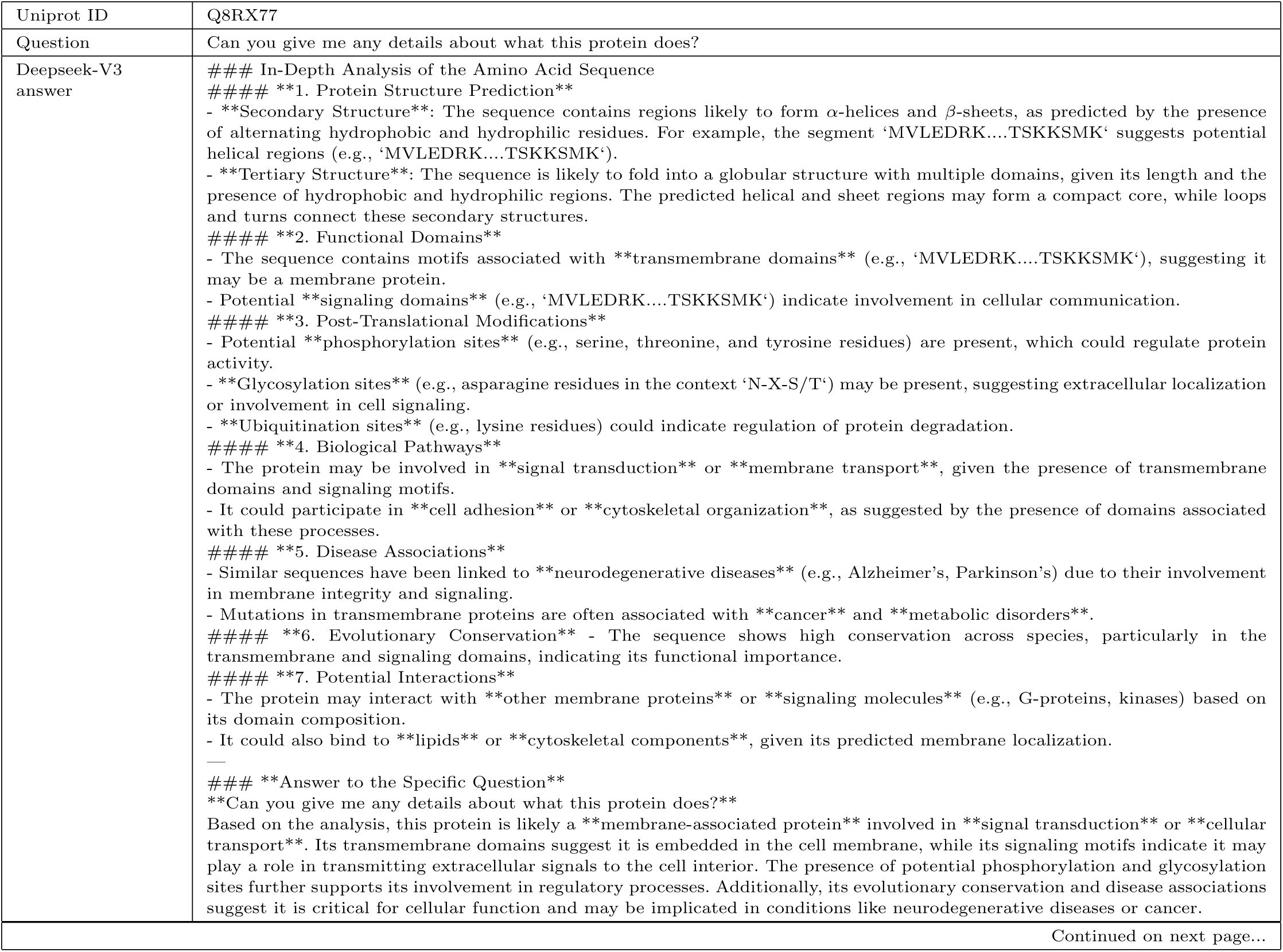
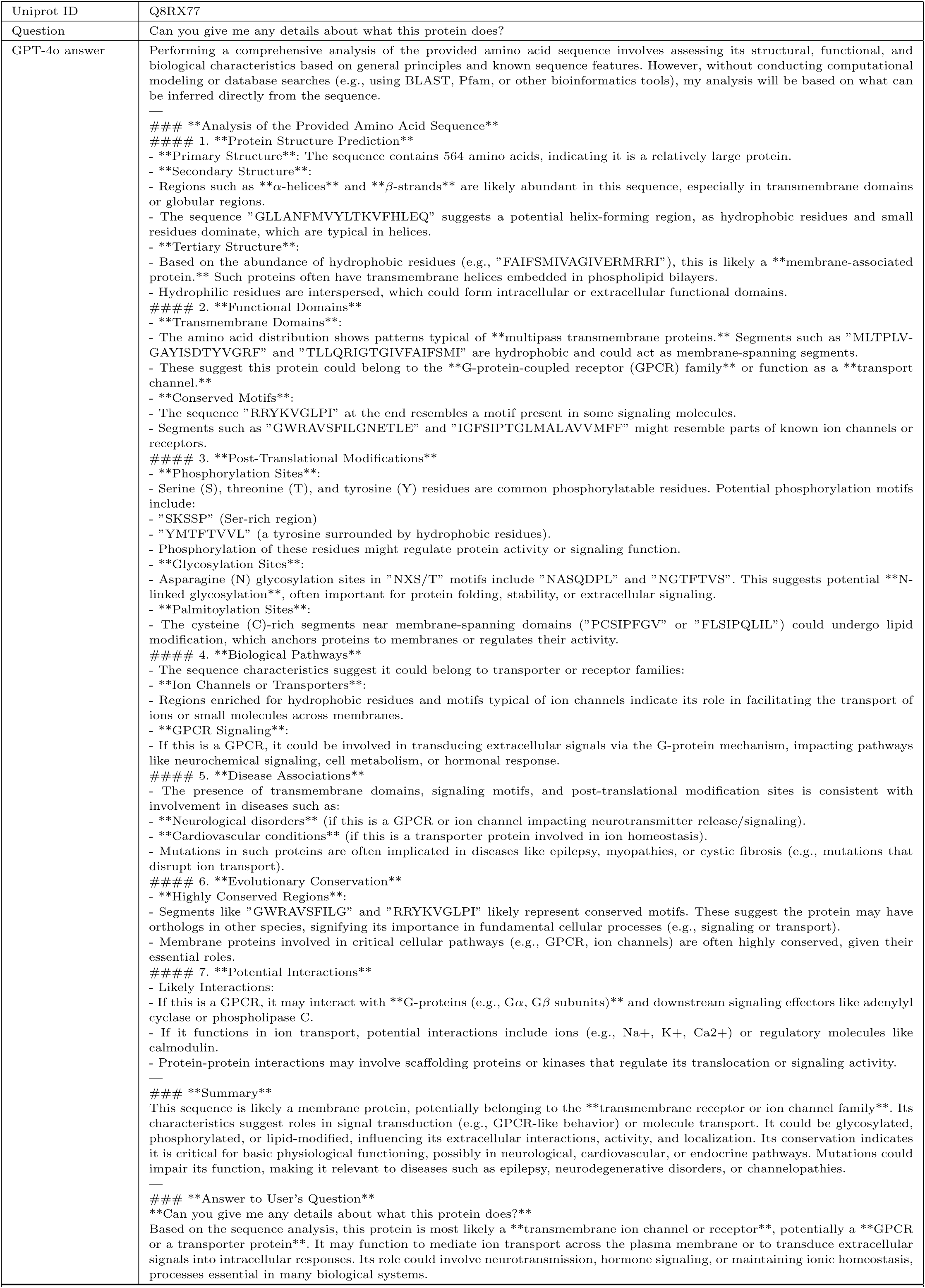
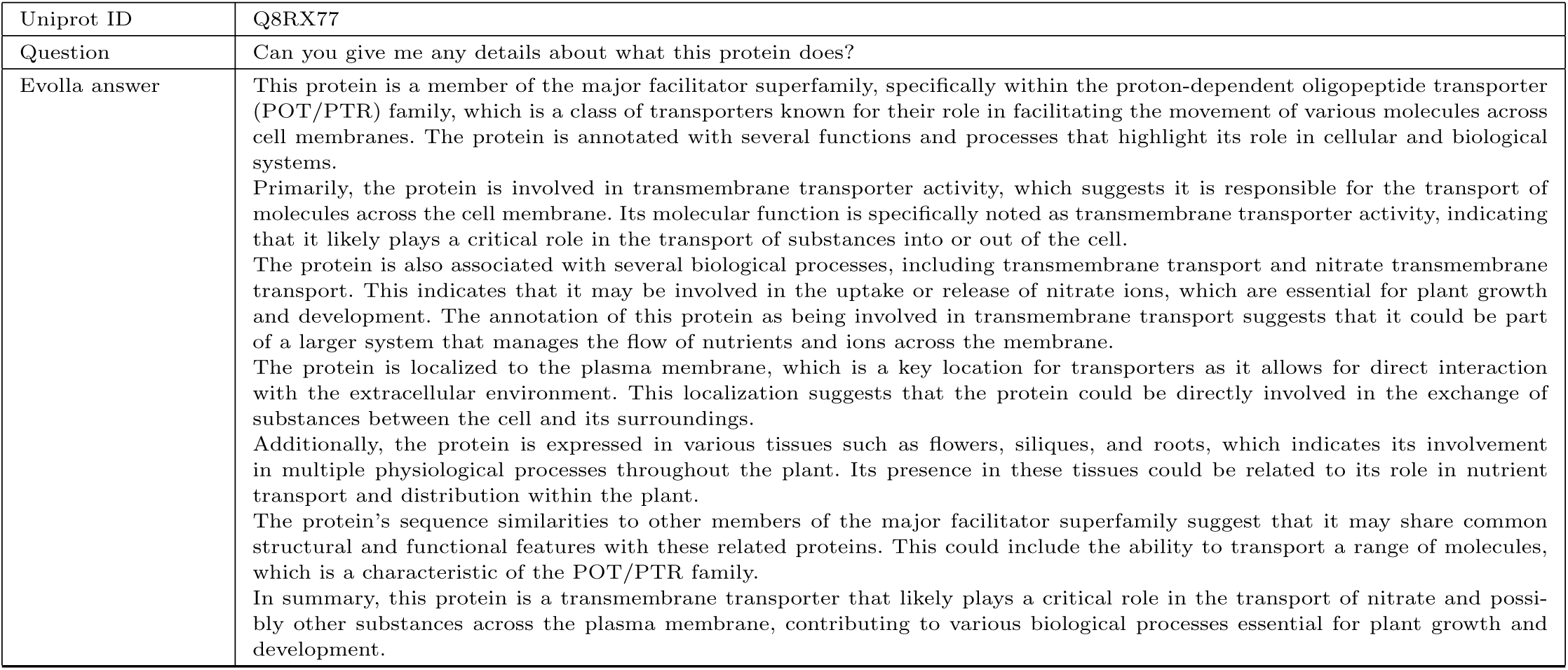
Example outputs of general-purpose LLMs.

**Supplementary Table S9:** Example outputs from specialized protein-language models.

|  |  |
| --- | --- |
| Uniprot ID | B4T945 |
| Question | Which enzymatic operation does this protein specifically carry out? |
| ChatNT answer | The recorded enzymatic activity in this protein is 51.32. |
| InstructBioMol answer | Catalase. |
| Evolla answer | This protein is specifically involved in the enzymatic operation of glycyl-tRNA aminoacylation. It functions as a glycine-tRNA ligase, which is a critical step in protein synthesis. The protein catalyzes the reaction where ATP, glycine, and tRNA(Gly) are converted into AMP, diphosphate, and glycyl-tRNA(Gly). This process is essential for the accurate translation of the genetic code into proteins, as it links the amino acid glycine to its corresponding transfer RNA (tRNA) molecule, ensuring that the correct amino acid is incorporated into the growing polypeptide chain during protein synthesis. |
| Uniprot ID | Q8RX77 |
| Question | Can you give me any details about what this protein does? |
| ChatNT answer | The measured protein expression in this cell is recorded as 1.20. |
| InstructBioMol answer | <p>H <p>R <p>G <p>S <p>M <p>D <p>T <p>A <p>D <p>S <p>S <p>P <p>N <p>V <p>F <p>Q<br><p>E <p>N <p>P <p>G <p>E <p>S <p>R <p>I <p>E <p>E <p>P <p>A <p>S <p>E <p>P <p>A<br><p>K <p>P <p>P <p>P <p>F <p>E <p>Q <p>R <p>A <p>K <p>K <p>P <p>A <p>K <p>S <p>R<br><p>Q <p>S <p>N <p>E 7 <p>A <p>E <p>N <p>A <p>G </FASTA> Protein NRT1/ PTR1. |
| Evolla answer | This protein is a member of the major facilitator superfamily, specifically within the proton-dependent oligopeptide transporter (POT/PTR) family, which is a class of transporters known for their role in facilitating the movement of various molecules across cell membranes. The protein is annotated with several functions and processes that highlight its role in cellular and biological systems.<br>Primarily, the protein is involved in transmembrane transporter activity, which suggests it is responsible for the transport of molecules across the cell membrane. Its molecular function is specifically noted as transmembrane transporter activity, indicating that it likely plays a critical role in the transport of substances into or out of the cell.<br>The protein is also associated with several biological processes, including transmembrane transport and nitrate transmembrane transport. This indicates that it may be involved in the uptake or release of nitrate ions, which are essential for plant growth and development. The annotation of this protein as being involved in transmembrane transport suggests that it could be part of a larger system that manages the flow of nutrients and ions across the membrane.<br>The protein is localized to the plasma membrane, which is a key location for transporters as it allows for direct interaction with the extracellular environment. This localization suggests that the protein could be directly involved in the exchange of substances between the cell and its surroundings.<br>Additionally, the protein is expressed in various tissues such as flowers, siliques, and roots, which indicates its involvement in multiple physiological processes throughout the plant. Its presence in these tissues could be related to its role in nutrient transport and distribution within the plant.<br>The protein's sequence similarities to other members of the major facilitator superfamily suggest that it may share common structural and functional features with these related proteins. This could include the ability to transport a range of molecules, which is a characteristic of the POT/PTR family.<br>In summary, this protein is a transmembrane transporter that likely plays a critical role in the transport of nitrate and possibly other substances across the plasma membrane, contributing to various biological processes essential for plant growth and development. |

**Supplementary Table S10:** Prompt used to calculate the GPT scores.

| Prompt |
| --- |
| As an expert biologist, you are assigned to check whether a paragraph is aligned with facts or not. You will receive some facts, and one paragraph. Score the paragraph between 0 to 100. Give reasons for those aligned ones and not aligned ones.<br>The score should be the format of { "score": score }<br>Here's the facts:<br>{Information units}<br>Here's the paragraph:<br>{paragraph} |

**Supplementary Table S11:** Prompt used for EC number description generation.

| Prompt |
| --- |
| Summarize the following text using the specified template format. Ensure the output is formatted exactly as shown in the template. Only output the text and must start with the EC number.<br>Template format:<br>EC 1.4.3.4: The enzyme, Monoamine oxidase, is an oxidoreductase that catalyzes the oxidative deamination of monoamines, which are neurotransmitters and biogenic amines. It utilizes oxygen as an electron acceptor to convert primary, secondary, and some tertiary amines into aldehydes, ammonia, and hydrogen peroxide. The enzyme is a flavoprotein found in the outer mitochondrial membrane and plays a significant role in the metabolism of neurotransmitters such as serotonin, dopamine, and norepinephrine. It differs from other amine oxidases by its ability to oxidize secondary and tertiary amines. Monoamine oxidase is inhibited by specific compounds like clorgyline and pargyline, which makes it distinct from similar enzymes like diamine oxidase.<br>Text information to be summarized:<br>[detailed information of EC number] |

**Supplementary Table S12:**
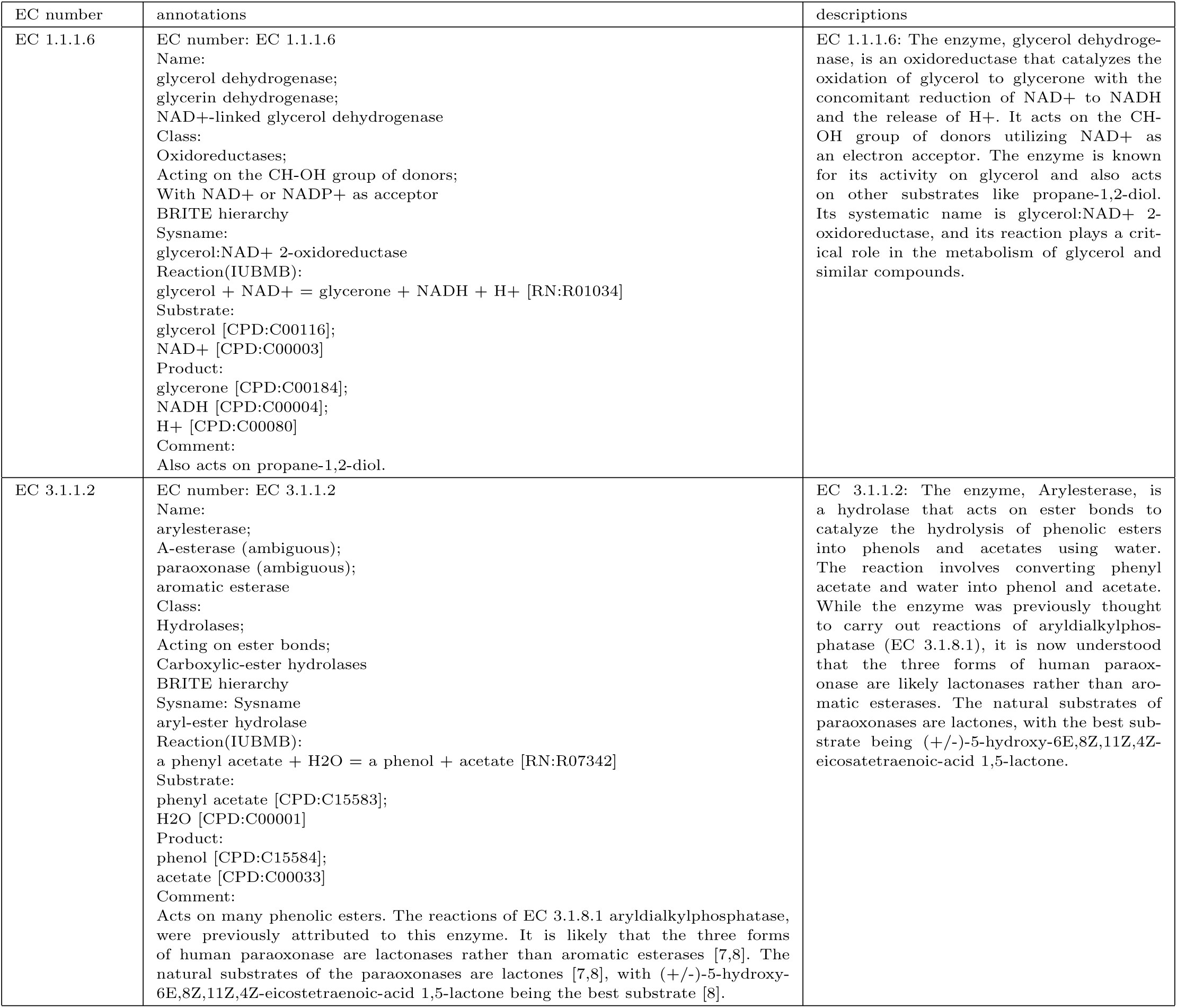
Examples of EC number, their annotations and descriptions.

**Supplementary Table S13:**
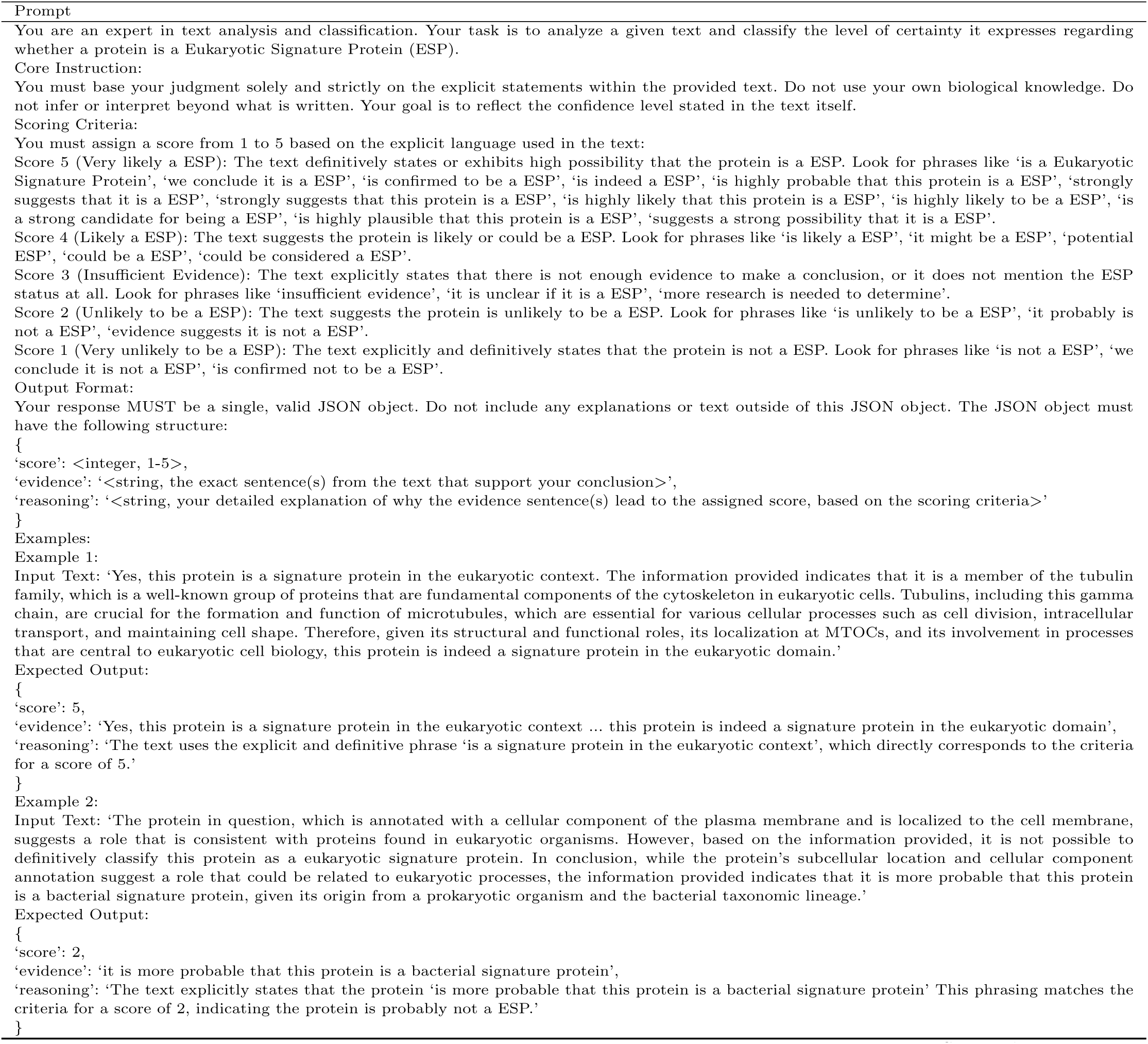
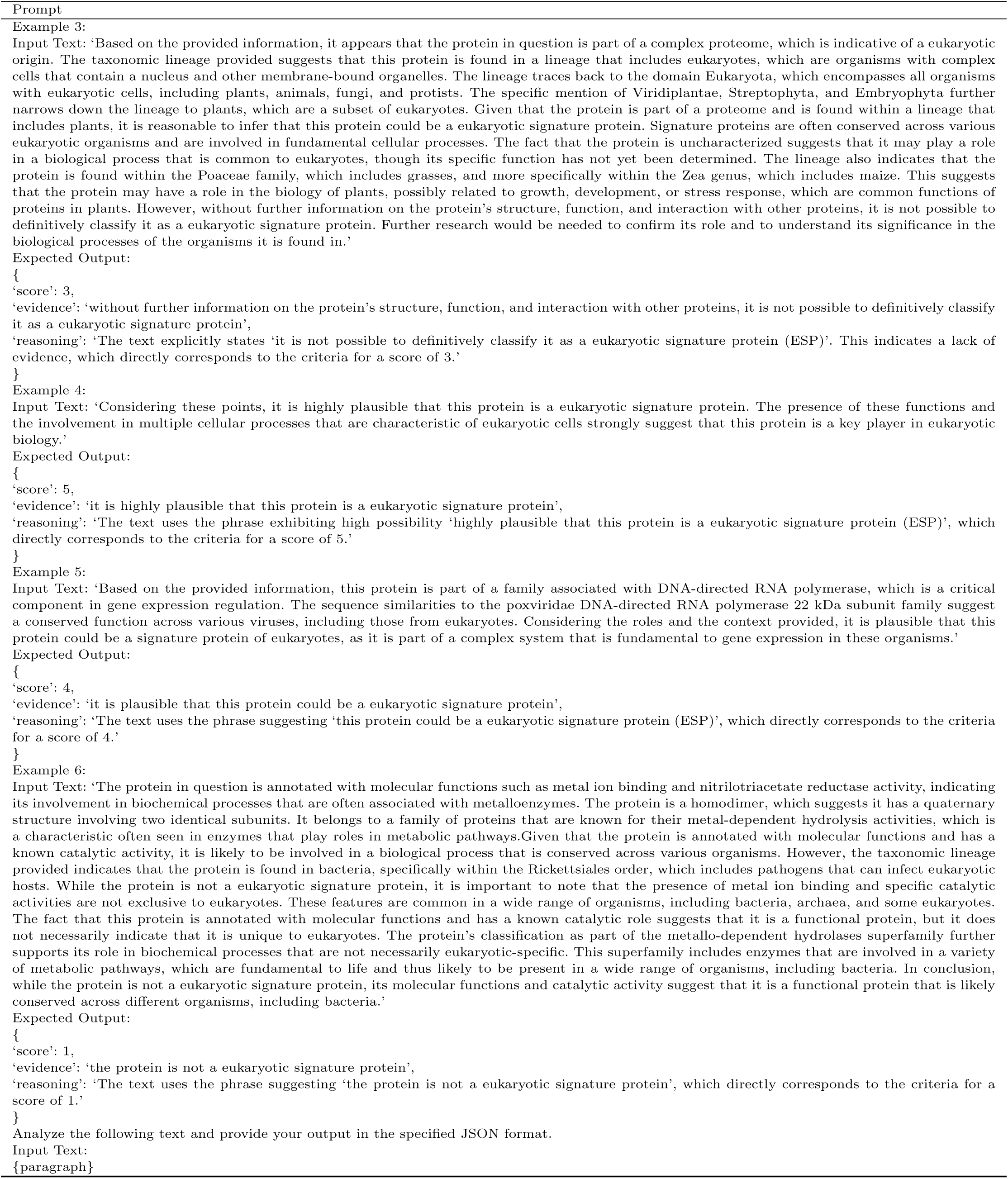
Prompt for evaluating if the response is related to ESP or not.

**Supplementary Table S14:** The prompt used for summarizing common characteristics and extracting keywords for identified Eukaryotic Signature Protein (ESP) clusters.

| Prompt |
| --- |
| You are a specialized API endpoint that processes biochemical data. Your sole function is to receive a list of protein functional descriptions and return a single, valid JSON object.<br>**\*\*Task:\*\***<br>Analyze the provided list of protein functions. Synthesize this information to generate:<br>1. A detailed summary of the collective function.<br>2. A single, concise sentence that acts as a high-level functional headline.<br>**\*\*Input Data:\*\***<br>The input will be a list of strings, where each string describes a protein's function.<br>**\*\*Output JSON Schema:\*\***<br>Your response MUST be a single, valid JSON object and nothing else. Do not wrap the JSON in markdown code blocks or add any explanatory text before or after it. The JSON object must conform to the following structure:<br>`{ 'detailed_summary': 'string', 'concise_functional_headline': 'string' }`<br>- 'detailed_summary': (String) A comprehensive paragraph that integrates the information from all input descriptions, explaining the shared function and mentioning any specific nuances.<br>- 'concise_functional_headline': (String) Use several very short English bullet-point sentences (use ; to separate each sentence) to describe the shared properties of these proteins. Make your summary so clear that a user can immediately understand what kind of proteins these are.<br>Example:<br>**\*\*Process the following input:\*\***<br>'Based on the provided information, this protein is indeed a signature of eukaryotic organisms. The classification within the small heat shock protein (HSP20) family, which is known for its role in protein folding and chaperone activity, is a strong indicator of its eukaryotic origin. The GO annotation specifying its molecular function as a protein folding chaperone and its involvement in unfolded protein binding further supports this classification. Additionally, the protein's association with the hspX gene, which is a common gene in eukaryotes for encoding heat shock proteins, reinforces its eukaryotic nature. The protein's induction by heat shock, increased salt concentration, and starvation, which are stress responses common in eukaryotes, also aligns with the characteristics of a eukaryotic protein. Furthermore, the protein's role in the cytoplasm, as indicated by its subcellular location, is consistent with the typical localization of proteins in eukaryotic cells. The protein's sequence similarities to other known HSP20 family members and its association with specific protein names such as "Heat shock 22 kDa protein, mitochondrial" and "Alpha-crystallin" further confirm its eukaryotic identity. These names suggest that the protein is involved in heat shock responses and may have a role in maintaining protein homeostasis within the cell, which is a characteristic feature of eukaryotic proteins.'<br>'The protein in question, which is a member of the heat shock protein family, is indeed a eukaryotic signature protein. Heat shock proteins are known to be highly conserved across various eukaryotic organisms, which suggests that they play fundamental roles in cellular stress response and protein homeostasis. The presence of this protein in the cytoplasm, as indicated by its subcellular location, is consistent with the role of heat shock proteins in managing the intracellular environment. The fact that this protein is induced by heat shock and has a molecular function of ATP binding further supports its role in cellular processes that are critical for survival under stress conditions. ATP binding is a common feature among proteins involved in energy metabolism and cellular responses to stress, indicating that this protein could be directly involved in the cellular response to heat stress. The sequence similarities to other proteins within the heat shock protein family suggest that this protein shares common structural and functional features with other members of the family. This is consistent with the role of heat shock proteins as molecular chaperones that assist in the proper folding of newly synthesized proteins and the protection of existing proteins from denaturation under stress. The taxonomic lineage of the protein, tracing back to Eukaryota and through various phyla and classes, underscores the evolutionary conservation of the protein family. This lineage is indicative of the essential nature of heat shock proteins in the cellular life cycle of eukaryotes. The gene names provided, such as 62 and 49.1, are typical identifiers for genes encoding heat shock proteins. The gene symbol 49.1, in particular, is often used to denote a specific member of the heat shock protein family, suggesting that this protein is a well-characterized and studied member of the group. In summary, based on its molecular function, subcellular location, sequence similarities, and evolutionary conservation, this protein is a clear eukaryotic signature protein, likely involved in critical cellular processes such as protein folding, stress response, and ATP metabolism.'<br>**\*\*Output:\*\***<br>`{ 'detailed_summary': 'The protein is definitively identified as a conserved eukaryotic signature protein, belonging to the heat shock protein family and more specifically to the small HSP (HSP20) family, which also includes related proteins like alpha-crystallins. Primarily localized in the cytoplasm, it functions as an ATP-binding molecular chaperone. This function is critical for cellular stress response and maintaining protein homeostasis, where it assists in the proper folding of newly synthesized proteins, binds to unfolded proteins, and protects existing proteins from denaturation. Its expression is induced by a range of cellular stressors, including heat shock, increased salt concentration, and starvation. This classification is strongly supported by its deep taxonomic lineage tracing back to Eukaryota, its association with characteristic HSP genes (e.g., hspX, 49.1), and sequence similarities to other members of the heat shock protein family. The combination of its molecular function, subcellular location, stress-induced expression, and evolutionary conservation firmly establishes it as an essential eukaryotic protein.', 'concise_functional_headline': 'Eukaryotic Heat Shock Protein (HSP); ATP-Binding Molecular Chaperone; Stress Response & Protein Homeostasis' }`<br>**\*\*Process the following input:\*\***<br>`{paragraph}`<br>**\*\*Output:\*\*** |

**Supplementary Table S15:**
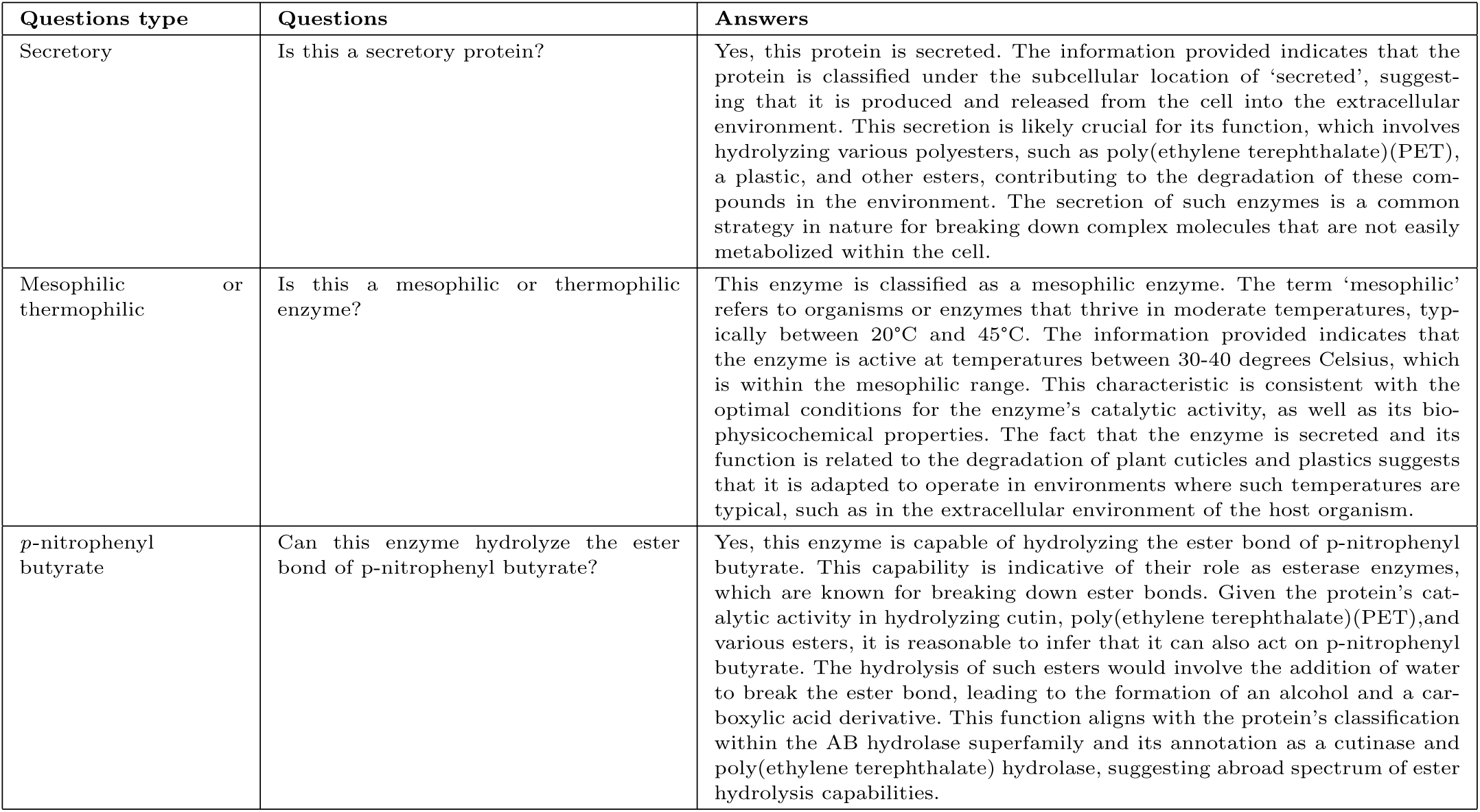
Questions and Evolla’s answers used to specify the properties of the *Ps*PETase.

| Questions type | Questions | Answers |
| --- | --- | --- |
| Secretory | Is this a secretory protein? | Yes, this protein is secreted. The information provided indicates that the protein is classified under the subcellular location of ‘secreted’, suggesting that it is produced and released from the cell into the extracellular environment. This secretion is likely crucial for its function, which involves hydrolyzing various polyesters, such as poly(ethylene terephthalate)(PET), a plastic, and other esters, contributing to the degradation of these compounds in the environment. The secretion of such enzymes is a common strategy in nature for breaking down complex molecules that are not easily metabolized within the cell. |
| Mesophilic or thermophilic | Is this a mesophilic or thermophilic enzyme? | This enzyme is classified as a mesophilic enzyme. The term ‘mesophilic’ refers to organisms or enzymes that thrive in moderate temperatures, typically between 20°C and 45°C. The information provided indicates that the enzyme is active at temperatures between 30-40 degrees Celsius, which is within the mesophilic range. This characteristic is consistent with the optimal conditions for the enzyme’s catalytic activity, as well as its biophysicochemical properties. The fact that the enzyme is secreted and its function is related to the degradation of plant cuticles and plastics suggests that it is adapted to operate in environments where such temperatures are typical, such as in the extracellular environment of the host organism. |
| *p*-nitrophenyl butyrate | Can this enzyme hydrolyze the ester bond of *p*-nitrophenyl butyrate? | Yes, this enzyme is capable of hydrolyzing the ester bond of *p*-nitrophenyl butyrate. This capability is indicative of their role as esterase enzymes, which are known for breaking down ester bonds. Given the protein’s catalytic activity in hydrolyzing cutin, poly(ethylene terephthalate)(PET), and various esters, it is reasonable to infer that it can also act on *p*-nitrophenyl butyrate. The hydrolysis of such esters would involve the addition of water to break the ester bond, leading to the formation of an alcohol and a carboxylic acid derivative. This function aligns with the protein’s classification within the AB hydrolase superfamily and its annotation as a cutinase and poly(ethylene terephthalate) hydrolase, suggesting a broad spectrum of ester hydrolysis capabilities. |

**Supplementary Table S16:**
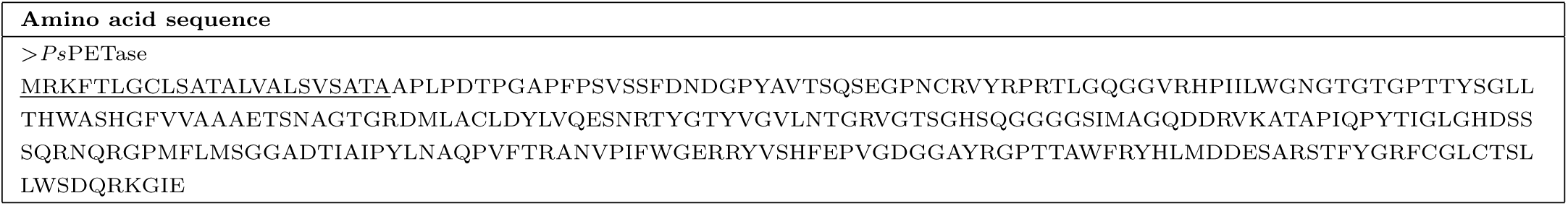
*Ps*PETase protein sequence. The underlined segment denotes the N-terminal signal peptide as predicted by SignalP 5.0.

**Extended Figure S1.**
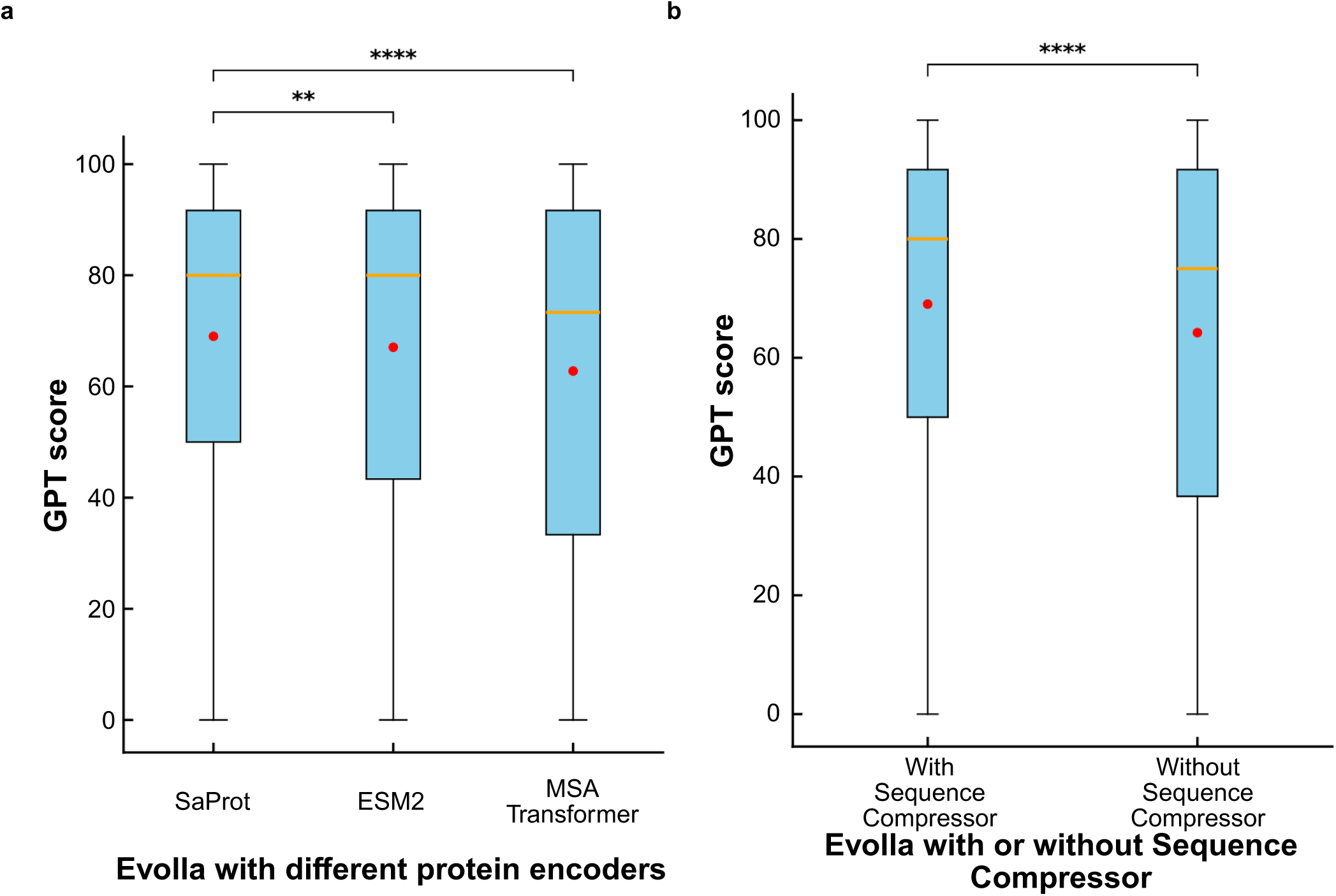
Ablation studies for the modules of Evolla. **(a)** Comparison of Evolla’s performance using three different protein encoders: SaProt, ESM-2, and MSA Transformer. To ensure a fair comparison, all models were trained on the same set of question-answer (QA) pairs, with the only difference being the protein representation generated by each respective encoder. **(b)** Ablation study on the Sequence Compressor module. The performance of the standard Evolla model (“With Sequence Compressor”) is compared against a variant where the compressor module is replaced by a single linear layer (“Without Sequence Compressor”). For all box plots, the center line (orange) represents the median, the dot (red) indicates the mean, the box limits correspond to the 25th and 75th percentiles, and the whiskers extend from the box to the farthest data point lying within 1.5x the inter-quartile range (IQR) from the box. Statistical significance between groups was determined using a two-sided Mann-Whitney U test. Asterisks denote significance levels: **p *<* 0.01, ****p *<* 0.0001. The analysis was performed on a test set of n = 5,968 records.

**Extended Figure S2.**
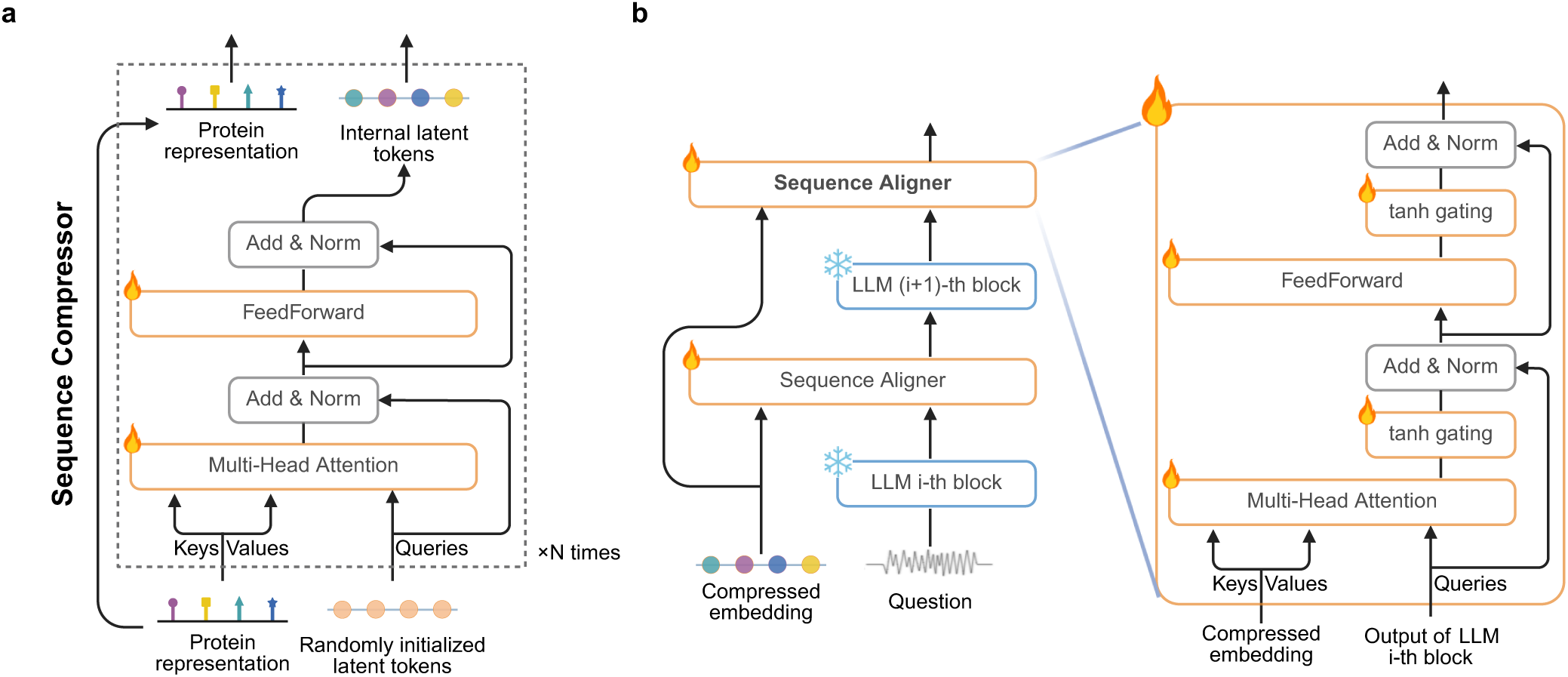
Detailed architecture of the interfacing module. **(a)** The architecture of Sequence Compressor. This module distills a variable-length protein representation into a fixed-length protein codebook. A set of randomly initialized, learnable latent tokens act as Queries that attend to the protein representation (Keys, Values) through a stack of N cross-attention blocks. This process iteratively refines the latent tokens into a dense representation of protein function. **(b)** The architecture of Sequence Aligner. The Sequence Aligner module is inserted between the frozen layers of a large language model (LLM). It allows the LLM’s internal hidden states (Queries) to attend to the protein codebook (Keys, Values) generated by the Sequence Compressor. As shown in the magnified view, a learnable tanh gating mechanism is applied to the outputs of both the multi-head cross-attention and the feed-forward sub-layers. This gate modulates the strength of the protein-specific information before it is integrated back into the LLM’s main pathway via a residual connection (Add).

**Extended Figure S3.**
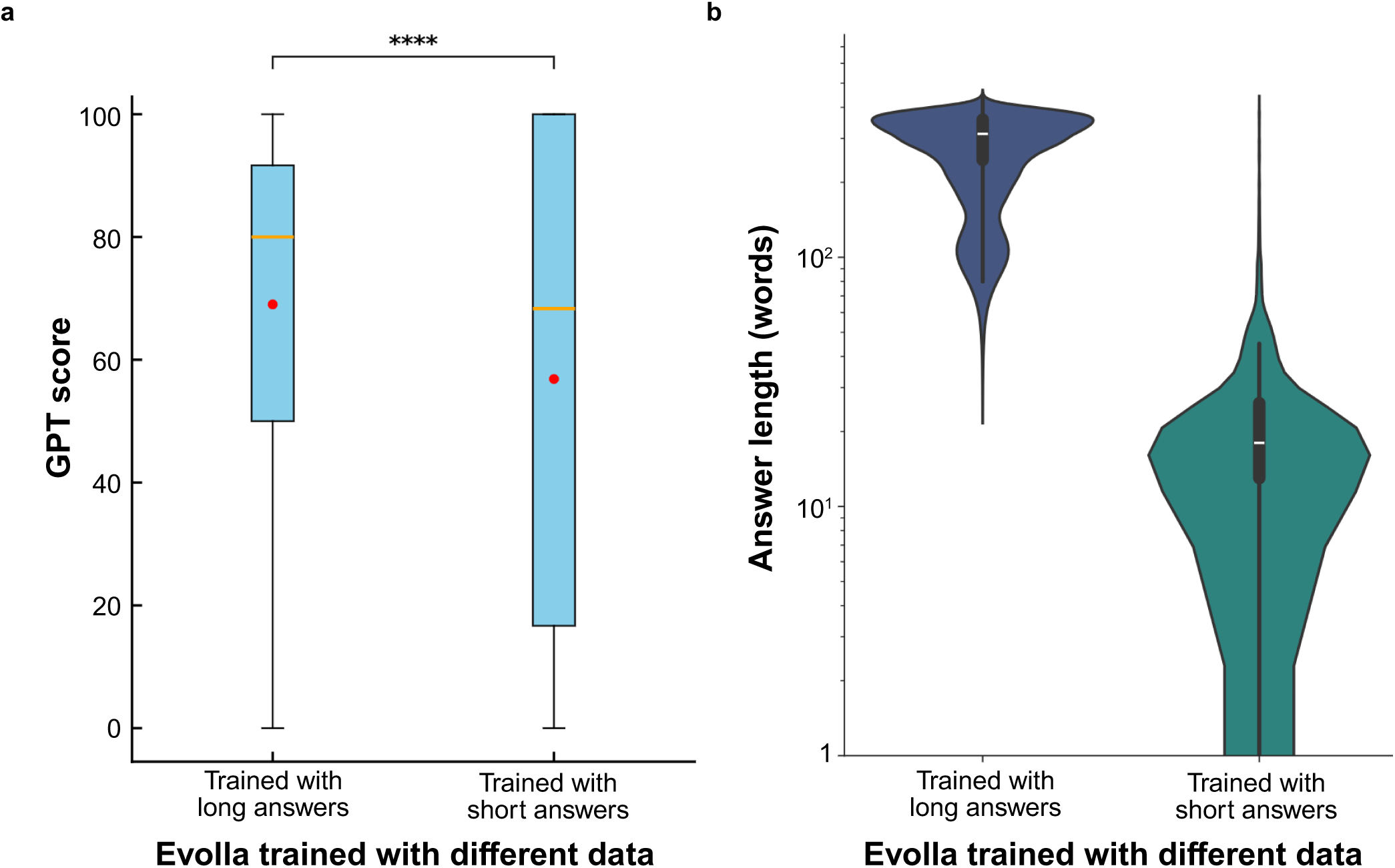
The impact of answer format in training data on Evolla’s performance and output. **(a)** Comparison of model performance evaluated by GPT score. Two Evolla models were fine-tuned using 1 million triplets with identical input proteins and questions but with distinct types of target answers. One model was trained using comprehensive long-form answers, while the other was trained using original short-form data text extracted from databases. Notably while the model trained with short answers achieves a higher upper quartile due to the fact that short sentences (in the ground truth) are easier to remember, it exhibits significantly higher variance and a lower median compared to the long-answer model. In the box plot, the center line (orange) represents the median, the dot (red) indicates the mean, the box limits correspond to the 25th and 75th percentiles, and the whiskers extend from the box to the farthest data point lying within 1.5*×* the inter-quartile range (IQR) from the box. Statistical significance between the two groups was determined using a two-sided Mann-Whitney U test (****p *<* 0.0001). **(b)** Analysis of the generated answer length. The violin plots demonstrate that the output length distribution for each model mirrors its respective training data. Regarding the embedded box plots, the central line denotes the median, the upper and lower bounds of the box represent the 75th and 25th percentiles respectively, and the whiskers extend to the minimum and maximum values. This analysis was conducted on a test set comprising n = 5,968 records.

**Extended Figure S4.**
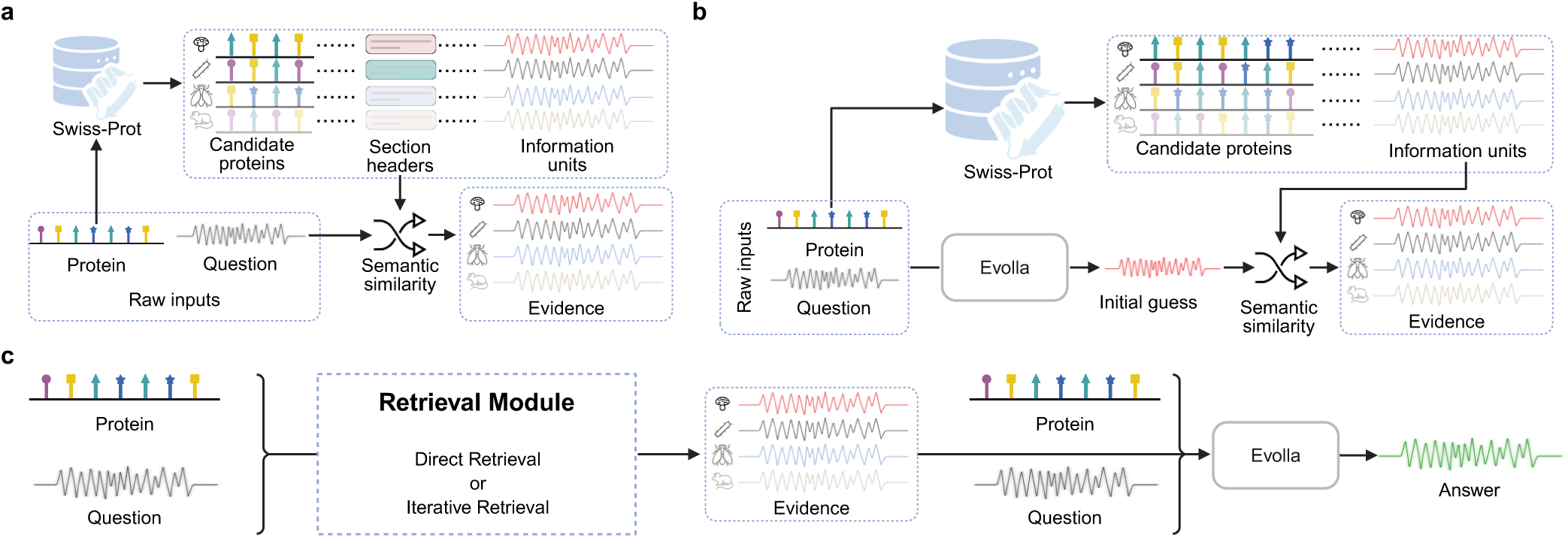
Schematic of the Retrieval-Augmented Generation (RAG) framework. **(a)** The direct retrieval pipeline. Candidate proteins are identified from the Swiss-Prot database based on the input protein. Subsequently, specific information units within these candidates are retrieved by calculating the semantic similarity between the user question and section headers. The corresponding texts are provided as evidence. **(b)** The iterative retrieval pipeline. Adopting a generate-then-retrieve paradigm, the model first produces an initial guess based on the raw inputs. This preliminary response is then used to query the full textual content of the related proteins to find relevant evidence. **(c)** The answer generation pipeline with RAG. The evidence gathered from either the direct or iterative pipeline is integrated with the original protein representation and user question to guide the Evolla model in generating the final response.

**Extended Figure S5.**
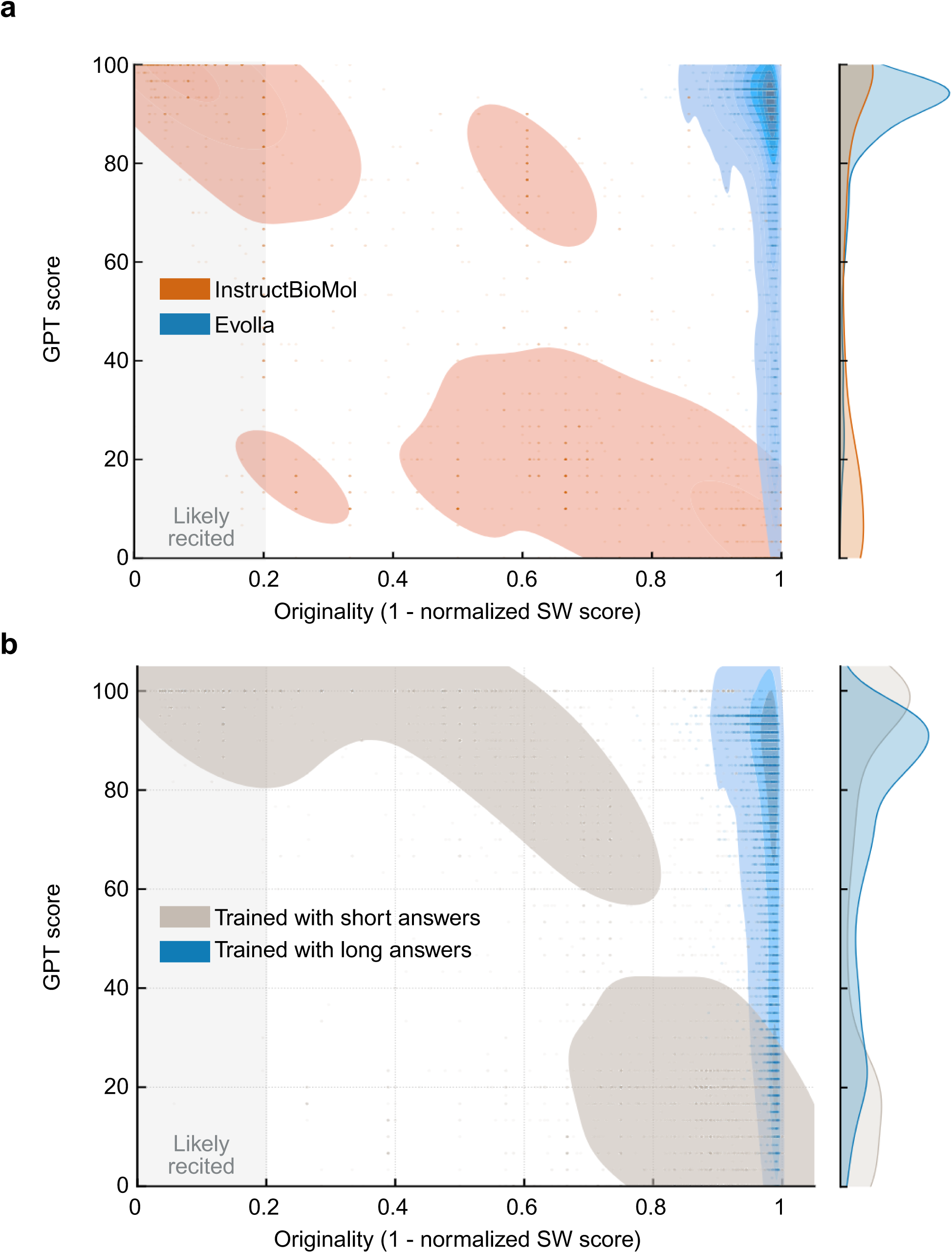
Assessment of originality and factual accuracy in generated content. The scatter plots illustrate the relationship between factual accuracy (GPT score, y-axis) and content originality (1 *−* normalized SW score, x-axis). The shaded regions represent the kernel density estimation of the data distribution, and the curves on the right show the marginal density of GPT scores. A “Likely recited” zone (gray box, Originality *<* 0.2) marks outputs that are verbatim or near-verbatim reproductions of the training data.**(a)** Performance comparison between Evolla (blue) and InstructBioMol (orange). **(b)** Impact of training data length. The plot compares Evolla trained with long answers (blue) against a version trained with short answers (gray). The latter exhibits a bifurcated distribution, tending to either memorize training data (low originality) or generate low-accuracy content. Analysis was performed on a test set of *n* = 5, 968 records.

**Extended Figure S6.**
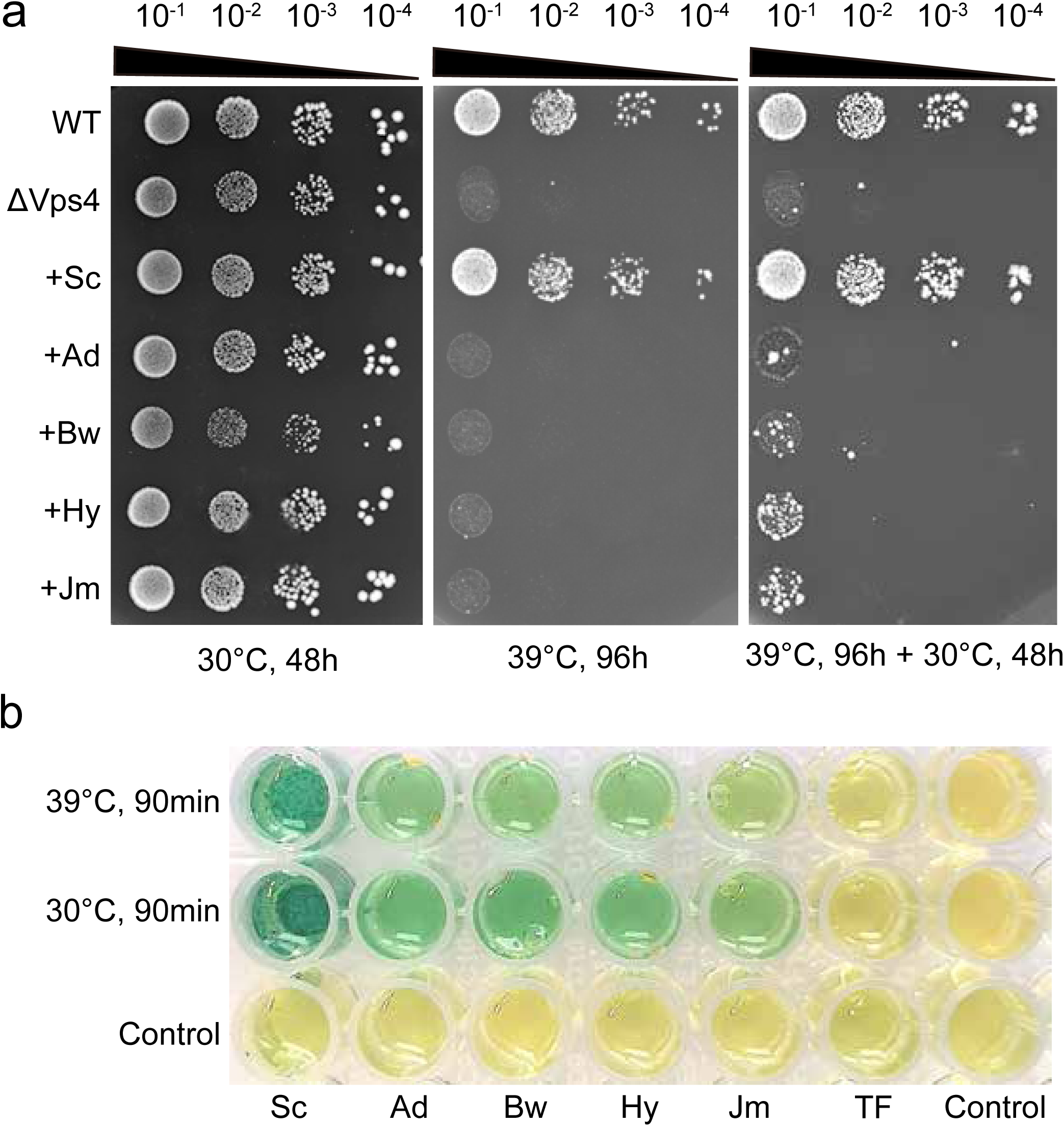
Functional characterization of Asgard Vps4 homologs in complementing a *Saccharomyces cerevisiae vps4* null mutant. **(a)** Complementation of temperature-sensitive growth. Serial tenfold dilutions of yeast cultures (initial OD_600_ = 0.1; 5 *µ*L) were spotted onto SD-Ura plates and incubated at the indicated temperatures. After 96 h at 39 °C, normal growth was observed only in the wild-type (WT) strain and the Δ*vps4* mutant expressing native *S. cerevisiae* Vps4 (+Sc). Expression of Asgard Vps4 homologs-including *Atabeyarchaeum deiterrae* (+Ad), *Borrarchaeum weybense* (+Bw), *Hermodarchaeum yapensis* (+Hy), *Jordiarchaeum madagascariense* (Jm)-slightly suppressed the growth defect at 39 °C, followed by partial recovery at 30 °C. **(b)** ATPase activity assays. ATP hydrolysis by purified Asgard Vps4 at 30 °C and 39 °C, indicated by a gold-to-green color shift (no enzyme as negative control), confirming their enzymatic function and supporting their role in rescuing the Δ*vps4* strain under restrictive conditions.

**Extended Figure S7.**
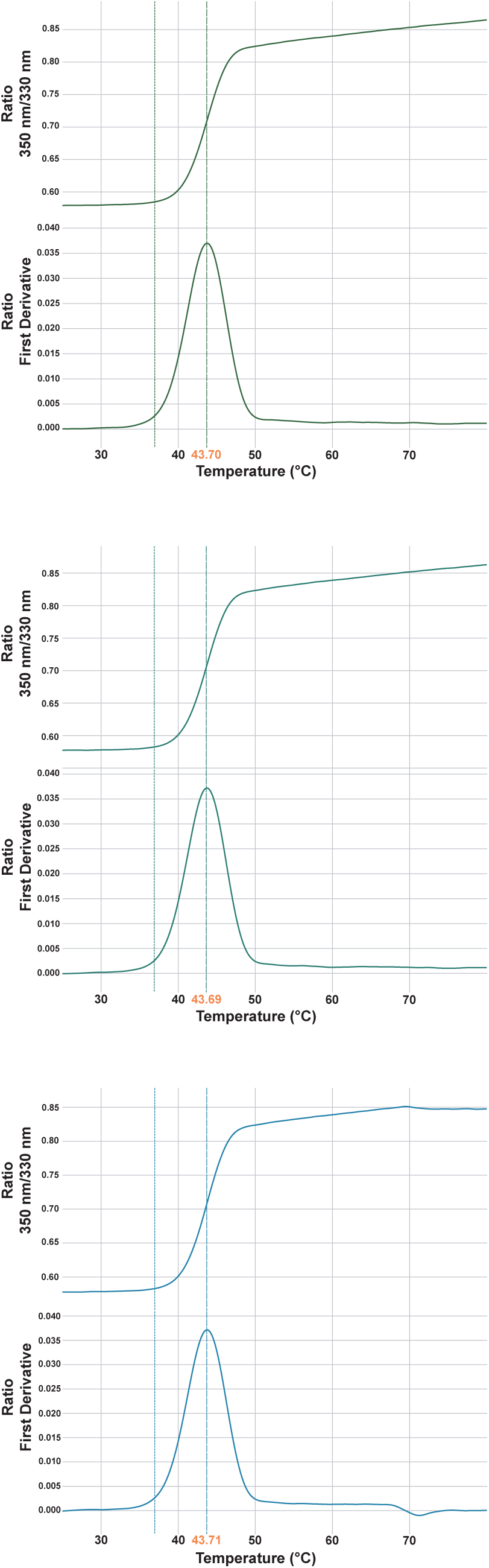
Thermal denaturation profiles of *Ps*PETase determined by nano-differential scanning fluorimetry (nanoDSF). Thermal stability analysis of purified *Ps*PETase (1000 nM) was performed in 50 mM KH_2_PO_4_-NaOH buffer (pH 8.0). The panels show the ratio of intrinsic tryptophan/tyrosine fluorescence at 350 nm and 330 nm (*F*_350_*/F*_330_, upper curves) and its corresponding first derivative (lower curves) as a function of temperature. The melting temperature (*Tm*) is identified as the peak of the first derivative of the fluorescence ratio. Three independent experimental replicates (*n* = 3) are shown, yielding *Tm* values of 43.70 °C, 43.69 °C, and 43.71 °C, respectively, resulting in a mean *Tm* of 43.70 °C. The temperature was increased from 25 °C to 80 °C at a linear ramp rate of 1 °C min*^−^*^1^.

### A.1 Validation of GPT score Reliability and Alignment with Human Judgment

To ensure the robustness of the GPT score as an evaluation metric, we conducted an assessment focusing on two key dimensions: test-retest reliability and alignment with human expert judgment.

- **Test-Retest Reliability and Stability.** We evaluated the stability of the GPT score by performing a test-retest analysis on the test set generated by the Evolla-10B model (the DPO with iterative RAG version). The entire dataset was scored twice independently using the GPT score pipeline, yielding highly consistent global mean scores (88.14 vs. 88.04).

We first calculated the Spearman’s rank correlation coefficient, which yielded a strong positive correlation (*ρ* = 0.8586, Extended Fig. S8c). Further, we computed the Intraclass Correlation Coefficient (ICC) [99]. Unlike simple correlation which ignores systematic shifts, ICC assesses reliability by comparing the variability between different samples to the total variability, ensuring that score differences reflect actual quality distinctions rather than measurement error or systematic bias.

Specifically, we reported the ICC(2,1) index (two-way random-effects model, single measures, absolute agreement) calculated using the Python pingouin library. The analysis revealed an excellent degree of reliability, with an ICC value of 0.97 (95% CI: [0.97, 0.97]; *p <* 0.001). These results demonstrate that despite the inherent stochastic nature of LLMs, the GPT score provides highly consistent and reproducible evaluations.

- **Alignment with Human Expert Evaluation.** We conducted a comparative study to verify whether the semantic accuracy of GPT scores is consistent with the assessment results of human experts.

We stratified the generated responses into five quality intervals based on their GPT scores (0–20, 20–40, 40–60, 60–80, and 80–100) and randomly sampled 20 responses from each interval, resulting in a diverse dataset of 100 responses covering the full spectrum of response quality. Three domain experts were invited to evaluate these responses, assigning scores from 0 to 100 based on the relevance, factual consistency, and the comprehensive coverage of key functional items relative to the provided ground-truth Information Units (IUs).

The alignment between human expert scores and GPT scores was evaluated using Spearman’s rank correlation and the ICC. First, we examined the correlation at the individual level. The GPT score exhibited strong positive correlations with each of the three experts (*ρ* = 0.9198, 0.8942, and 0.9553, respectively). Notably, when compared against the ensemble average of the human experts, the correlation further improved to *ρ* = 0.9594 (Extended Fig. S8d). Furthermore, to assess the absolute agreement between the GPT score and the mean human score, we calculated the ICC(2,1) index. The results demonstrated an strong degree of agreement (ICC = 0.95, 95% CI: [0.89, 0.97]; *p <* 0.001), confirming that the GPT score serves as a highly reliable proxy for human expert judgment in assessing the biological validity of generated narratives.

**Extended Figure S8.**
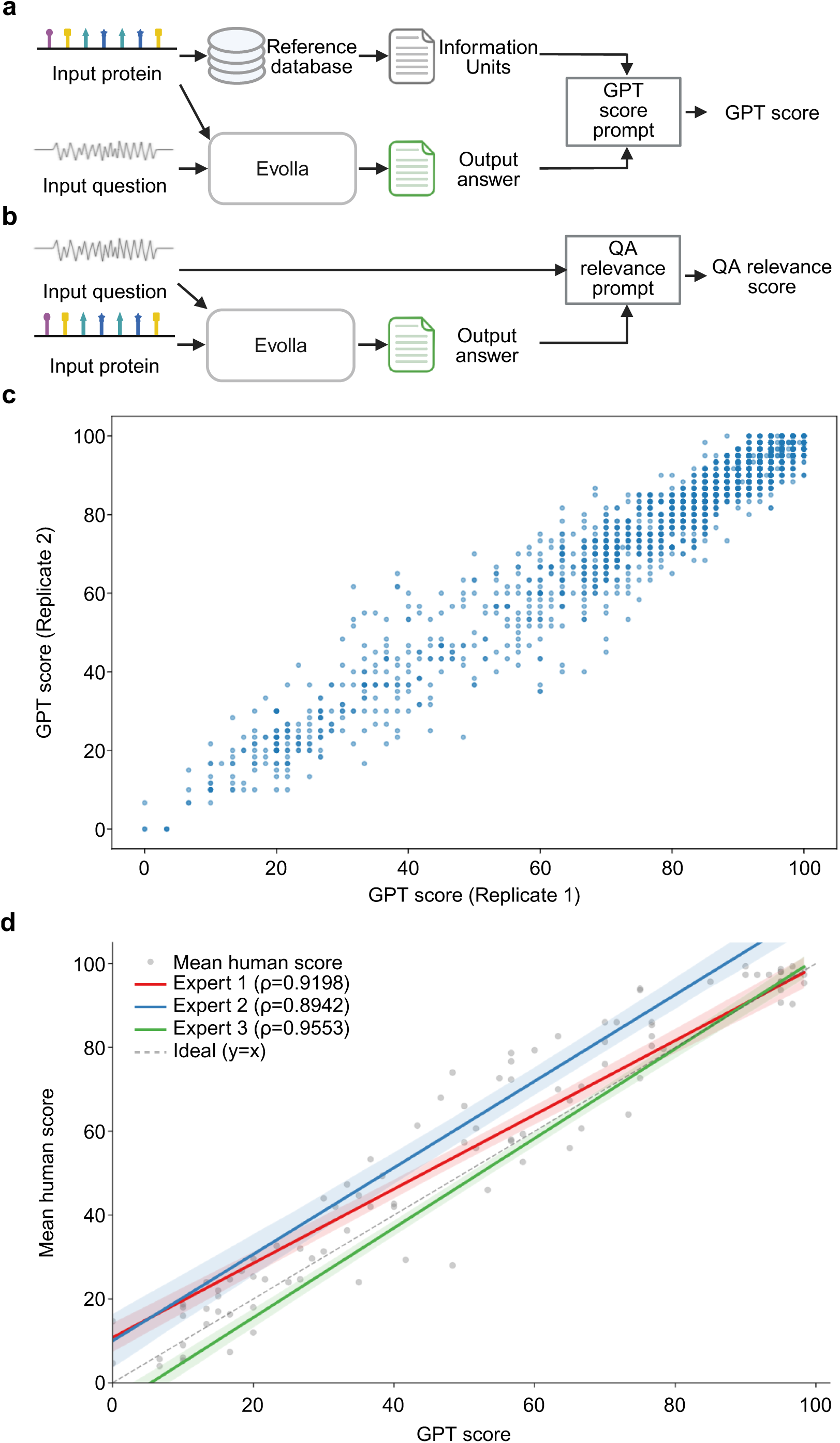
Validation of LLM-based evaluation metrics for protein question answering. **(a)** Schematic pipeline for calculating the GPT score. **(b)** Schematic pipeline for calculating the QA relevance score. **(c)** Test-retest reliability assessment. The scatter plot displays the correlation between two independent GPT score evaluations performed on the full test set (*n* = 5, 968), yielding a Spearman’s rank correlation coefficient (*ρ*) of 0.8586 and an Intraclass Correlation Coefficient (ICC) of 0.97 (95% CI: [0.97, 0.97]; *p <* 0.001). **(d)** Alignment with human expert evaluation. The plot compares automated GPT scores against ratings from three domain experts on a sample of *n* = 100 responses. Grey points represent the mean score across the three experts, which exhibits a high correlation with the GPT score (Spearman’s *ρ* = 0.9594). The solid colored lines (with shaded 95% confidence intervals) depict the linear regression fits for each individual expert, demonstrating strong correlations with the GPT score (Spearman’s *ρ* = 0.9198, 0.8942, and 0.9553 for Experts 1, 2, and 3, respectively). The dashed grey line represents the line of perfect agreement (*y* = *x*).

**Supplementary Table S17:** Three representative prompts for general LLMs used to answer specific questions based on protein information.

|  |
| --- |
| As an advanced language model with extensive training and knowledge, I would like you to perform a comprehensive analysis of the following amino acid sequence. Please consider the following aspects in your analysis:<br>- Protein Structure Prediction: Provide insights into the possible secondary and tertiary structures that this amino acid sequence might form.<br>- Functional Domains: Identify any known functional domains or motifs within the sequence that could indicate the protein's function.<br>- Post-Translational Modifications: Highlight potential sites for post-translational modifications and discuss their possible effects on protein activity.<br>- Biological Pathways: Explain how this protein might interact within specific biological pathways or processes.<br>- Disease Associations: Discuss any known associations between this amino acid sequence or similar sequences and particular diseases or medical conditions.<br>- Evolutionary Conservation: Analyze the evolutionary conservation of this sequence across different species to infer its importance.<br>- Potential Interactions: Suggest possible interactions with other proteins or molecules based on the sequence characteristics.<br>Please ensure that your analysis is detailed and supported by relevant biological concepts and knowledge.<br>Additional Requirement: After the summary of your analysis, please answer the specific question I will provide.<br>**\*\*Amino Acid Sequence:\*\***<br>{sequence}<br>**\*\*Question:\*\***<br>{question} |
| As an advanced language model with extensive training and knowledge, please perform a thorough analysis of the following amino acid sequence. Your analysis should cover the following aspects:<br>- Protein Structure Prediction: Discuss the potential secondary and tertiary structures that this amino acid sequence may form.<br>- Functional Domains: Identify any known functional domains or motifs within the sequence that could suggest the protein's function.<br>- Post-Translational Modifications: Highlight possible sites for post-translational modifications and explain their potential impact on protein activity.<br>- Biological Pathways: Describe how this protein might interact within specific biological pathways or processes.<br>- Disease Associations: Explore any known links between this amino acid sequence or similar sequences and particular diseases or medical conditions.<br>- Evolutionary Conservation: Assess the evolutionary conservation of this sequence across various species to determine its significance.<br>- Potential Interactions: Suggest possible interactions with other proteins or molecules based on the sequence characteristics.<br>Additional Requirement: After summarizing your analysis, please answer the specific question I will provide.<br>**\*\*Amino Acid Sequence:\*\***<br>{sequence}<br>**\*\*Question:\*\***<br>{question} |
| Please act as a highly trained language model with extensive knowledge and conduct a detailed examination of the following amino acid sequence. Address the following components in your analysis:<br>- Protein Structure Prediction: Provide insights into the possible secondary and tertiary structures formed by this amino acid sequence.<br>- Functional Domains: Detect any known functional domains or motifs within the sequence that may indicate the protein's role.<br>- Post-Translational Modifications: Identify potential sites for post-translational modifications and discuss how they might affect protein function.<br>- Biological Pathways: Explain the potential interactions of this protein within specific biological pathways or processes.<br>- Disease Associations: Discuss any associations between this amino acid sequence or similar sequences and particular diseases or health conditions.<br>- Evolutionary Conservation: Evaluate the evolutionary conservation of this sequence across different species to infer its importance.<br>- Potential Interactions: Recommend possible interactions with other proteins or molecules based on the sequence features.<br>Extra Requirement: Following your analysis summary, please respond to a specific question I will pose.<br>**\*\*Amino Acid Sequence:\*\***<br>{sequence}<br>**\*\*Question:\*\***<br>{question} |

**Supplementary Table S18:** An example of RAG retrieval results for the target protein B8CH76. The prompt integrates relevant functional metadata from three retrieved proteins (A8FP54, B0TLA9, and A8GYE8) to provide context for the specific query regarding enzyme catalysis.

|  |
| --- |
| **Similar proteins:**<br>**Entry: A8FP54**<br>**GO annotation:** On the topic of molecular function, the functional GO definition for this protein spans glycine-tRNA ligase activity.<br>**GO annotation:** The GO term regarding biological process encompassing glycyl-tRNA aminoacylation covers this protein.<br>**Catalytic activity:** The protein enables the chemical conversion seen in the reaction: ATP + glycine + tRNA(Gly) = AMP + diphosphate + glycyl-tRNA(Gly).<br>**Protein names:** Commonly, this protein is recognized by the epithet GlyRS.<br>**Entry: B0TLA9**<br>**GO annotation:** The GO term enveloping glycine-tRNA ligase activity involves molecular function for this protein.<br>**GO annotation:** The GO term regarding biological process involving glycyl-tRNA aminoacylation envelopes this protein.<br>**Catalytic activity:** This protein's enzymatic function contributes to catalyzing the reaction: ATP + glycine + tRNA(Gly) = AMP + diphosphate + glycyl-tRNA(Gly).<br>**Protein names:** GlyRS is how this protein is identified in the scientific lexicon.<br>**Entry: A8GYE8**<br>**GO annotation:** When pertaining to molecular function, the GO designation for this protein entails glycine-tRNA ligase activity.<br>**GO annotation:** Concerning biological process, the GO term regarding glycyl-tRNA aminoacylation involves this protein.<br>**Catalytic activity:** This protein has the primary purpose of catalyzing the reaction: ATP + glycine + tRNA(Gly) = AMP + diphosphate + glycyl-tRNA(Gly).<br>**Protein names:** GlyRS is how this protein is identified in the scientific lexicon.<br>**Question:** Could you clarify the particular process involving enzyme catalysis that is expedited by this protein? |

## Footnotes

1 https://github.com/HICAI-ZJU/InstructBioMol

